# IRF8-mutant B cell lymphoma evades immunity through a CD74-dependent deregulation of antigen processing and presentation in MHC CII complexes

**DOI:** 10.1101/2023.10.14.560755

**Authors:** Zhijun Qiu, Jihane Khalife, An-Ping Lin, Purushoth Ethiraj, Carine Jaafar, Lilly Chiou, Gabriela Huelgas-Morales, Sadia Aslam, Shailee Arya, Yogesh K. Gupta, Patricia L. M. Dahia, Ricardo C.T. Aguiar

**Affiliations:** Division of Hematology and Medical Oncology, Mays Cancer Center, University of Texas Health Science Center San Antonio, San Antonio, Texas, 78229, USA; Department of Biochemistry and Structural Biology, University of Texas Health Science Center San Antonio, San Antonio, Texas, 78229, USA; South Texas Veterans Health Care System, Audie Murphy VA Hospital, San Antonio, Texas, 78229, USA

## Abstract

In diffuse large B-cell lymphoma (DLBCL), the transcription factor IRF8 is the target of a series of potentially oncogenic events, including, chromosomal translocation, focal amplification, and super-enhancer perturbations. IRF8 is also frequently mutant in DLBCL, but how these variants contribute to lymphomagenesis is unknown. We modeled IRF8 mutations in DLBCL and found that they did not meaningfully impact cell fitness. Instead, IRF8 mutants, mapping either to the DNA-binding domain (DBD) or c-terminal tail, displayed diminished transcription activity towards CIITA, a direct IRF8 target. In primary DLBCL, IRF8 mutations were mutually exclusive with mutations in genes involved in antigen presentation. Concordantly, expression of IRF8 mutants in murine B cell lymphomas uniformly suppressed CD4, but not CD8, activation elicited by antigen presentation. Unexpectedly, IRF8 mutation did not modify MHC CII expression on the cell surface, rather it downmodulated CD74 and HLA-DM, intracellular regulators of antigen peptide processing/loading in the MHC CII complex. These changes were functionally relevant as, in comparison to IRF8 WT, mice harboring IRF8 mutant lymphomas displayed a significantly higher tumor burden, in association with a substantial remodeling of the tumor microenvironment (TME), typified by depletion of CD4, CD8, Th1 and NK cells, and increase in T-regs and Tfh cells. Importantly, the clinical and immune phenotypes of IRF8-mutant lymphomas were rescued in vivo by ectopic expression of CD74. Deconvolution of bulk RNAseq data from primary human DLBCL recapitulated part of the immune remodeling detected in mice and pointed to depletion of dendritic cells as another feature of IRF8 mutant TME. We concluded that IRF8 mutations contribute to DLBCL biology by facilitating immune escape.

## Introduction

The immune system can detect early signs of, and eliminate, malignant cell transformation. To thrive, cancer cells acquire genetic changes that can disengage this surveillance system^1,2^. Pharmacological strategies that reestablish anti-tumor immunity have all but cemented the pivotal role of the immune system in cancer biology^3^.

In diffuse large B-cell lymphoma (DLBCL), a common, genetically diverse, and still often fatal B cell malignancy, the mutations that disarm the immune system range from highly predictable suspects, genes encoding products directly involved in antigen processing/presentation (e.g., *B2M, CIITA, HLA-A/B/C*)^4-6^, to less immediately obvious candidates, such as chromatin modifiers (e.g., *CREBBP and EZH2*)^7-9^.

The genetic heterogeneity of DLBCL is well recognized and more than 100 protein-coding genes are recurrently mutated or targeted by somatic structural changes in this B cell lymphoma^10,11^. The most current classification systems identify approximately six DLBCL subgroups driven by distinct biological processes^12,13^. In multiple instances, the mechanism by which a given driver gene contributes to DLBCL development has been, at least partially, elucidated^14^. However, in the case of interferon regulatory factor 8 (IRF8), which is a common target for somatic mutations in DLBCL, it remains unclear how these variants may contribute to lymphomagenesis. Notably, IRF8 is also deregulated by series of potentially oncogenic events, including chromosomal translocation^15^, focal amplification^12,13^, and super-enhancer perturbations^16,17^, in principle suggesting that the mutations may also represented gain-of-function events.

IRF8, a member of the IRF family of transcription factor, plays important roles in both myeloid differentiation and B cell development^18,19^. IRF8 binds to specific DNA sequences as a heterodimer with partner transcription factors and can act either as an activator or a repressor. In mice, germline deletion of *Irf8* yields a dominant myeloid phenotype (a chronic myeloid leukemia-like disease), which agrees with Irf8’s essential role on monocyte and dendritic cell (DC) development ^20^. Concordantly, in humans, germline loss-of-function *IRF8* missense mutations, cause rare, at times severe, inborn errors of immunity (IEI), characterized by susceptibility to infections, depletion of dendritic cells (DC) and reduced numbers and/or impaired activity of NK cells^21-24^.

IRF8’s contribution to B cell physiology is more nuanced. IRF8 is expressed in multiple B cell developmental stages, predominates in the germinal center and it is absent in plasma cells^18,25^. IRF8 regulates several genes important for normal and malignant B cell biology (e.g., *BCL6* and *AICDA*)^26^, but deletion of *Irf8* in the mouse B cell compartment yielded only a modest phenotype^27^, which could be better appreciated in mice with B cell specific double *Irf8*/*Spi1* (encoding PU.1) knockout. In these instances, Irf8 was shown to be important for the development of follicular B cells, germinal center responses and to constrain plasma cell differentiation^28,29^. However, many of the validated IRF8 targets are actually known to be important for antigen processing/presentation^30-33^, raising the possibility that IRF8 may also play a role in the immune composition of the tumor microenvironment, while its dysfunction could facilitate immune escape.

Here, we show that in DLBCL somatic *IRF8* and *CIITA* mutations are mutually exclusive, and that IRF8 mutants, mapping either to the n-terminal DNA-binding domain (DBD) or c-terminal tail, displayed diminished transcription activity towards CIITA, a direct IRF8 target and master regulator of *de novo* transcription of MHC CII complex genes^33-35^. Further, we show that while MHC CII expression on the cell surface was not meaningfully decreased in IRF8-mutant lymphoma models, these tumors displayed significant downmodulation of CD74 and HLA-DM, key regulators of antigen peptide processing/loading in the MHC CII complex^34^. More importantly, mice harboring IRF8 mutant lymphomas displayed higher tumor burden, in association with remodeling of the tumor microenvironment (TME), an immunophenotype which was rescued in vivo by ectopic expression of CD74 and validated in primary human DLBCLs. We concluded that IRF8 mutations are hypomorphic and contribute to DLBCL biology by facilitating immune evasion. Together with the earlier evidence of oncogenic aberrations targeting its locus^15-17^, we propose that IRF8’s role in DLBCL is uniquely multilayered and that both its loss and gain of function may be lymphomagenic.

## Results

### Landscape of IRF8 mutation in DLBCL

Approximately 8% of DLBCLs harbor IRF8 mutation (n=244 from 3187 previously reported DLBCL sequences, Supplemental Table 1)^7,12,13,36-41^. The mutations cluster into two recognized IRF8 domains, DNA-binding, and protein interaction domains (IRF association domain, IAD), but also in a less well-defined c-terminal region of the protein. Missense variants predominate in the DNA- binding and IAD, whereas most of the mutation in the c-terminal tail are truncating (Figures 1A and B, Supplemental Table 2). In all instances in which variant allele frequency (VAF) could be defined, the IRF8 mutations were found to be monoallelic (Supplemental Table 3). Most of the IRF8-mutant DLBCLs are of the GCB-like subtype (COO classifier) and EZB category (Figure 1C, Supplemental Table 4). IRF8 is also frequently mutant in follicular lymphoma^42,43^, Burkitt lymphoma^44,45^ and to lesser extent in marginal zone lymphoma^46^ (Supplemental Figure 1A, Supplemental Table 5), all with a mutation pattern similar to that found in DLBCL (Supplemental Table 6).

**Figure 1.**
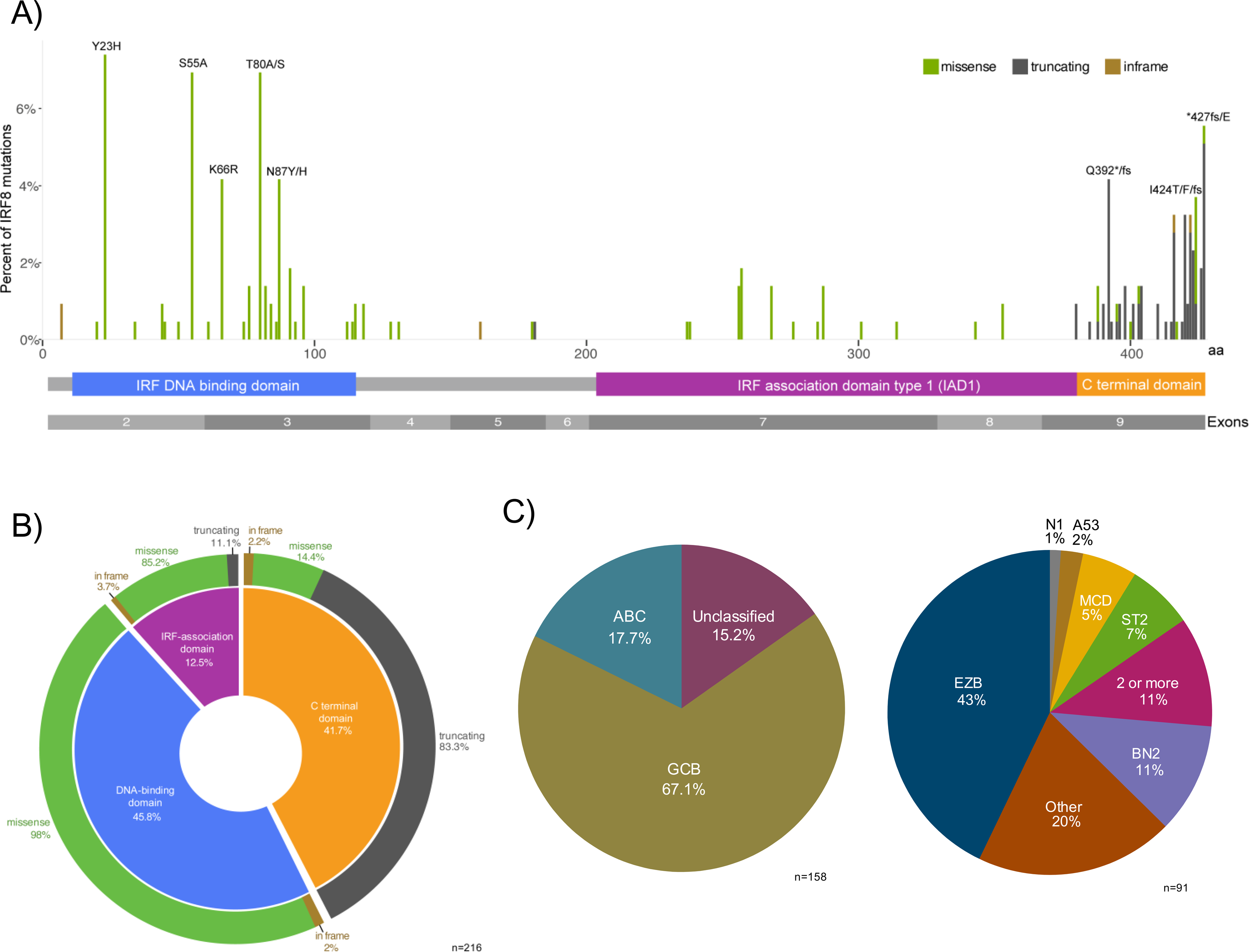
Distribution of *IRF8* variants in DLBCL. (A) Diagram of 216 *IRF8* gene variants reported in publicly available DLBCL cohorts distributed along the linear amino acid sequence of the protein and its recognized domains, DNA binding domain, IRF-Associated Domain and c-terminal domain; the frequency of occurrence (%) of the variants is indicated by the height of the line, and the type of mutation is color coded based on the mutation class (missense, truncating or in-frame), related to Supplemental Tables 1-3 (diagram excludes samples for which variant-specific details are not provided); (B) Donut-shape graph showing the distribution (%) of the 216 *IRF8* variants shown in (A) based on the mutation class and functional protein domains, related to Supplemental Table 2. (C) Distribution of *IRF8* variants based on transcription- (left, ABC, GBC or unclassified, n=158) and genetic-based DLBCL subgroups (right, EZB, BN2, A53, MCD, ST2, N1, other or two or more groups, n=91), related to Supplemental Table 4.

### Impact of IRF8 mutations on DLBCL fitness

Earlier, we reported on a small subset of DLBCL with a t(14;16) (q32;q24) juxtaposing the 5’ regulatory regions of the *IRF8* gene to IgH enhancers (Eα and Eµ)^15^. We confirmed that this archetypal gain of function rearrangement resulted in elevated IRF8 expression in primary DLBCL, which in turn conferred a survival benefit to DLBCL cell line models, a finding confirmed by others^47^. A putative oncogenic IRF8 super-enhancer has also been reported^16,17^. Therefore, we first examined if the IRF8 mutations could enhance DLBCL fitness. To generate suitable models for downstream examination of IRF8 mutations, we screened a panel of DLBCL cell lines and found that Toledo did not express IRF8. Therefore, we created Toledo cells expressing IRF8 WT or a series of frequent mutant variants, which map to distinct IRF8 domains (S55A, N87Y, D400G and I424T) (Figure 2A). In addition, we generated IRF8 KO models in two additional DLBCL cell lines, SU-DHL4 and SU-DHL6 (Figure 2A). We confirmed that these DLBCL models express PU.1, an important component of the IRF8 functional complex (Supplemental Figure 2A) ^48^. We next tested if genetic modulation of IRF8 modified DLBCL growth. In comparison to WT IRF8, expression of each IRF8 mutant examined negatively impacted DLBCL growth (Figure 2B). In addition, in agreement with earlier evidence for the growth promoting effect of WT IRF8 overexpression^15^, we found that IRF8 KO limited DLBCL growth (Figure 2B). We concluded that in the DLBCL models tested, IRF8 mutations did not stimulate cell growth, were more akin to KO models, which led us to consider the possibility that these variants may represent a cancer cell intrinsic defect that primarily influences the lymphoma microenvironment.

**Figure 2.**
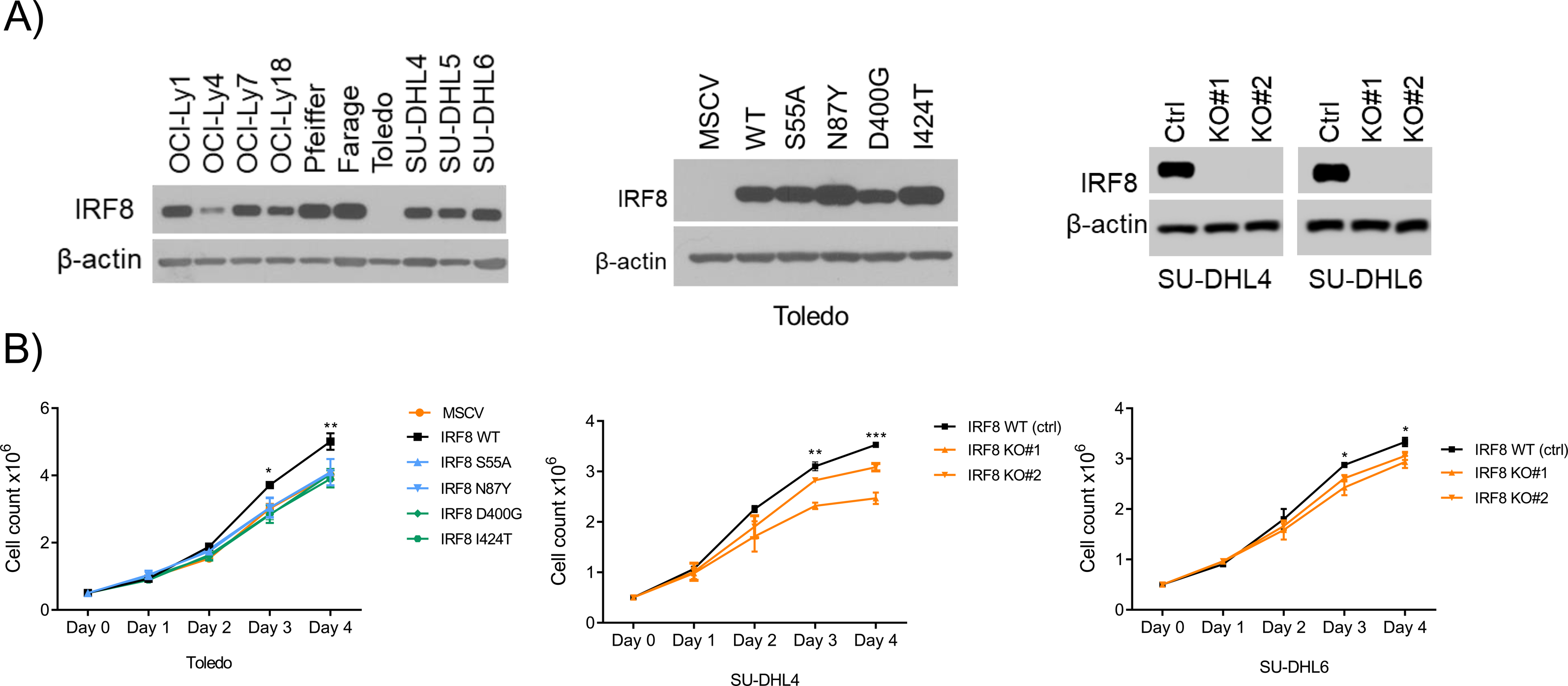
IRF8 influence on DLBCL growth. (A) **left to right**, western blot analyses of IRF8 expression in parental DLBCL cell lines, in the Toledo cell line stably expressing and empty vector (MSCV), IRF8 wild-type or four mutant isoforms (middle panel), and in two DLBCL cell lines with CRISPR-Cas9-based knockout (KO) of IRF8. (B) **left to right**, cell growth pattern, determined with automated fluorescent cell counter, for the cell models described in A. Data are mean ± SD of three biological replicates. P values, WT vs. each mutant or WT vs. KO were calculated with two-sided Student’s t-test *(p<0.05), ** (p<0.01).

### IRF8 mutants have impaired transcriptional activity towards the CIITA promoter

IRF8 regulates a broad array of genes involved in antigen presentation, perhaps chiefly among them CIITA, a transcriptional co-activator and master regulator of MHC CII gene expression^35^, and consequently antigen processing, loading and display on the cell surface^34^. To examine the impact of the IRF8 mutations towards this target, we first created IRF8 KO and “rescue” models (human IRF8 WT, S55A, N87Y, D400G and I424T) in the RAW 264.7 murine macrophage cell line, an interferon gamma (IFN-γ)-responsive model, often used to functionally investigate the IRF proteins^49^ (Figure 3A). These stable cell models were transfected with CIITA WT promoter, or a CIITA construct with point mutations in the Ets-IRF composite element (EICE) site (a gift from Kenneth Wright, Moffit Cancer Center), which has been previously shown to abrogate the recruitment of the IRF8/PU.1 complex to DNA^48^, and luciferase activity measured. We found that the CIITA reporter activity was significantly lower in cells expressing any of the IRF8 mutants in comparison to IRF8 WT (Figure 3B). In agreement with the reporter assay data, *CIITA* expression was significantly higher in IRF8 WT than in IRF8 mutant RAW 264.7 and human DLBCL Toledo cells (Figure 3C, Supplemental Figure 3A). Giving further support to the concept that these four IRF8 mutations may represent “loss-of-function” models, we found that IRF8 KO cells (RAW 264.7, SU-DHL4 and SU-DHL6) displayed lower *CIITA* mRNA levels than their IRF8 WT isogenic controls (Figure 3D). Finally, we examined a large cohort of primary human DLBCL (n=168) and IRF8 and CIITA mutations were found to be mutually exclusive (Figure 3E). In fact, IRF8 mutations in DLBCL were mutually exclusive with mutation in a series of genes that are directly (*CIITA, B2M, HLA-B, HLA-C*) or indirectly (*CREBBP* and *EP300*) involved with antigen presentation and/or remodeling of the TME (Supplemental Figure 3B); mutations in *IRF8* and *HLA-A* or *EZH2*, also has a mutually exclusive tendency, although not meeting statistical significance (Supplemental Figure 3B). We concluded that similarly to CIITA mutations in B cell lymphomas, IRF8 variants may add to the pathogenesis of these tumors by impairing antigen presentation in the context of MHC CII.

**Figure 3.**
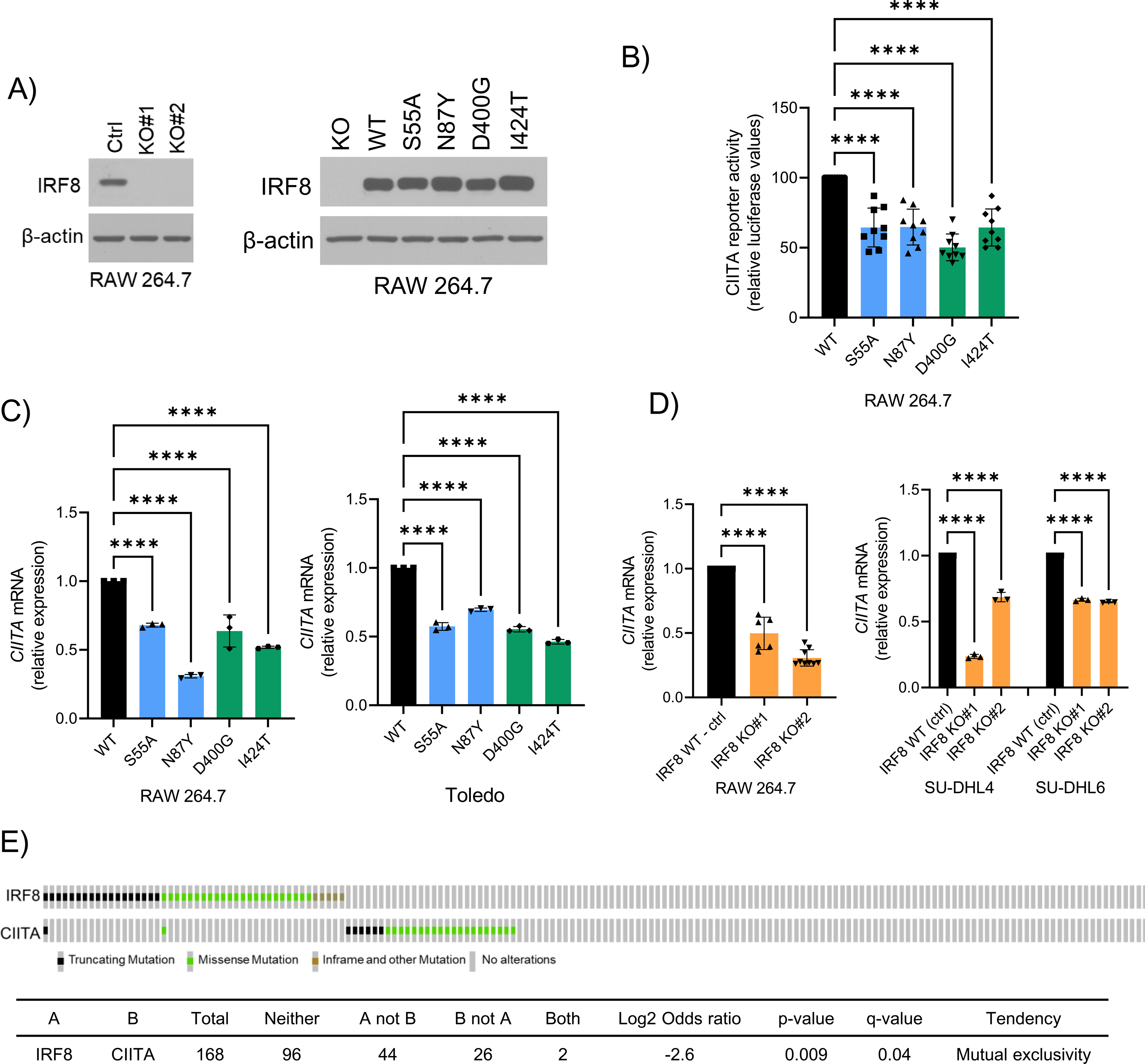
IRF8 modulation of CIITA. (**A**) **left to right**, western blot analyses of IRF8 expression in RAW 264.7 with IRF8 KO or stably expressing IRF8 wild-type and four mutant isoforms. (**B**) Relative luciferase CIITA reporter activity in RAW 264.7 cells expressing IRF8 WT or mutant. Data are mean ± SD of three biological replicates each performed in technical triplicates (all 9 data points shown). Statistical analysis is from one-way ANOVA with Bonferroni post-test, **** (p<0.0001). (**C**) Relative *CIITA* mRNA levels in RAW 264.7 and Toledo models expressing IRF8 WT or mutant. **D)** Relative *CIITA* mRNA levels in RAW 264.7, SU-DHL4 and SU-DHL6 models of IRF8 KO. In C) and D) Data are mean ± SD. Statistical analysis are from one-way ANOVA with Bonferroni post-test, **** (p<0.0001). All assays performed in two to three biological replicates, each with technical triplicates. (**E**) Oncoprint display of comparative distribution of *IRF8* and *CIITA* gene mutations in DLBCLs; total number of samples, mutations distribution, pairwise log2 odds score, and P and Q values of the correlation between the co-occurrence (positive score) or mutually exclusive (negative score) are shown in table format. Symbols are color coded based on the mutation class.

### IRF8 mutation impairs antigen processing/presentation and CD4 activation

To test the idea that IRF8’s role in DLBCL biology may be related to a defective antigen processing/presentation, we first created IRF8 KO models in three murine B cell lymphoma cell lines, A20, 2PK-3 and BCL1 (Figure 4A). Subsequently, we used DO-11.10 murine CD4 cells (a T cell hybridoma specific for chicken ovalbumin, OVA, a gift from Philippa Marrack, National Jewish, Denver, CO, and Jennifer Maynard, UT Austin, Austin, TX) to test the ability of IRF8 WT or KO lymphoma cells to present antigen and, in the context of MHC CII, activate CD4 T cells. IRF8 KO cells were significantly less capable of eliciting a robust CD4 response to OVA presentation, as defined by IL-2 secretion and CD25 cell surface expression in CD4 cells (Figure 4A, Supplemental Figure 4A). These effects were not detected in the absence of OVA (or B lymphoma cells), mitigating concerns about a putative effect of IRF8 KO lymphoma cells on T cells activation outside the antigen presentation axis (Supplemental Figure 4B). These data also confirmed, and expanded on, earlier reports demonstrating the ability of the A20 murine B cell lymphoma cells to effectively present antigen and elicit an immune response ^50,51^. To establish the scope of defect in antigen presentation associated with IRF8 KO, we also tested the MHC CI-CD8 axis. Here, we isolated T cells from the OT-I T cell-receptor (TCR) transgenic mice, which recognizes OVA residues 257-264 in a MHC CI-restricted context^52^, and quantified the resulting CD8 activation/differentiation in vitro by measuring their cytotoxicity towards isogenic IRF8-WT or KO murine B cell lymphoma cells. OVA presentation by the lymphoma cells readily activated the OTI CD8 cells, as measured by secretion of granzyme B and CD8-mediated cytotoxicity (Supplemental Figure 4C). However, contrary to the differential CD4 activation, isogenic IRF8 WT or KO lymphoma cells equally induced CD8-mediated cytotoxicity (Supplemental Figure 4C). Next, we rescued IRF8 expression (WT, S55A, N87Y, D400G and I424T) in the IRF8 KO A20 and 2PK-3 B cell lymphomas models (Figure 4B, Supplemental Figure 4D) and used the DO-11.10 CD4 assay to test the impact of IRF8 mutations on antigen presentation/T-cell activation (here, no analysis of CD8 OTI cells was warranted given that IRF8 KO did not impact CD8 activation). We consistently found that following OVA loading, A20 or 2PK-3 expressing IRF8 WT induced a significantly more robust CD4 activation, measured by IL-2 secretion and CD25 expression, than each of the IRF8 mutations examined (Figure 4B, Supplemental Figure 4E). Of note, IRF8 residues S55, N87, D400 and I424 are fully conserved between human and mouse, increasing confidence on the correlation between output data from either model system. We concluded that in murine models of B cell lymphomas examined in vitro, IRF8 regulates antigen OVA-driven T cell activation in the context of MHC CII but not MHC CI.

**Figure 4.**
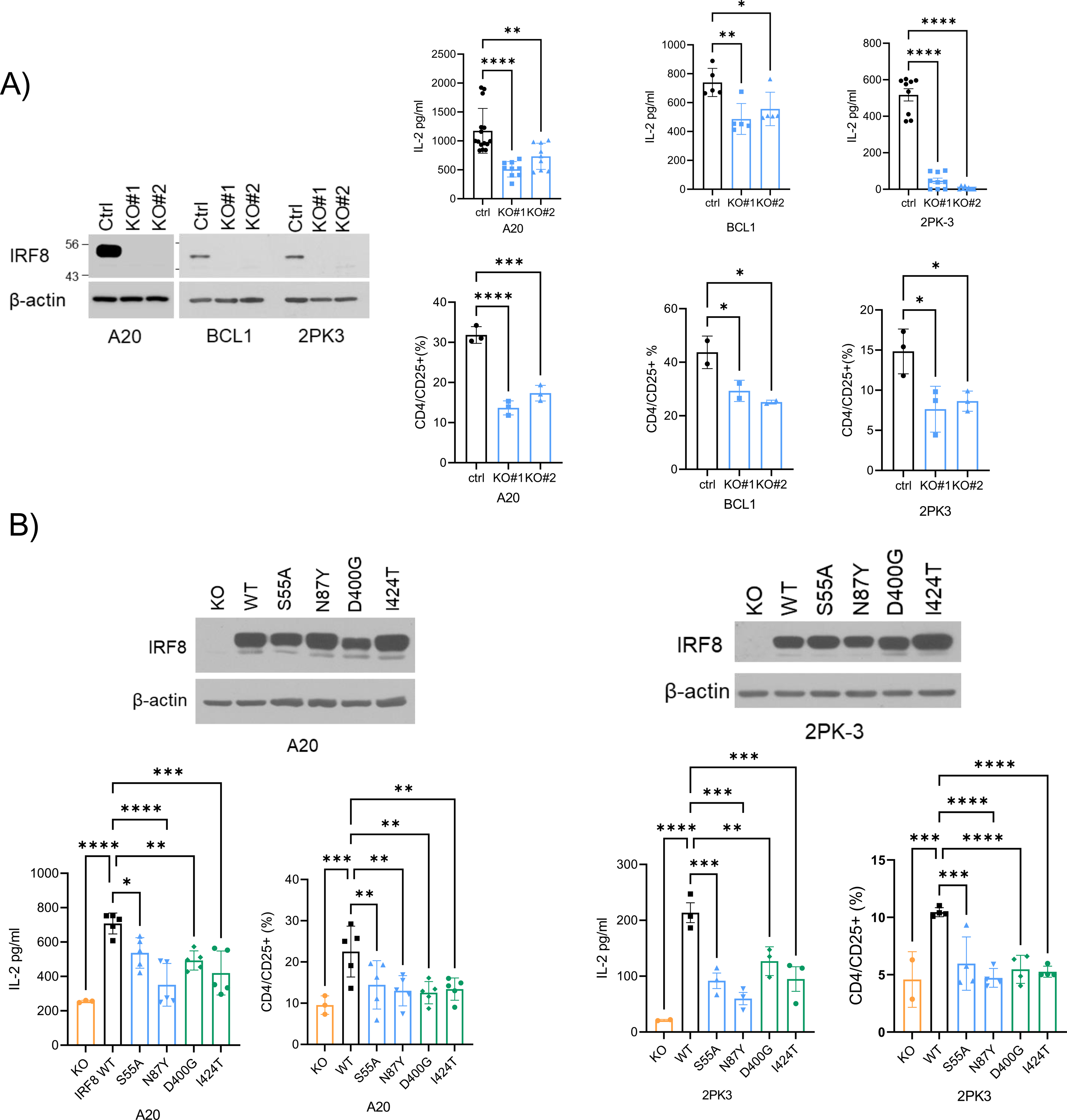
IRF8 impact on antigen driven CD4 activation. (A) **left**, western blot of IRF8 KO models in mouse cell lines A20, BCL1 and 2PK-3 – ctrl is empty vector; **right**, **top and bottom**, respectively, IL-2 levels in the conditioned media and percentage of CD4/CD25+ expression in DO-11.10 cells co-cultured with IRF8 WT or KO antigen presenting cell (APC) models. (B) **top**, western blot of IRF8 A20 (left) or 2PK-3 (right) KO models “rescued” with stable expression of IRF8 wild-type or mutant; **bottom**, IL-2 levels in the conditioned media and percentage of CD4/CD25+ expression in DO-11.10 cells co-cultured with IRF8 KO, WT or mutant APC models. In A) and B) data are mean ± SD of three biological replicates. P values are from one-way ANOVA, Bonferroni or Fisher’s LSD post-test, *(p<0.05), **(p<0.01), ***(p<0.001), **** (p<0.0001).

### Mediators of IRF8 effects on antigen processing/presentation

The defective activation of the CIITA promoter and of CD4 (but not CD8) T-cells by IRF8 mutant B cell lymphomas, together with the well-recognized role of MHC-CII deficiency in DLBCL biology^53^, suggested that IRF8 mutation may deregulate genes and processes that are involved in antigen processing and/or presentation. To examine this possibility, we first focused on the expression of HLA-DR (mouse H-2 IA/IE) in IRF8 WT and KO models. Surprisingly, we found that deletion of IRF8 in five distinct B cell lymphoma models (murine A20, BCL1 and 2PK3 and human SU-DHL4, SU-DHL6,) did not meaningfully change H-2 IA/IE or HLA-DR cell surface expression (Figure 5A). We then considered the possibility that CD74 (Ia antigen-associated invariant chain, Ii) and/or HLA-DM, critical intracellular regulators of antigen processing/loading, which are also transcriptional targets of both CIITA and IRF8 ^30,32,33,54^, may be modified by IRF8 deletion (KO) or mutation. Using intracellular FACS analyses, we detected a significant suppression of CD74 and HLA-DM (H2-DM) in five independent models of IRF8 KO models (Figure 5B-C). Interestingly, in the mouse models, the effects of IRF8 were more robust towards CD74 whereas in human cells, HLA-DM suppression was more marked. Next, we queried the impact of IRF8 WT or mutant expression on CD74 and H2-DM levels. In the A20 and 2PK-3 mouse models, the expression of CD74, as well as H2-DM in A20, was significantly increased with ectopic expression of IRF8 WT but not in IRF8 mutant expressing cells (Figure 5D). Likewise, in the IRF8-null Toledo DLBCL cell line, the expression of CD74 and HLA-DM increased with ectopic expression of IRF8 WT but not of mutant IRF8 (Figure 5E, Supplemental Figure 5A). All of these changes were also captured at mRNA levels (Supplemental Figure 5B). The effects of IRF8 on CD74 and HLA-DM expression suggest that in IRF8 mutant tumors, correct antigen loading in the MHC CII complex may be impaired in at least two nodes. In brief, the abnormally lower expression of CD74/invariant chain (Ii), which physiologically associates with MHC CII αβ dimers, could result in incorrect loading of endogenously derived peptides at the endoplasmic reticulum. Through sequential proteolysis of the remaining li in IRF8-mutant cells, the CLIP (class II-associated invariant chain peptide) fragment is generated and remains bound to MHC CII (to prevent premature peptide loading), until it is dislodged by HLA-DM allowing for proper binding of mature antigen (reviewed in ^34^). Thus, the lower abundance of HLA-DM in IRF8 mutant cells may inefficiently dislodge CLIP and blunt the loading of the correct antigen into the mature MHC CII for cell surface presentation. If this latter assumption is correct, then in IRF8 KO or mutant cells, there will be more MHC CII-CLIP on the cell surface. Using, FACS we validated this prediction in two human DLBCL models (SU-DHL4 IRF8 KO, Toledo IRF8 WT and mutants) with detectable CLIP levels on the cell surface (Figure 5F, Supplemental Figure 5A). We note than an antibody that detects CLIP in mouse cells is not available, so these quantifications could not be extended to the IRF8 modified murine B cell lymphomas models. We concluded that in B cell lymphoma models, IRF8 KO and/or expression of mutant variants decrease CD74 and HLA-DM expression, potentially mainly deregulating intracellular antigen processing and loading, rather than MHC CII expression on the cell surface.

**Figure 5.**
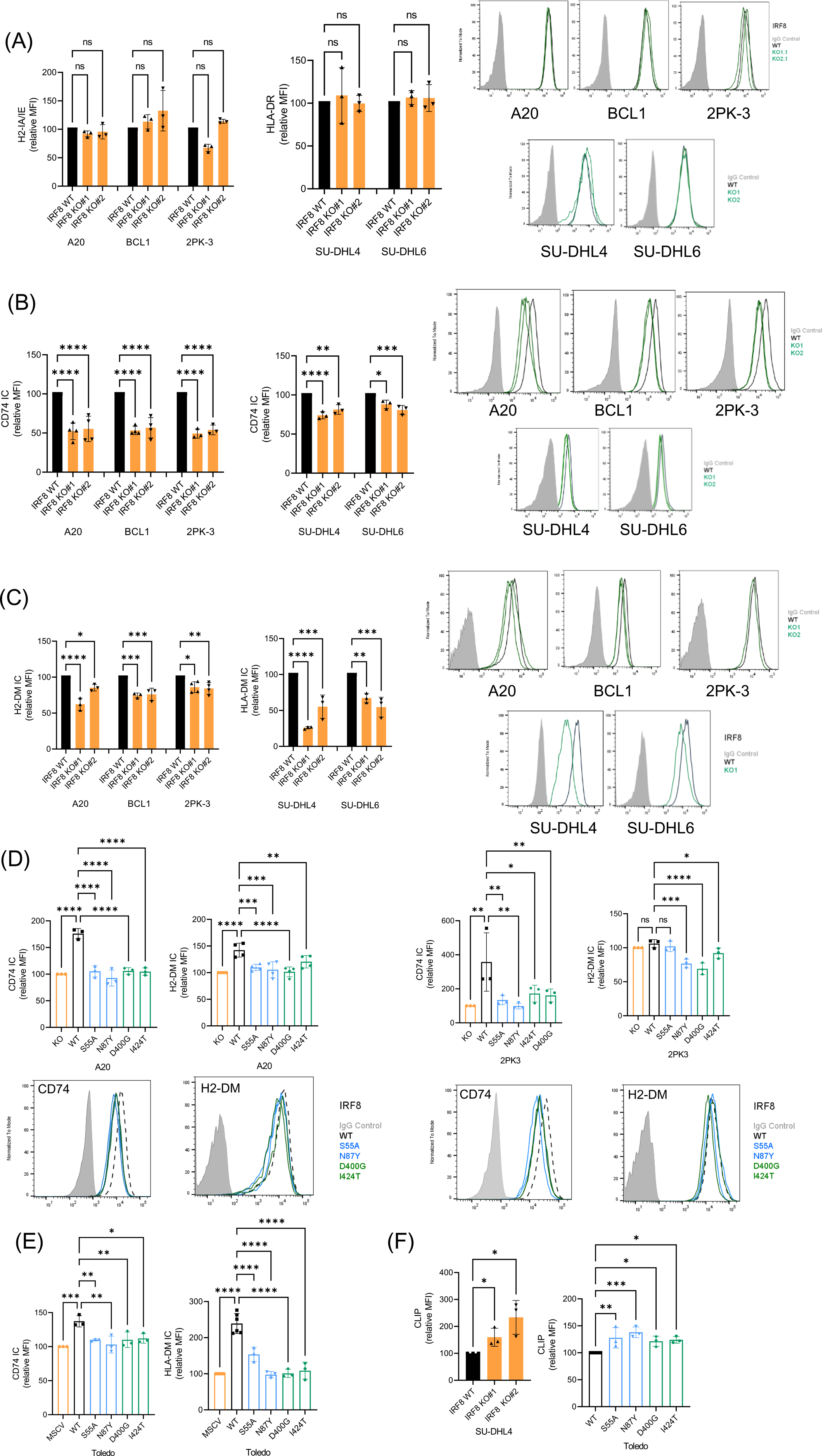
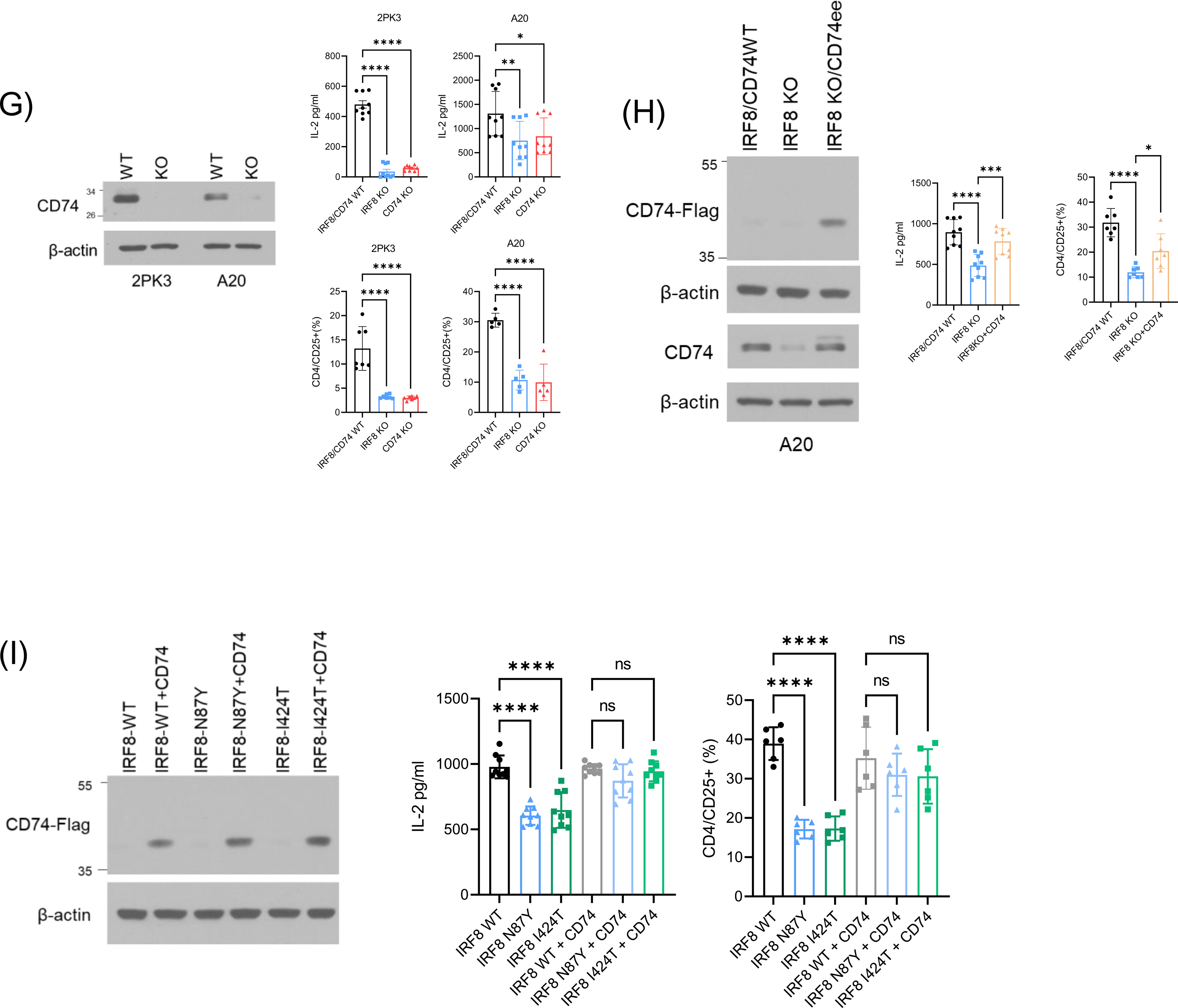
IRF8 control of components of the MHC CII complex. (A) FACS analysis of H2-IA/IE (human HLA-DR) in the cell surface of B cell lymphoma models of IRF8 KO – representative histograms shown to right. (B) FACS analysis of intra-cellular (IC) CD74 in B cell lymphoma models of IRF8 KO – representative histograms shown to right. (C) FACS analysis of intra-cellular (IC) H2-DM (human HLA-DM) in B cell lymphoma models of IRF8 KO – representative histograms shown to right. (D) **left panels** - FACS analysis of intra-cellular (IC) CD74 and H2-DM in the IRF8 KO A20 B cell lymphoma model “rescued” with stable expression of IRF8 WT or mutants; **right panels** - FACS analysis of intra-cellular (IC) CD74 and H2-DM in the IRF8 KO 2PK-3 B cell lymphoma model “rescued” with stable expression of IRF8 WT or mutants - representative histograms shown at the bottom. (E) FACS analysis of intra-cellular (IC) CD74 and HLA-DM in the human DLBCL cell line stably expressing and empty vector (MSCV), IRF8 WT or mutants. (F) FACS analysis of CLIP expression in the cell surface of SU-DHL4 (IRF8 WT vs. KO) and Toledo (IRF8 WT vs mutant). (G) **left**, WB analysis of CD74 KO models in mouse B cell lymphoma 2PK-3 and A20; **right, top and bottom**, IL-2 levels in the conditioned media and percentage of CD4/CD25+ expression in DO-11.10 cells co-cultured with IRF8/CD74 WT, IRF8 KO or CD74 KO APC models. (H) **left,** WB analysis of CD74 with IRF8/CD74 WT, IRF8 KO or IRF8KO+CD74-FLAG in A20 models with anti-FLAG or CD74 antibodies; **right,** IL-2 levels in the conditioned media and percentage of CD4/CD25+ expression in DO-11.10 cells co-cultured with IRF8/CD74 WT, IRF8 KO or IRF8KO+CD74 KO APC models. (I) **left**, WB analysis of ectopically expressed CD74-FLAG in the IRF8 WT, IRF8 N87Y and IRF8 I424T mouse B cell lymphoma A20 models; **right**, IL-2 levels in the conditioned media and percentage of CD4/CD25+ expression in DO-11.10 cells co-cultured with IRF8 WTT, IRF8 N87Y and IRF8 I424T (with or without CD74 ectopic expression) APC models. In all panels data are mean ± SD of three biological replicates. FACS analysis is displayed as relative mean fluorescence intensity (MFI). P values are from ANOVA, with Bonferroni or Fisher’s LSD post-test, or two-sided Student’s t-test. *(p<0.05), **(p<0.01), ***(p<0.001), **** (p<0.0001).

In murine B cell lymphoma models, which allow for in vitro (CD4 DO-11.10 cells) and in vivo (see below) functional examination of defective antigen presentation, CD74 was the main “target” of IRF8 effects (Figure 5A-D). To validate the relevance of this interplay, we used CRISPR-Cas9 to KO CD74 expression in the A20 and 2PK-3 B cell lymphoma models, and found that similarly to loss of IRF8, KO of CD74 resulted in significantly diminished OVA-induced CD4 activation (Figure 5G, Supplemental Figure 5C). These data reinforce the essentiality of Ii in preventing premature loading of antigen peptide into the MHC CII complex towards proper cell surface presentation ^34^, and eventual CD4 T cells activation, and start to point to a mechanism by which IRF8 dysfunction may facilitate immune escape. Next, and more importantly, we ectopically expressed mouse CD74 (p41 isoform) in the IRF8 KO A20 B cell lymphoma model and showed that this genetic modulation was sufficient to rescue the defective OVA-elicited CD4 activation (IL2 secretion and CD25 expression) associated with IRF8 KO (Figure 5H, Supplemental Figure 5D). Finally, to mitigate concerns that the ectopic expression of CD74 may enhance CD4 activation irrespective of IRF8 status, and to define if CD74 can rescue the IRF8 mutant phenotype, we tested CD74 effects on B cell lymphoma models expressing IRF8 WT, the DNA binding domain mutant N87Y, or the “c-terminal” domain mutant I424T. Reassuringly, we found that ectopic CD74 expression significantly increased OVA-driven CD4 activation in IRF8 N87Y and I424T, but it had a negligible effect on IRF8-WT cells (Figure 5I, Supplemental Figure 5E). We concluded that CD74 is an important mediator of IRF8 dysfunction in B cell lymphomas.

### IRF8 mutant B cell lymphomas display a T-cell deficient tumor microenvironment (TME) and increased aggressiveness – role of CD74

Our in vitro data suggested that expression of mutant IRF8 in B cell lymphomas deregulates antigen processing/loading and ultimately presentation in an MHC CII context thus eliciting a subpar CD4 response. Here, we investigated if these in vitro findings translated into an IRF8 status-driven remodeling of the TME, and if it influenced tumor growth in vivo. To that end, we examined multiple cohorts of syngeneic B cell lymphoma developed with A20 cells expressing IRF8 WT, N87Y or I424T (thus covering the two functional domains/clusters of IRF8 mutation in DLBCL) grown in syngeneic BalbC mice (attempts to engraft 2PK-3 cells in BalbC mice were unsuccessful). In all assays, each mouse was injected subcutaneously with 5 million A20 cells, tumor volume was quantified at defined intervals and immune landscape examined by FACS immediately following mice sacrifice. In the TME, we initially quantified the total T-cell infiltrate (CD3), the CD4+ and CD8+ subpopulations, T-regs (CD4+/CD127-, CD25+/intracellular Foxp3+), NK cells (NKp46+, i.e., CD335+), monocyte/macrophages (CD11b+), and the myeloid derived suppressor cells (CD11b+, Gr1+). Mice harboring lymphomas expressing IRF8 N87Y and I424T displayed significantly increased tumor growth (Figure 6A). Consistently, a significant depletion of T-cells (CD4+ and CD8+) was detected in the TME of lymphomas driven by an IRF8 mutant allele, irrespective of the variant analyzed (Figure 6B, Supplemental Figure 6A). In addition, T-regs were significantly enriched in IRF8 N87Y and I424T lymphomas, NK cells depleted in N87Y, while the monocytes/macrophages and MDSC infiltrate were quantitatively indistinguishable between IRF8 WT and mutant lymphomas (Figure 6C, Supplemental Figure 6A). The impact of IRF8 status on the in vitro growth pattern of these A20 models was unremarkable, with no significant difference between IRF8 WT and N87Y or I424T expressing cells (Supplemental Figure 6B). Together, these data confirm that the role of IRF8 mutations is principally unveiled in vivo furthering the concept that immune escape, rather than a lymphoma cell intrinsic effect, is at the core of IRF8 dysfunction in B-cell lymphomas. Importantly, the differences noted in respect to tumor burden and the TME infiltrate were not related to the levels of ectopic IRF8 expression, which were tightly controlled by protein (Figure 4B), mRNA (Supplemental Figure 4D) and GFP (bicistronic with IRF8) quantifications (Supplemental Figure 6C). Finally, in agreement with the suggestion that IRF8 may be an essential gene in lymphoma^37^, which could also explain why IRF8 mutations are mono-allelic, A20 IRF8 KO cells failed to fully engraft in the syngeneic BalbC mouse (Supplemental Figure 6D).

**Figure 6.**
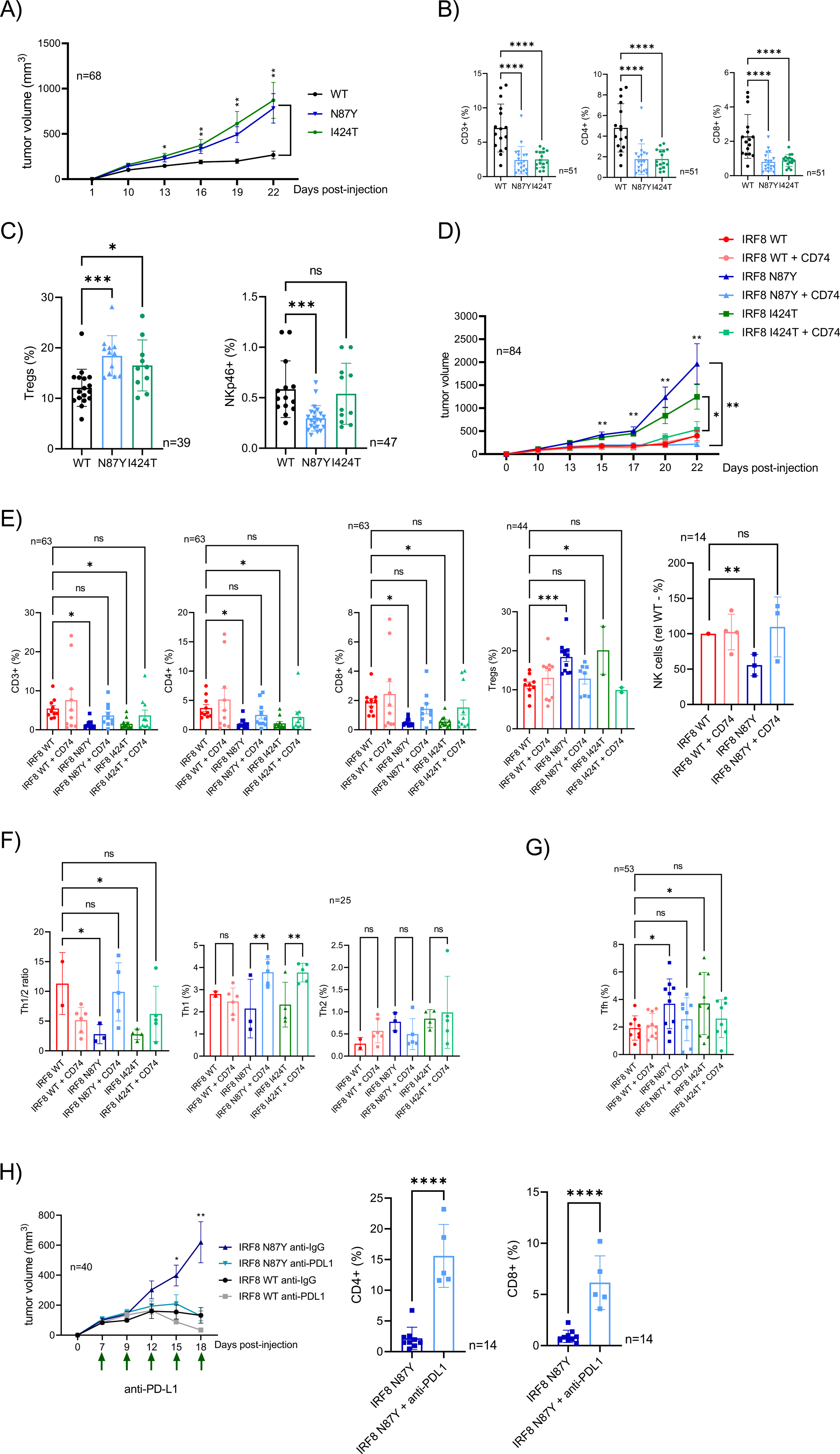
IRF8 effects on B cell lymphoma aggressiveness and immune microenvironment. (A) **left to right**, growth curve (volume) of lymphomas expressing IRF8 WT, N87Y or I424T. Data are mean ± SD of four independent cohorts; p values are from Mann-Whitney test. (B) **left to right**, FACS-based quantification of CD3, CD4 and CD8 T cells in the microenvironment of IRF8 WT or mutant lymphomas. (C) **left to right**, FACS-based quantification of T- regs and NK cells in the microenvironment of IRF8 WT or mutant lymphomas. In B) and C) data are mean ± SD of four independent cohorts; p values are from one-way ANOVA with Fisher’s LSD post-test. (D) growth curve (volume) of lymphomas expressing IRF8 WT, IRF8 N87Y or IRF8 I424T (with or without CD74 ectopic expression). Data are mean ± SD of two independent cohorts (n=84 mice); p values are from two-sided Student’s t-test – asterisk at the top is for IRF8 WT vs. N87Y or I424T, asterisk on the side is for IRF8 N87Y vs N87Y+CD74 and IRF8 I424T vs. I424T+CD74. (E) **left to right**, FACS-based quantification of CD3, CD4 and CD8, T-regs and NK cells in the microenvironment of lymphomas expressing IRF8 WT, IRF8 N87Y or IRF8 I424T (with or without CD74 ectopic expression). (F) **left to right**, Th1/Th2 ratio calculated from FACS-based quantification of Th1 and Th2 cells in the microenvironment of lymphomas expressing IRF8 WT, IRF8 N87Y or IRF8 I424T (with or without CD74 ectopic expression). (G) FACS-based quantification of Tfh cells in the microenvironment of lymphomas expressing IRF8 WT, IRF8 N87Y or IRF8 I424T (with or without CD74 ectopic expression); in E), F) and G), data are mean ± SD of multiple independent cohorts (n indicated in the figure) and the p values are from two-sided Student’s t-test. (H) **left to right**, growth curve (volume) of lymphomas models expressing IRF8 WT or IRF8 N87Y grown in mice treated with control antibody or anti-PDL1 antibody; CD4 and CD8 quantification in IRF8 N87Y lymphomas treated with ctrl control or anti-PDL1 antibody; tumor volume data are mean ± SEM of 40 mice, CD4 and CD8 data are mean ± SD of 14 lymphomas. P value is from two-sided Student’s t-test for IRF8 N87Y lymphomas treated with control vs anti-PDL1 antibody. For all panels, *(p≤0.05), **(p ≤ 0.01), ***(p ≤ 0.001), **** (p ≤ 0.0001).

Next, we investigated if the IRF8 N87Y and I424T phenotypes could be corrected by ectopic expression of CD74 in vivo. Tumor growth was significantly suppressed in IRF8 mutant lymphomas co-expressing CD74, while no difference was detected between IRF8 WT and IRF8 WT + CD74. (Figure 6D); once more only negligible differences were detected in in vitro growth rates supporting the role of the microenvironment in the differences detected in vivo (Supplemental Figure 6E). Indeed, this reduced tumor aggressiveness was accompanied, and possibly secondary to, a marked remodeling of the TME, which after CD74 ectopic expression was not significantly different between IRF8 WT and IRF8 mutant lymphomas. In brief, ectopic co-expression of CD74 increased CD3, CD4, CD8, and NK infiltrate, and decreased Tregs abundance in IRF8 N87Y and I424T lymphomas (Figure 6E, Supplemental Figure 6F). In an attempt to define the entire spectrum of IRF8 mutant associated changes in subpopulations of CD4 T-cells, we expanded the investigation to T helper type 1 (Th1), T helper type 2 (Th2), and T follicular helper (Tfh) cells. In IRF8 N87Y and I424T driven lymphomas, we detected a significantly smaller Th1/Th2 ratio, which was corrected to the IRF8 WT “baseline” by CD74 expression, primarily via modulation of the abundance of Th1 levels (Figure 6F, Supplemental Figure 6F). Furthermore, we found a significant increase in Tfh cell infiltrates in IRF8 mutant lymphomas, which was also normalized by CD74 co-expression (Figure 6G, Supplemental Figure 6F).

A recent report showed that IRF8 expression in tumor associated macrophages (TAMs) promoted T cell exhaustion and that, in agreement with our findings in B cell lymphoma models, IRF8 was required for TAMs’ to properly present cancer cell antigens^55^. T cell exhaustion is often a harbinger for diminished immune checkpoint therapy^3^. Therefore, we decided to examine if the IRF8 mutant-associated remodeling of the TME, and attendant growth advantage that we detected in B cell lymphoma models, was still amenable to correction with checkpoint inhibitors of the PD-1/PD-L1 pathway. To test this possibility, we designed a treatment trial in which, following tumor engraftment, mice harboring IRF8-WT or IRF8-N87Y lymphomas were dosed with an anti-PD-L1 antibody or control antibody every 72h. Anti-PD-L1 treatment was effective in inhibiting the highly aggressive IRF8 N87Y mutant, with accompanying increase in the CD4/CD8 infiltrate (Figure 6H). We are cognizant that nodal DLBCLs are notoriously unresponsive to checkpoint inhibitors^56^, and our examination here was not meant to test the impact of IRF8 mutation on that lack of response, but rather to try to complement the data from IRF8 overexpression on TAMs

We concluded that IRF8 mutation rewires the lymphoma microenvironment towards a broad pro-cancer profile. This perturbation increases tumor aggressiveness of IRF8 mutant lymphomas, which is clinically and immunologically corrected by CD74 expression, or anti-PD-L1 treatment.

### Impact of IRF8 mutation in the immune composition of the microenvironment of primary human DLBCL

To test if the IRF8 mutant driven remodeling of the lymphoma microenvironment found in mouse models of B cell lymphoma could be recapitulated in primary human DLBCL we utilized deconvolution of bulk RNAseq data (dbGaP Study Accession: phs001444.v2.p1). This cohort of 480 DLBCLs, was first re-analyzed for IRF8 mutation calling, and a total of 45 lymphomas were found to harbor an IRF8 variant allele (Supplemental Table 7). Next, we employed the xCell algorithm^57^ to identify putative distinctions in the non-malignant cell component of IRF8 WT and mutant DLBCLs. In agreement with the mouse B cell lymphoma models, IRF8 mutant DLBCLs displayed significantly diminished signatures of NK cell infiltrate as well as selected CD4 subpopulations, including Th1, but not Th2 (Figure 7A). In addition, the TME of IRF8 mutant DLBCLs was also significantly depleted of plasmacytoid dendritic cells (pDC), and of the related interstitial dendritic cells (iDC). Interestingly, NKT, Th1 and pDC all secrete proinflammatory cytokines, chiefly among them IFN-γ ^58-60^. Still, using xCell, we did not detect significant changes in T-regs in IRF8 mutant primary DLBCL (Supplemental Figure 7A), which were consistently detected in the A20 BalbC models (Tfh cells are not uniquely identified with xCell). This result was somewhat surprising because recently IRF8 mutation in DLBCL was identified as a putative marker for poor response to CD19-CART cells^61^. Moreover, in that study it was postulated that this worse outcome could be related to an increase in T-regs in the microenvironment of IRF8 mutant DLBCL, which therein was detected by deconvolution of bulk RNA-seq signatures using another algorithm, CIBERSORTx^62^. Thus, we reanalyzed the primary DLBCL data with CIBERSORTx, and in a cohort that excluded tumors with mutations in other genes known to deregulate antigen presentation (CIITA, B2M, CREBBP, EP300 and EZH2), we detected an increase in T-regs (p=0.05) in IRF8 mutant DLBCLs (Figure 7B, Supplemental Figure 7B). These data also mitigate concerns that, due to the high frequency of IRF8 mutations in the EZB subgroup (Figure 1C), part of the IRF8 signature may derive from co-occurring EZH2 mutation. Importantly, using CIBERSORTx, and also adding a third algorithm, MCP-counter^63^, we confirmed depletion of NK cells as the most reproducible change in the TME of IRF8 mutant human DLBCL (Figure 7C, Supplemental Figure 7C). Of note, Th1 cells and pDC are not uniquely identified subpopulations in deconvolution performed with CIBERSORTx or MCP counter, thus the xCELL data on those subsets could not be cross validated. We concluded that in the highly heterogenous setting of primary DLBCL, the presence of an IRF8 mutation associates with a TME signature characterized by depletion of a series of pro-inflammatory/anti-cancer cell types (of the innate and adaptive immune system) and a potential increase in the suppressive T-regs. This profile partially recapitulates the microenvironment remodeling detected in the mouse model of IRF8 WT or mutant B cell lymphoma that we developed.

**Figure 7.**
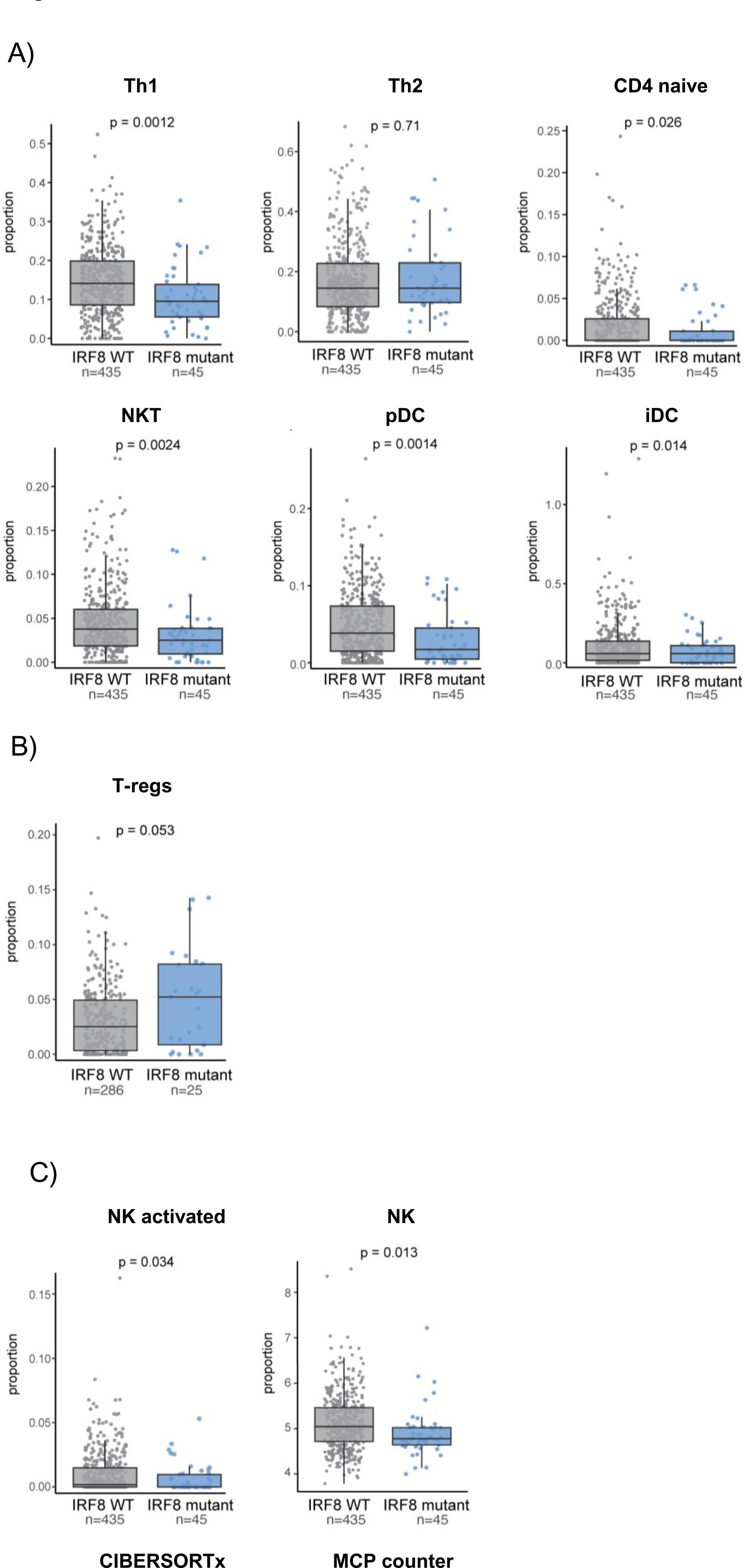
Immune composition of the microenvironment of IRF8-mutant human primary DLBCLs. (A) **From left to right, top to bottom**. xCELL-defined scores estimatimation of follicular T cell (Th1), Th2, naïve CD4, NKT, plasmacytoid dendritic cells (pDC), immature dendritic cells (iDC) in IRF8 WT (n=435, gray box plot) or IRF8 mutant (n= 45, blue boxplot) DLBCLs. (B) Estimated T-regs proportions in IRF8 WT (n=286 gray box plot) or IRF8 mutant (n= 25, blue box plot) measured using CIBERSORTx. (C) **left to right** - Estimated proportions of NK activated (CIBERSORTx) or estimated scores of total NK cells (MCP counter) in IRF8 WT (n=435, gray boxplots) or IRF8 mutant (n=45, blue boxplots) DLBLCs. In all panels the boxplots show median and inter-quartile range. P values are from two-sided Students’ t-test.

### Structural modeling of IRF8 mutation

To gain insights into the putative mechanisms by which IRF8 mutations may deregulate protein function, we performed a comparative structural analysis using experimental structures of the DNA binding domain (DBD) of IRF3 (PDB: 2PI0) bound to a double-stranded DNA^64^ (Figure 8A). No experimental structure is yet available for partial or full-length IRF8, thus, we used a 3D model of IRF8 as predicted by AlphaFold^65^. The AlphaFold model shows a bilobal architecture of IRF8 with its N-terminus DNA binding (DBD; aa 1-116) connected via a flexible linker (aa 117-169) to the “c-terminal domain” (CTD), which is composed of a well-defined protein-protein interaction (aa 170-382) and a still uncharacterized c-terminal region (aa 383-426) (Figure 8B). Next, we overlayed the IRF3 and IRF8 DBDs (Figure 8C), which enabled us to examine IRF8’s protein-protein and protein-DNA interactions in detail and model the putative mechanism of protein dysfunction associated with the mutations that we characterized in vitro and in vivo.

**Figure 8.**
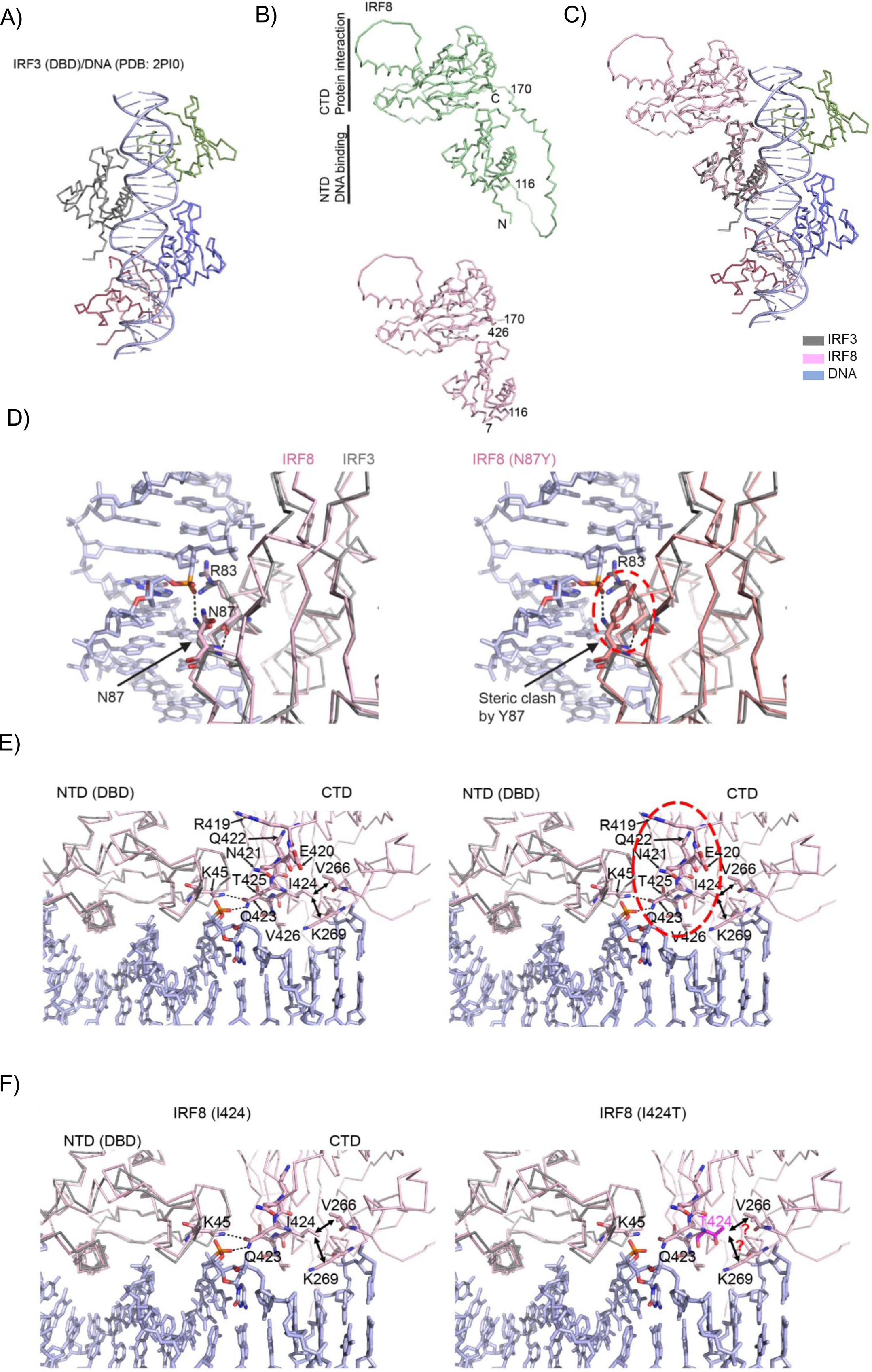
Computational modeling of human IRF8. (A) Crystal structure of IRF3 DBD/DNA complex (PDB: 2PI0).(B) **Top and bottom** - AlphaFold model of full-length IRF8 or IRF8 with the two flexible loops (aa 1-6 and 117-169) omitted. CTD, c-terminus domain; DBD, DNA-binding domain. (C) Overlay of the DNA binding domains of IRF3 (grey) and IRF8 (pink). DNA-bound to IRF3 DBD is shown in light blue. (D) **left** - close-up of the N87 residue in the IRF8 DBD in contact with DNA; **right** - Steric clash by Y87 side chain, highlighted by red circle. (E) **left to right** – c- terminal tail residues (R419 to V426) may mediate electrostatic interactions with DNA and DBD and are sandwiched between the DBD and CTD. (F) **left** - I424 side chain faces the interior of CTD and may stabilize it via interactions with V266 and K269, and the polar side chain of Q423 may form h-bonds to K45 from DBD and the phosphoryl oxygens of the DNA backbone; **right** - the polar side chain of mutant T424 may disrupt the hydrophobic interior of CTD and displace Q423 from its original position, resulting in a loss of interaction with DBD and DNA.

Upon superimposing the IRF3-DNA structure with the IRF8 model, we found that the polar side chain of N87 is within the h-bonding distance of the DNA backbone, akin to N85, the equivalent residue in IRF3. A bulkier and hydrophobic side chain of N87Y mutant will lose this critical h-bond and electrostatic interaction with DNA. Moreover, a Y at position 87 can sterically clash with the side chain of R83, another important amino acid required for charge-charge interaction with DNA (Figure 8D). Another DBD mutation that we studied in vitro is S55A (Figures 2-5). S55 locates within the α2 helix of the IRF8 DBD (Supplemental Figure 8A). Its side chain stabilizes the loop1 between α2 and β1. This loop enters the minor groove of DNA. An alanine at position 55 may result in loss of this stabilizing interaction and negatively impact DNA binding (Supplemental Figure 8A). Thus, phenotypic changes detected in vitro and/or in vivo in B cell lymphomas expressing the S55A or N87Y mutations are possibly related to loss of, or sub-optimal, binding of IRF8 to DNA.

The CTD of IRF8 is predicted to primarily promote interaction with other proteins^66^. However, the AlphaFold model of the full-length IRF8, and its comparative analyses with DNA-bound IRF3 structure, revealed the orientation of IRF8’s CTD close to the upstream portion of DNA and suggested that in the very c-terminus tail (aa 419-426) sandwiches between DBD and CTD, thus acting as a hinge to stabilize the two domains and bring the CTD close to DNA (Figure 8E). In a close examination of I424, we found that its side chain faces the interior of CTD to stabilize it via hydrophobic interactions with V266 and the aliphatic portion of the side chain of K269 (Figure 8F). In addition, the polar side chain of Q423 faces outward to form bidentate h-bonds to K45 from DBD and the phosphoryl oxygens of the DNA backbone. We predict that the polar side chain of the I424T mutant that we examined in vitro and in vivo will disrupt the hydrophobic interior of CTD. As a result, Q423 may be displaced from its original position, resulting in a loss of interaction with DBD and DNA (Figure 8F). Another CTD mutation that we examined in vitro is D400G. An overlay of AlphaFold IRF8 model and IRF3 CTDs reveals that D400 lies close to the putative protein-protein interface of IRF8, in an “outer loop”; a G at position 400 may influence the flexibility of this region affecting the potential protein-protein interaction (Supplemental Figure 8B).

We concluded that the similarity in phenotype that we detected between DBD and c-terminal tail mutants (N87Y and I424T, respectively) may reflect the fact that both mutants could disrupt IRF8/DNA contact. This unexpected finding may explain the high frequency of truncating mutations clustered at the c-terminal tail of IRF8.

## Discussion

Here, we showed that B cell lymphomas carrying *IRF8* mutations are better equipped to evade the immune system than those with IRF8 in a WT configuration. Mechanistically, this escape is mediated by a defective expression of IRF8 targets that tightly control antigen loading into the MHC CII complex, including the CD74/invariant chain (Ii) and HLA-DM, as well as CIITA, the master regulator of MHC CII associated antigen presentation^34^. In the models that we developed, mutant IRF8 (or IRF8 KO) did not meaningfully change the cell surface expression of MHC CII. This interesting finding may explain why IRF8 was not identified in a broad screen that attempted to define the genetic basis for acquired deficiency of MHC expression in DLBCL^7^.

In our model system, ectopic expression of CD74 was sufficient to rescue the immune and clinical phenotypes detected in IRF8 mutant B cell lymphomas but it did not significantly change the immune landscape or growth pattern of IRF8 WT B cell lymphomas. These data place IRF8 at a control hub for antigen loading and reinforces the concept that these processes must be executed with precision, lest antigen presentation is defective, and, in a cancer context, immune evasion is elicited. Fortuitously, we detect a significant impact of IRF8 KO or mutation in an assay (CD4 DO-11.10 cells activation) in which the antigen loaded into the lymphoma cells was already processed (the OVA peptide). These findings, together with defective expression of CD74 and HLA-DM, and the CD74-mediated rescue of IRF8 mutant lymphomas, suggest that of the many steps involved in the antigen processing/presentation cascade, IRF8 impinges primarily on loading, and possibly not on acquisition, tagging, proteolysis, trafficking, and display^34^.

The mutation pattern of IRF8 in DLBCL (and related B cell malignancies) is similar to that of *CREBBP*, *EP300*, *KMT2D, TET2*, among other genes^12,13,36,37^. The variants are predominantly missense or truncating, often hemizygous, clustered in important domains of the protein, and functionally hypomorphic. A model of haploinsufficiency can be ascribed to this pattern. However, we suggest that IRF8 may be unique in its lymphomagenic potential since it has also been associated with a variety of oncogenic structural defects (translocation, amplification, super-enhancer deregulation^15-17^) in which excess of IRF8 expression may contribute to lymphoma development and progression. In agreement with this postulate, in an unbiased CRISPR screen in DLBCL cell lines, IRF8 was identified as an “essential oncogene”^37^, and we found the IRF8 deletion abolishes B cell lymphoma growth in vivo. Together, we propose that both gain or loss of IRF8 function play a role in B lymphoma biology, possibly by influencing cell intrinsic processes and immune evasion, respectively.

In mice, the IRF8-mutant driven remodeling of the lymphoma microenvironment was diverse, but consistently reflected a pro-cancer profile (decrease in CD4, CD8, NK, Tfh1 and increase in T-regs and Tfh)^60^. The initiating event, we postulate, was the defective antigen loading (and its eventual display on the cell surface) since ectopic expression of CD74 restored the entire repertoire of changes to an “IRF8 WT baseline”. Of the immune perturbations that we detected in the in vivo model of IRF8 mutant B cell lymphoma, previous associations existed with NK cells depletion and T-regs accumulation. In respect to the former, IRF8 mutations (some of which are also present in DLBCL) were found in familial cases of human NK deficiency^23^. In those instances, biallelic germline IRF8 variants were associated with decreased NK numbers, as well as cell defective NK cell maturation and function. In addition, in a mouse of conditional Irf8 deletion in NK cells, IRF8 was found to be essential for NK cell proliferation, expansion and adaptive cell responses^67^. However, in these two instances, IRF8 is either mutated in the germline or specifically deleted in developing NK cells, thus not fully recapitulating our model (or primary DLBCLs), wherein only the B cell lymphoma cell is IRF8 mutant. Interestingly, loss of IRF8 can also impair NK function in a cell extrinsic manner, including defective cytokine secretion by IRF8 null dendritic cell^68,69^. Thus, in the context of IRF8 mutant B cell lymphoma, it is possible that still to be identified secreted factors may limit NK recruitment to the tumor milieu. A similar process may be operational towards the plasmacytoid DC, which although not measured in the in vivo mouse model, were significantly depleted in the IRF8 mutant primary DLBCLs. Thus, it is possible to envision a circuitry wherein the poor antigen presentation by IRF8 mutant lymphomas (due to deregulated CD74/antigen loading) blunts the activation and differentiation of naïve CD4+ T cells towards the Th1 phenotype, decreasing the secretion of pro-inflammatory cytokines such as IFN-γ, which in turn limits the recruitment of NK, NKT and pDC cells, all themselves significant sources of IFN-γ and TNFα^58,59^, thus further suppressing the beneficial inflammatory profile of the TME.

The potential link between IRF8 and T-regs emerged more recently, when IRF8 mutations in relapsed DLBCL was identified as a putative marker for poor response to CD19-CAR T cells, possibly in association with an increase in T-regs in the lymphoma microenvironment^61^. Our in vivo model of IRF8 mutant B cell lymphoma validated this initial observation and linked it to defective antigen loading into MHC CII complexes, since it could be rescued with CD74 ectopic expression. Further, using the same deconvolution algorithm employed in the CAR-T study, we detected an increase in T-regs in a subset of IRF8-mutant DLBCLs. Lastly, we also detected a significant increase in Tfh cells in the mouse models of IRF8 mutant B cell lymphoma, which was corrected by CD74 ectopic expression. This finding made immediate sense given the role of these cells in supporting B cell expansion in germinal centers^70^, their role in the lymphomagenesis associated with Cathepsin S- and *TNFRSF14*-mutant lymphomas^71,72^, and their positive association with poor prognosis in B cell lymphomas^73^. However, this finding was not validated in primary DLBCL tumors, although this examination was limited by the fact that only one of the three algorithms that we used to deconvolute the bulk RNAseq data identifies Tfh cells as a discrete sub-population. Further studies in primary DLBCLs are needed to confirm or refute our mouse model data.

By computational modeling of IRF8 structure and building on the solved structure of the related IRF3 protein in complex with DNA, we gained insight into the mechanisms by which IRF8 mutations may deregulate protein architecture and function. Surprisingly, these preliminary observations implied that in addition to the well-defined N-terminal DBD, the C-terminal tail of IRF8 may also contact the DBD/DNA. This unexpected finding may provide clues to the similarity in phenotype that we detected between DBD and C-terminal tail mutants (N87Y and I424T, respectively), and it could help explain the high frequency of truncating mutations clustered at the C-terminus of IRF8.

Our study has limitations. The impact of IRF8 mutation on HLA-DM expression appears to be stronger in human DLBCL than in murine B cell lymphomas whereas CD74 defect is clearer in the latter. The selection to examine further CD74, instead of HLA-DM, was dictated by the model system but it is possible that in a human system, ectopic expression of HLA-DM may also rescue the IRF8 mutant phenotype. The IRF8 mutant models that we examined in vitro and in vivo are monoallelic, and it is possible that the presence of WT allele may uncover further aspects of the IRF8 role in B cell lymphomas. Nonetheless, all comparisons were made to a similarly generated IRF8 WT model, mitigating concerns that IRF8 dosage, instead of the nature of the IRF8 isoform expressed, accounted for the results.

## Material and Methods

### Cell lines

The DLBCL cell lines SU-DHL4 (male), SU-DHL6 (male), Toledo (female), the mouse B cell lymphoma cell line A20 (Balb/c), 2PK-3 (Balb/c), the B cell leukemia/lymphoma cell line BCL1 (Balb/c), and the mouse macrophage cell line RAW 264.7 were cultured at 37°C in 5%CO2 in RPMI-1640 medium (Invitrogen) containing 10% (vol/vol) fetal bovine serum (FBS). The HEK-293T cells (Thermo-Fisher Scientific) were maintained in Dulbecco’s modified Eagle media (DMEM; Mediatech) with 10% FBS, as we described^74^. The identity of the cell lines was confirmed by VNTR analysis and verified online at the DSMZ and ATTC cell banks and tested for Mycoplasma. All cell lines were preexistent in the investigator’s laboratory and/or were earlier obtained from ATCC or Thermo-Fisher.

### Establishment of genetic models. Human DLBCL cell lines

IRF8 was knocked-out in SU-DHL4 and SU-DHL6 IRF8 using CRISPR-Cas 9 technology. In brief, two distinct guide RNAs (Supplemental Table 8), were independently cloned into lentiCRISPR v2 plasmid, which was co-transfected with the PAX2 and pMD2.G plasmids into HEK-293T cells to generate lentivirus, as we described^75^. Subsequently, the SU-DHL4 and SU-DHL6 were transduced twice by spinoculation, selected with puromycin selection and clones generated by limiting dilution, as we reported^76^. The DLBCL cell line Toledo, which does not express endogenous IRF8, was transduced with empty MSCV-eGFP or IRF8 WT and mutants S55A, N87Y, D400G, I424T, all in a MSCV-eGFP backbone. In brief, IRF8 WT or mutant was generated by RT-PCR and site directed mutagenesis, as we described^77^, and Sanger sequence validated. These constructs were co-transfected with VSVG and pKAT into HEK-293T to produce retrovirus. The retrovirus was transduced twice into Toledo cells by spinoculation, followed by GFP sorting using FACS. **Mouse cell lines**. IRF8 KOs were generated in the macrophage cell line RAW 264.7, in the B cell lymphoma cell lines A20 and 2PK3 and in the B cell leukemia/lymphoma cell line BCL1 (all from ATCC), as described above for human cell lines, except for the use of four independent guide RNAs (Supplemental Table 8). In each cell line, at least two KO clones (obtained by limiting dilution) derived from different guide RNAs were used in downstream assays. In addition, A20 and 2PK- 3 IRF8 KO clones, were transduced with human IRF8 WT or mutants (S55A, C84R, N87Y, D400G and I424T) cloned into a MSCV-eGFP backbone, followed by FACS sorting to purity (>95% GFP+ cells), as described above for human cell lines. CRISPR-Cas9 technology, with guide RNA cloned into lentiCRISPR v2 plasmid, was also used to KO cd74 in A20 and 2PK-3 cell lines. Lastly, murine CD74 (p41 isoform) cloned into a lentivirus backbone with an MSCV promoter (pLV[Exp]-Neo-MSCV>FLAG/mCd74, Vector Builder, Inc), was transduced in A20 IRF8 KO cells, as well as A20 irf8 KO cells stably expressing IRF8 WT, IRF8 N87Y and IRF8 I424T. All KO and ectopic expression models were confirmed by sequencing and/or Western blotting.

### Mice and lymphoma development in vivo studies

Six-week-old male and female BALB/cJ mice were purchased from the Jackson Laboratory (Strain #:000651). Each mouse was injected subcutaneously with 5 million A20 cells murine B lymphoma cell line A20, as we described^78^. Six independent cohorts were generated (#1 = 25 mice, #2 = 25 mice, #3 = 40 mice, #4 = 20 mice, #5 = 60 mice, #6 = 30 mice). The A20 cell models utilized were generated by a CRISPR-Cas9-mediated knockout of mouse Irf8 (lentiCRISPR-V2 puromycin lentivirus), and subsequent stable ectopic expression of empty MSCV-eGFP, IRF8 WT, N87Y or I424T, alone or together with murine CD74 (p41 isoform) cloned into a lentivirus backbone with an MSCV promoter (pLV[Exp]-Neo-MSCV>FLAG/mCd74, purchased from Vector Builder, Inc). The mice were monitored daily and starting 10 days after cell injection, the subcutaneous tumors were measured every 72h using an electronic caliper, as we described^79^. In one cohort, mice harboring lymphomas expressing IRF8 WT or N87Y were randomized to receive five doses of anti-mouse PD-L1 (clone 10 F.9G2, Cat# BE0101, Invivomab) or rat IgG2b isotype control (Cat# BE0090, Invivomab), administered IP (200 μg per injection) at 2–3 days intervals. Following mice sacrifice, all the collected tumors were weighed, measured, photographed and single cells immediately generated for downstream FACS analysis. A local colony of TCR (Vα2, Vβ5) transgenic mice (OT-I) (Strain #:003831, Jackson Laboratories) was maintained by breeding homozygous transgenic mice. Male and female mice, 8-12 weeks old, were euthanized, spleens harvested and T lymphocytes isolated, and CD8 cells differentiated in vitro for antigen presentation assays (see below). All the animal procedures were approved by the Animal Care and Use Committee of the UT Health San Antonio.

### Data retrieval and somatic variants selection

The data used in this study was obtained from the Genomic Variation in Diffuse Large B Cell Lymphomas project (project ID NCICCR-DLBCL) available on the GDC data portal (dbGaP Study Accession: phs001444.v2.p1). The .bam files obtained from the GDC data portal had already been aligned to the GRCh38.p0 reference genome using BWA, the Mark Duplicates, and BQSR workflows. The variants in the aligned files were called and selected as previously reported^13^ with the modifications listed below. In brief, we defined the genomic regions of interest, utilized the VarScan2 software^80^, and selected the parameters of a read count greater than or equal to 3, and a variant read frequency greater than or equal to 0.1 to select relevant variants. To identify variants that had the highest impact on gene function, we implemented a series of filtering steps. We applied specific exclusion criteria to filter out variants with a population frequency above 0.0001 in the Genome Aggregation Database (gnomAD). We restricted our analysis to missense or nonsense point mutations, truncating mutations, and exonic indels. Additionally, we considered the variant selections made previously^13^ to finalize our list of somatic variants for downstream analysis. Variant allele frequency was obtained from VarScan2 or cBioportal cohorts listed in Supplemental Table 1. Classification in COO groups or subgroups were assigned from the original reports (Supplemental Table 1) or by the LymphGen data portal (https://llmpp.nih.gov/lymphgen/lymphgendataportal.php?version=2.0#).

### Mutual exclusivity analysis

We performed a mutual exclusivity analysis using the Oncoprint function available in cBioPortal^81^. We first selected a set of genes of interest based on their known relevance in DLBCL. We then used the Oncoprint function to visualize the mutual exclusivity patterns of these genes in our selected dataset. We defined mutual exclusivity as the absence of co-occurrence of mutations between two or more genes within a sample. We calculated the statistical significance of the mutual exclusivity using the Fisher’s exact test and corrected for multiple testing using the Benjamini-Hochberg procedure. We considered an stringent q-value (FDR) of <0.05 as statistically significant.

### Reporter Assay

The CIITA reporter constructs (-709 to +100 of CIITA promoter region), cloned into pgL3 in WT configuration or with point mutations in the Ets-IRF composite element (EICE) site, were a gift from Kenneth Wright, Moffit Cancer Center, and had been characterized earlier^48^. Their identity was confirmed by Sanger sequencing (EICE WT site GGTTTTCACTTC, mutant G<u>CAG</u>TTCACTTC). The CIITA WT or EICE mutant promoter vectors were independently co-transfected with pCMV β-gal plasmid into RAW 264.7 cells seeded at 5 x 10^5^ in 12- well plates. The RAW 264.7 cells were first subjected to CRISPR-Cas9-mediated knockout of Irf8 (CRISPR-V2 puromycin lentivirus) and subsequently transduced to stably express human IRF8 WT or mutant (S55A, N87Y, D400G, I424T) in the MSCV-eGFP retrovirus, as we reported^82^. Twenty-four hours after transfection, the RAW 264.7 cells were washed with ice-cold PBS, and harvested in 250 µl of reporter lysis buffer (Promega luciferase assay system) with gentle shaking for 15 min at room temperature. The lysate mixture was spun at 14.000 rpm for 30 sec and the supernatant collected for the luciferase assay. Briefly, 20 µl of lysate was added to a 1.5 ml Eppendorf tubes containing 100 µl of luciferase assay reagent II (Promega luciferase assay system, Cat# E1500) and the mixture pipetted in and out three times. The luminescence reading was performed in Modulus single tube multimode reader (Turner Biosystems). The β-Galactosidase activity was measured in 30 µl of lysate incubated with 20 µl of reporter lysis buffer and 50 µl of 2X β-gal Enzyme Assay buffer (Promega, Cat# E2000). The absorbance was read in iMark microplate reader (BioRad), in 96-well plates, following incubation at 37°C for 2h and addition of 150 µl stop solution (1 M sodium carbonate).

### Antigen presenting assay – DO-11-10 CD4 T cells

Mouse B cell lymphoma B cell lines, A20, 2PK-3 and BCL-1 were plated at 5 x 10^4^ cells/well/100µl culture media in 96-well plates, followed by “antigen loading” with the addition of the ovalbumin peptide (OVA-323-339) (Cat# 12787-01, Bio-Synthesis) for 3 hours. A20 and BCL1 cells were “loaded” with 5µg/ml whereas 2PK-3 cells were loaded with 10ug/ml of OVA. Cells were then washed with growth medium and co-cultured with DO-11.10 CD4 T cells, as reported^51^; 8 x 10^4^ of DO-11.10 cells in 200µl of culture media were added to the A20 and BCL1 cells whereas 1.3 X 10^5^ of DO-11.10 cells were added to 2PK3 cells per well. The plates were centrifuged at 500g for 5 minutes to initiate contact between the cells and placed at 37°C in a CO2 incubator for multiple time points. In brief, after 6 hours in co-culture, supernatants from the A20 and DO-11.10 cells were collected for IL-2 detection by enzyme-linked immunosorbent assays (ELISA) using commercially available sandwich-based kit (R&D systems, Cat# M2000), according to the manufacturer’s instructions. After 24 hours in co-coculture, these cells were harvested and analyzed for surface expression of CD4 and the T cell activation marker CD25 by flow cytometry as described below. Supernatants from BCL1 and 2PK3 cells in co-culture with DO-11.10 were collected after 24 hours of incubation and analyzed for IL-2 detection by ELISA. At the same time, cells were harvested for surface expression of CD4 and CD25.

### Isolation/activation of CD8 T cells from OT-I mice and in vitro cytotoxicity assay

Spleens were collected from 8- 12 week old TCR (Vα2, Vβ5) transgenic mice (OT-I) (Strain #:003831, Jackson Laboratories) and T lymphocytes purified using Easy Sep^TM^ Mouse T cell isolation kit (STEMCELL Technologies, Cat# 19851A), as we described^83^. To generate CD8+ T lymphoblasts responsive to ovalbumin peptide (OVAp; SIINFEKL OVA257–264, Cat# S7951, Sigma-Aldrich), T cells were incubated for 48h with OVAp (10 ng/ml) and mouse recombinant IL2 (50 U/ml) (Cat# 575402, Biolegend), was added for a period of five additional days, as described^84^, at which time FACS analysis confirmed that 99.9% of the cells were CD8+. Subsequently, these activated CD8+ T lymphoblasts were co-cultured with IRF8 WT or KO mouse B cell lymphoma cell lines (A20, 2PK-3 and BCL1). In brief, lymphoma cell lines were pulsed with 1 µM OVAp for two hours, washed twice with PBS and cocultured with mice CD8+ T cells at an effector to target ratio of 5:1 for 6h at 37°C, after which the cytotoxicity towards IRF8 WT or KO lymphoma cells was assessed by FACS using 7-aminoactinomycin D (7-AAD) (BD Biosciences, Cat# 51-68981). Controls of non-OVAp pulsed B cell lymphomas were included in all instances. All assays were subjected to three biological replicates.

### Granzyme B detection assay

Conditioned media from co-culture of OT-I activated CD8+ T cells with mouse B cell lymphoma (OVAp pulsed and non-pulsed controls) were assessed for the concentrations of Granzyme B by ELISA using mouse Granzyme B specific kit (RayBiotech, Cat# ELM-GranzymeB-1), according to the manufacturer’s instructions. In brief, after 6h of co-culture, cell supernatants were diluted 1/100 in an assay diluent buffer provided and loaded on an ELISA plate pre-coated with mouse Granzyme B antibody for 2.5 hours at room temperature. Plates were washed followed by an hour thirty-minute incubation with secondary biotinylated anti-mouse granzyme B antibody. HRP-conjugated streptavidin was then added to the plates for an hour at room temperature. A colorimetric signal was developed on the enzymatic reaction of HRP with a chromogenic substrate, 3,3’,5,5’-tetramethylbenzidine (TMB) for 30 minutes. After stopping the reaction with 0.2 M sulfuric acid, the values were determined by measuring the optical density (OD) of ELISA plate wells at 450 nm. Concentrations were calculated in pg/ml based on the standard curve generated. All assays were subjected to three biological replicates.

### Flow cytometry - cell surface or intracellular staining – cell line models

1 million viable human DLBCL and mouse B lymphoma cell lines were counted and resuspended in 100 µl FACS buffer (2% FBS, 1mM EDTA in PBS). Cells were stained with the respective fluorochrome-conjugated antibodies for detection of HLA-DR, H2-IA/IE or CLIP surface proteins and incubated for 30 minutes at 4 °C. Cell were then washed with FACS buffer and resuspended in FACS buffer containing DAPI (5 mg/ml, diluted 1:5,000) for live/dead cell exclusion. For intracellular detection of HLA-DM, H2-DM or Li invariant chain/CD74, cells were fixed and permeabilization using the Intracellular Fixation & Permeabilization Buffer kit (eBioscience, Cat# 88-8824-00) and stained with the corresponding antibodies. **Flow cytometry - cell surface or intracellular staining – in vivo lymphoma microenvironment**. To quantify and characterize the immune infiltrates in the syngeneic lymphoma engraftment models, tumors were harvested, and single cells suspensions were generated, as reported^85^. Two million viable cells were resuspended in 100µl FACS buffer with 1 µl of Fc Block (anti-mouse Fc block CD16/32, Cat# 101302, Biolegend) for 20 minutes at 4°C. Cells were then washed with FACS buffer and stained with surface markers antibodies to detect the following populations: 1) T-cell subpopulations (CD3, CD4, and CD8 surface markers), 2) Monocytes/macrophages subpopulations (CD11b surface marker), 3) Natural killer cells population (Nkp46 surface marker), 4) Myeloid-derived suppressor cells (MDSC) population (double positive CD11b, Gr1 surface markers), 5) T-helper-1 (Th1) cell population (CD4, CXCR3 and CCR5 positive surface markers), 6) T-helper-2 (Th2) cell population (CD4, CCR3, and ST2 positive surface markers), 7) T-follicular-helper (Tfh) cell population (CD4, CXCR5, and PD-1 positive surface markers). Cells were washed with FACS buffer and resuspended in FACS buffer containing a viability dye 7AAD (Cat# 51-6898, BD Bioscience) or DAPI (Cat# 62248, Thermo Fisher) (1:5.000 dilution), according to the fluorophores used. Fluorescence compensation was performed using UltraComp eBeads™ (Cat# 01-2222-42, Invitrogen). For *ex vivo* detection of T-regulatory cells (T-regs), 6 million viable single cells harvested from mouse lymphomas were surface stained with BV421 anti-mouse CD4, APC anti-mouse CD127 (IL-7Rα) and BV650 anti-mouse CD25, followed by fixing/permeabilization using the True-Nuclear transcription factor buffer set (Biolegend) according to the manufacturer’s instructions. Cells were next stained with PE anti-mouse FoxP3 antibody. CD4+/CD127- cells were analyzed for CD25 and intracellular FoxP3. A pre-incubation with Fc blocking antibody was performed. Instrument compensation was performed with UltraComp eBeads^TM^.

Antibodies used for flow cytometry were the following: Anti-human HLA-DR and Anti-human HLA-DM, (Cat# 361610 and Cat# 358004, Biolegend), Anti-human CD74 (cat# FAB35901N, R&D Systems). Anti-mouse H2I-A/I-E, (Cat#10763), Anti-mouse CD3 (Cat# 100206), Anti-mouse CD8a (Cat# 100712), Anti-mouse CD11b (Cat# 101208), Anti-mouse Nkp46 (Cat# 137612), Anti-mouse CD195 (CCR5)(Cat# 107011), Anti-mouse CD279 (PD-1) (Cat# 135224), Anti-mouse CD185 (CXCR5) (Cat# 145506), Anti-mouse IL-33Rα (ST2)(Cat# 146607), Anti-mouse CD193 (CCR3) (Cat# 144527), Anti-mouse CD4 (for T-regs quantification, Cat# 116023), Anti-mouse FoxP3 (Cat# 126403), Anti-mouse CD25 (Cat# 102037), Anti-mouse CD127 (IL-7Rα) (Cat# 135011), Anti-mouse Fc block CD16/32 (Cat# 101302), all from Biolegend. Anti-mouse H2-DM (Cat# 624048), Anti-mouse CD4 (Cat# 563232), Anti-mouse CD25 (Cat# 553866), Anti-mouse CD183 (CXCR3) (BDB755832), all from BD Biosciences. Anti-mouse CD74 (FAB7478A, R&D Systems), Anti-mouse Ly-6G/Gr1 (Cat# 17-9668-82, eBioscience).

All flow cytometry analyses, were performed on a BD LSR II (Becton Dickinson), equipped with a BD FACSDiva software v8.0.1. (BD Bioscience), except for Th1 cells which were analyzed in a BD FACSCelesta instrument. All subsequent compensation and gating were performed with FlowJo analysis software (BD LifeSciences). Gating strategies were set using the appropriate isotype control for each fluorochrome-conjugated staining antibody used. All isotype control antibodies were from BD Biosciences and used at 1:500 dilution.

### Protein isolation and western blotting

Mouse and human whole cell lysates were extracted with a buffer containing 1% NP40, 50 mM Tris pH 8.0, 150 mM NaCl, 10% glycerol, 2 mM EDTA and 10X concentration of Halt Protease and Phosphate Inhibitor Cocktail (ThermoFisher) for 30 mins on ice and spun at 13,000 rpm for 15 mins at 4°C. Protein samples were quantified using Protein Assay Dye Reagent (Bio-Rad). Prior to electrophoresis, samples were denatured in 5X laemmli buffer (8% SDS, 0.4 M DTT, 0.2 M Tris-HCl, 4.3 M glycerol, 6 mM bromophenol blue) and separated by sodium dodecyl sulfate-polyacrylamide gel electrophoresis (SDS-PAGE), transferred to immobilon-P polyvinylidene difluoride (PVDF) membrane (Millipore) at 100V for 1-2 h at 4°C. Membranes were blocked for 1h and probed with 5% nonfat dry milk with the following primary antibodies: anti-IRF8 (clone D20D8, Cat# 5628, Cell Signaling), anti-CD74 (sheep anti-mouse cd74, polyclonal antibody, Cat# AF7478, R&D system), anti-PU1 (clone 9G7, rabbit monoclonal, Cat# 2258, Cell Signaling), anti-FLAG (Cat# 8146, Cell Signaling), and anti-β-Actin (#A2228, Sigma-Aldrich). Membranes were developed with the SuperSignal West Pico PLUS Chemiluminescent Substrate (Thermo Fisher) and captured on X-Ray film. PVDF membranes were stripped with OneMinute Western Blot Stripping Buffer (#GM6001, GM Biosciences) and re-probed with relevant antibodies for loading control.

### In vitro cell growth assay

Models of human DLBCL cell lines (Toledo, SU-DHL4, SU-DHL6) and mouse B cell lymphoma cell lines (A20, 2PK-3 and BCL1), expressing an empty control vector, harboring IRF8 KO, or expressing IRF8 WT and mutants (S55A, N87Y, D400G and I424T), or with CD74 KO or its ectopic expression were analyzed for in vitro cell growth. In brief, cells were seeded at 5 x 10^5^/mL or 2.5 x 10^5^/mL in 6 well plates in triplicate, and quantified daily, for 96h, using an automated cell counter (Cellometer K2 Fluorescent Cell Counter, Nexcellom). Each assay was performed with three biological replicates.

### RNA isolation and Reverse Transcription Polymerase Chain Reaction (RT-PCR)

Cells were harvested and total RNA was isolated using Trizol (Invitrogen) as we described^86^. The RNA integrity and purity was determined by gel electrophoresis and by measuring 260/280 and 260/230 ratios in a nanodrop spectrophotometer, and cDNA generate with the High-Capacity cDNA Reverse Transcription Kit (Applied Biosystems). Q-RT-PCR were performed in triplicate using iTaq Universal SYBR Green (Bio-Rad) on QuantStudio 5 real-time PCR system (Applied Biosystems). All reactions included a no-reverse transcriptase sample and no-template control (water) to control for genomic DNA amplification or contamination. TBP was used as the internal control. Relative gene expression was calculated using the 2^-ΔΔ^Ct method, as we reported^87^. (Oligonucleotides sequences are listed in Supplemental Table 8).

### Estimation of immune-cell fractions based on gene expression

To estimate the relative abundance of cell types that might be present in the bulk RNAseq of the NCICCR-DLBCL tumor cohort, we used three computational tools based on enrichment or deconvolution of mixed sample population. First, we applied the xCell software, set to the default parameters, to generate enrichment scores for 64 immune and stromal cell types trained from a comprehensive database of reference signatures derived from over 1,800 pure cell types to predict the enrichment of these immune cell types in a mixed sample population^57^. Each cell type enrichment scores were then compared between IRF8 mutant or wild-type using two-sided Student’s t-test. We also applied CIBERSORTx, a method based on support-vector regression for the deconvolution of 22 immune cell types using default settings^62^. Additionally, we estimated the relative proportions of immune cell types employing the MCP-counter software, which is based on enrichment of marker genes and produces profiles of 11 cell types, including 8 immune cells^63^. Variations in the underlying methodology and reference cell types from these three algorithms provide complementary as well as supportive information.

### Structural modeling

For comparative structural analyses we used a crystal structure of IRF3 DBD/DNA complex (PDB: 2PI0) and AlphaFold model of IRF8 full-length (AF-Q02556-F1). IRF3/DNA complex structure comprises four IRF3 DBDs bound to a complete PRDIII-1 regulatory element of the human IFN-beta enhancer. We superposed the DBD of IRF8 over one of the four DNA-bound DBDs of IRF3 in Coot ^88^. The structure figures were prepared using Pymol (Schrodinger Suite).

### Quantification and statistical analysis

Analyses were performed using a one-way ANOVA, with Bonferroni’s multiple comparison or Fisher’s LSD post-hoc test, two-tailed Student’s t-test, and Mann-Whitney test. P<0.05 was considered significant. Data analyses were performed in the Prism 9 software (version 9.4.1, GraphPad Software Inc) and Excel (Microsoft 365 – Office).

## Acknowledgements

We thank Jamie Myers for technical help in the initial phase of this project. Y.K.G. is supported by funding from NIH/NIAID (R01AI161363) and the Welch Foundation (AQ-2101). P.L.M.D. is recipient of funds from the NIH (GM114102 and CA264248), the Neuroendocrine Tumor Research Foundation, the VHL Alliance, and is the holder of the Robert Tucker Hayes Distinguished Chair in Oncology. This work was also supported by funds from UT System Star Awards (to P.L.M.D.). R.C.T.A. acknowledges funding support from the Cancer Prevention and Research Institute of Texas (CPRIT, RP190043), the NIH (R01ES031522 and R01GM140456) and the Veterans Administration (I01BX001882).

## Author Contributions

Z.Q. and J.K. designed, performed, and interpreted assays. A-P.L., P.E., C.J., L.C., S.A. performed experiments. S.A. and Y.K.G. performed structural modeling. G.H-M. performed bioinformatics analysis. P.L.M.D. designed and supervised the bioinformatics analysis. R.C.T.A. conceived the project, designed, and interpreted the assays, and wrote the manuscript, which was reviewed by all authors.

## Figures Legend

**Supplemental Figure 1A.**
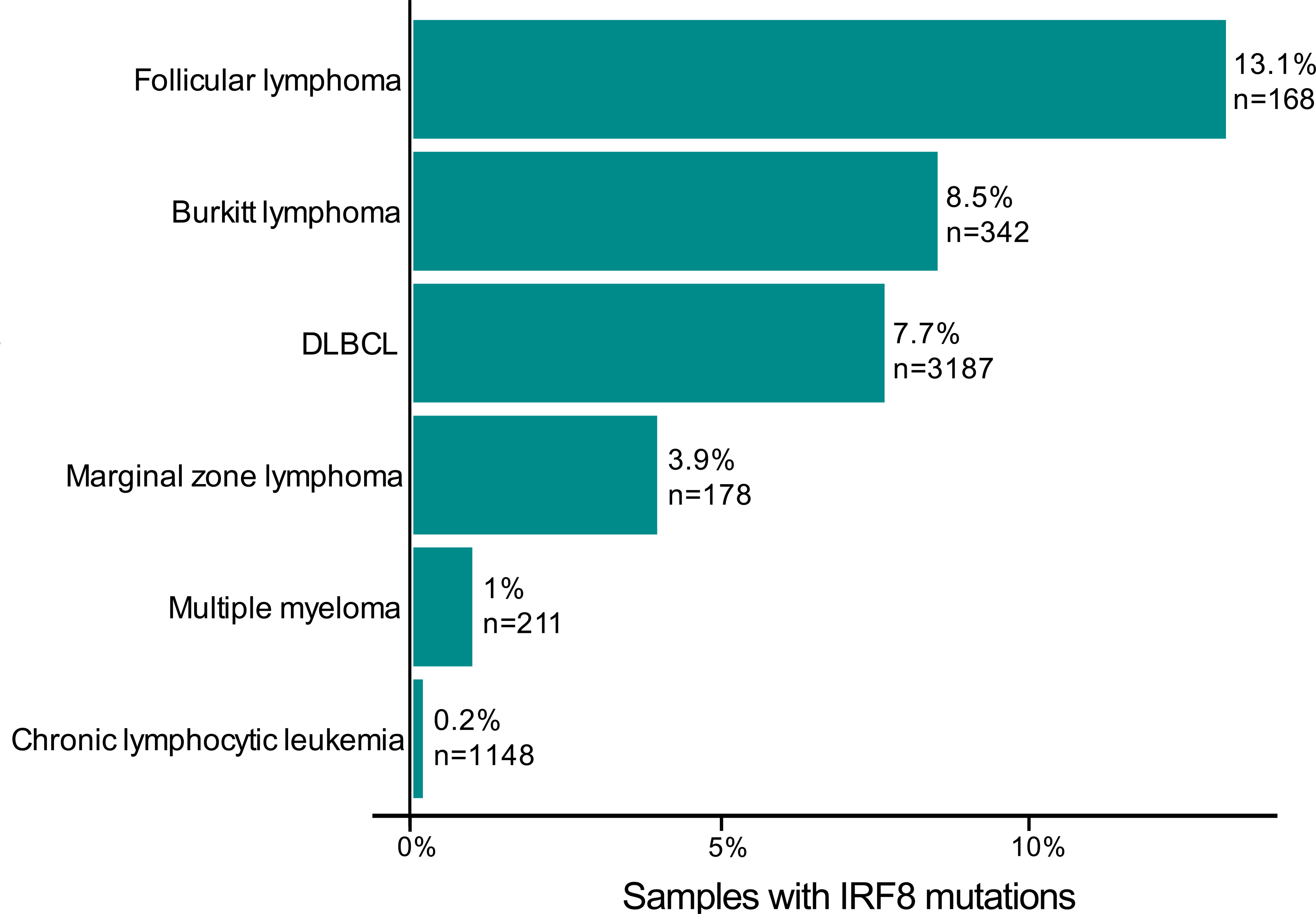
Frequency of *IRF8* variants in mature B-cell malignancies. Data are from publicly available cohorts, shown are size of cohort and % of *IRF8* gene mutations (related to Supplemental Tables 1, 5 and 6).

**Supplemental Figure 2A.**
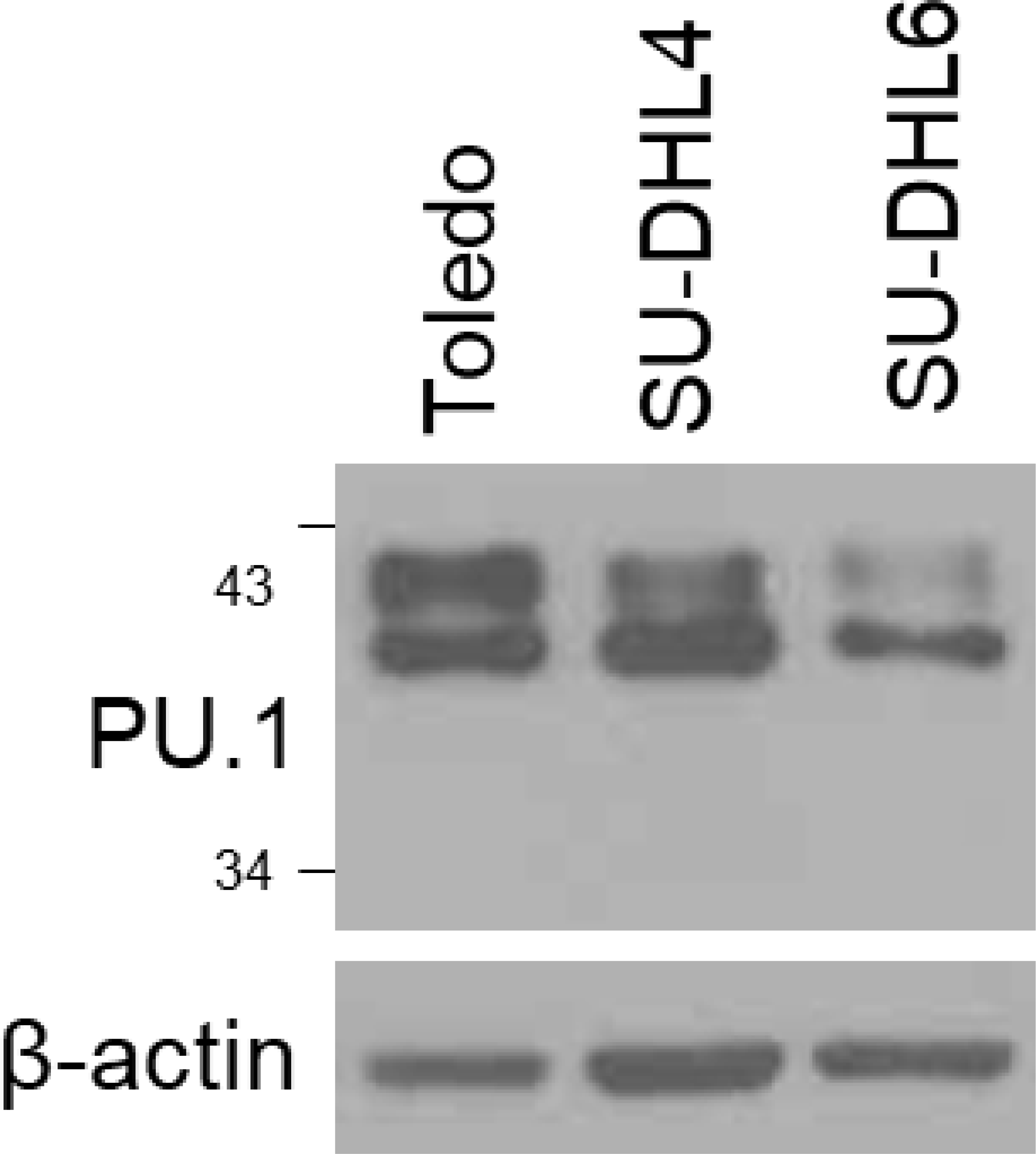
PU.1 expression. Western blot analysis of PU.1 expression in human DLBCL.

**Supplemental Figure 3A.**
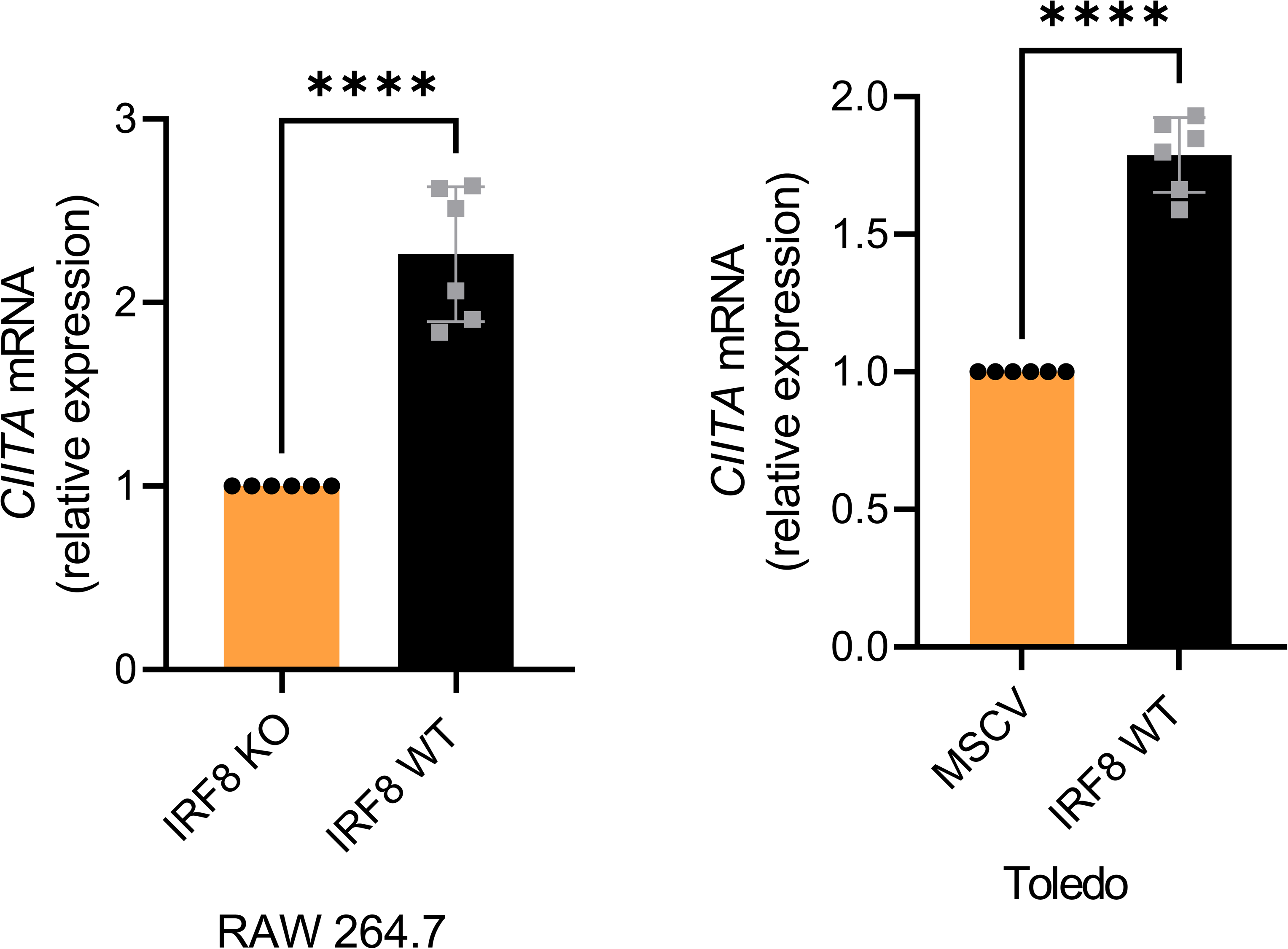
*CIITA* q-RT-PCR. *CIITA* mRNA expression in cell lines with IRF8 KO (RAW 264.7, left) or lacking endogenous IRF8 expression (Toledo, right), followed by stable ectopic expression of human IRF8 WT. Data are mean ± SD of a total of four biological replicates; P values are from two-sided Student’s t-test (**** = <0.0001).

**Supplemental Figure 3B.**
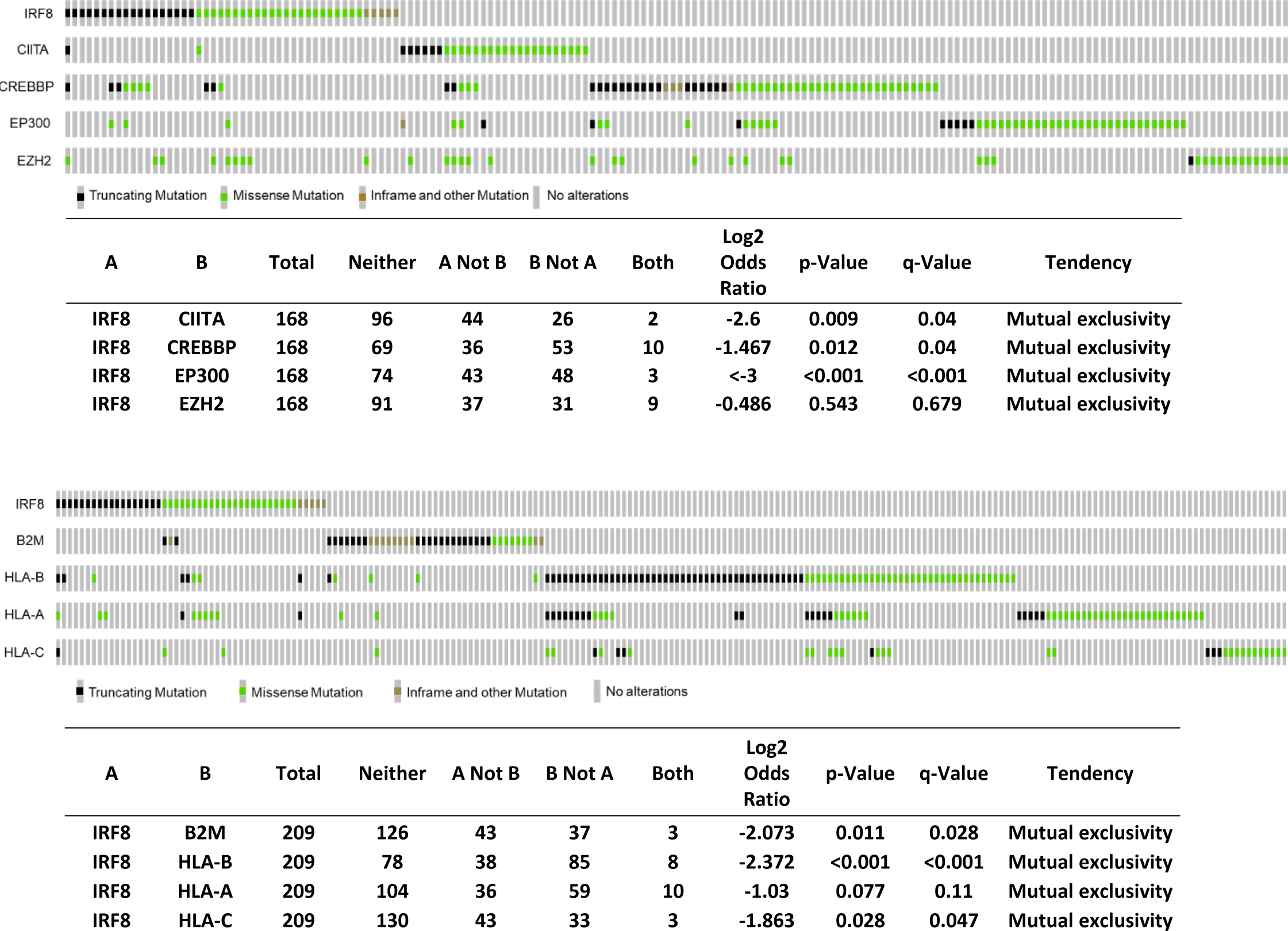
IRF8 mutation mutual exclusivity in DLBLC. **Top** - Oncoprint display of comparative distribution of IRF8, CIITA, CREBBP, EP300 and EZH2 gene mutations in 168 DLBCLs; pairwise log2 odds score, as well as P and Q values of the correlation between the co-occurrence (positive score) or mutually exclusive (negative score) presence of mutation, are shown in a table format. **Bottom** - Oncoprint display of comparative distribution of IRF8, B2M, HLA-A, HLA-B and HLA-C gene mutations in 209 DLBCLs; pairwise log2 odds score, as well as P and Q values of the correlation between the co-occurrence (positive score) or mutually exclusive (negative score) presence of mutation, are also shown in table format. Symbols are color coded based on the type of mutation. q value <0.05 was considered significant.

**Supplemental Figure 4A.**
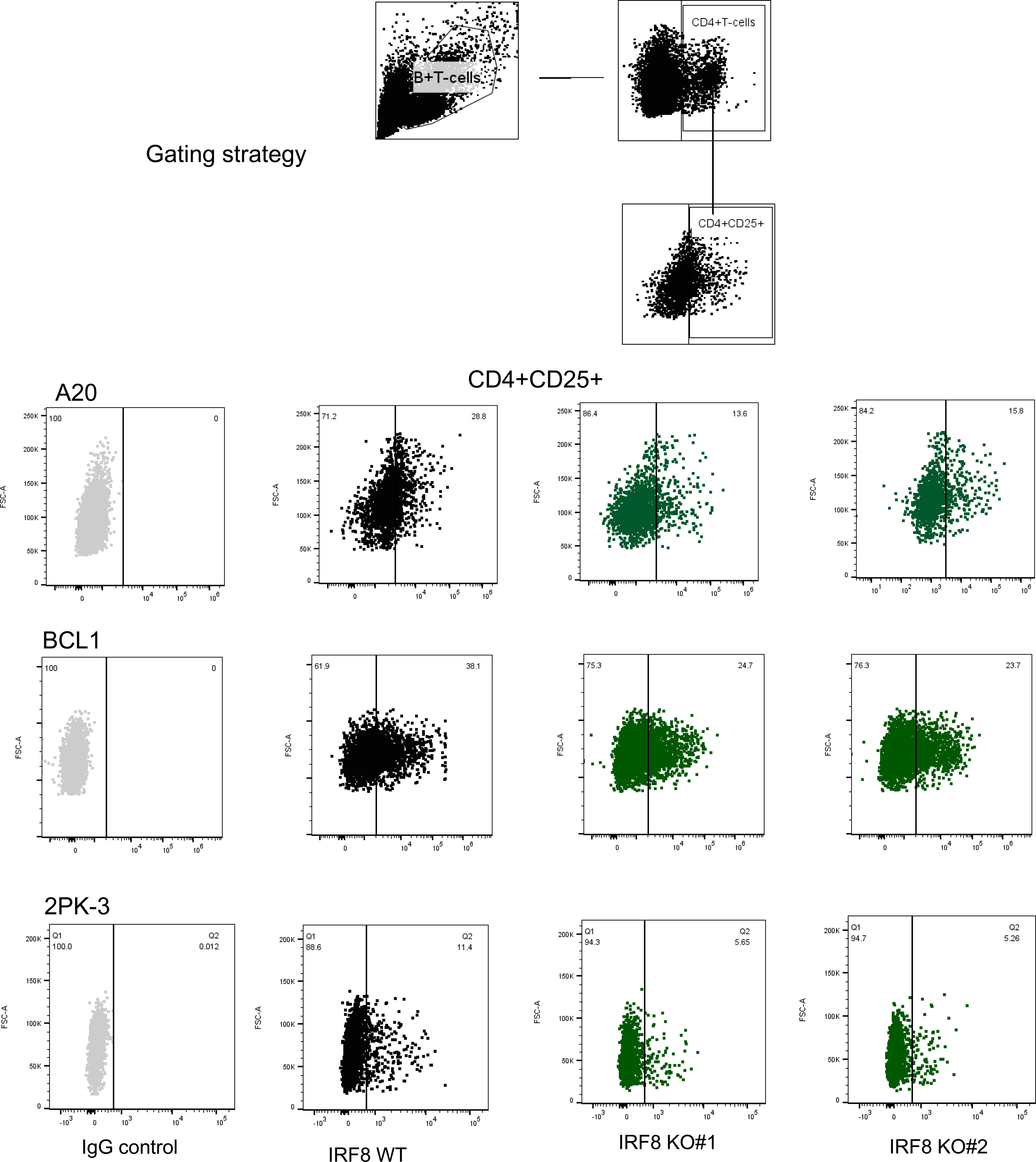
FACS analysis of CD25. Gating strategy and representative displays of CD25 quantification by FACS in CD4+ DO-11.10 murine cells following antigen (OVA) presentation by three B cell lymphoma cells (A20, BCL1, 2PK-3), WT or KO for IRF8 (2 independent KO clones).

**Supplemental Figure 4B.**
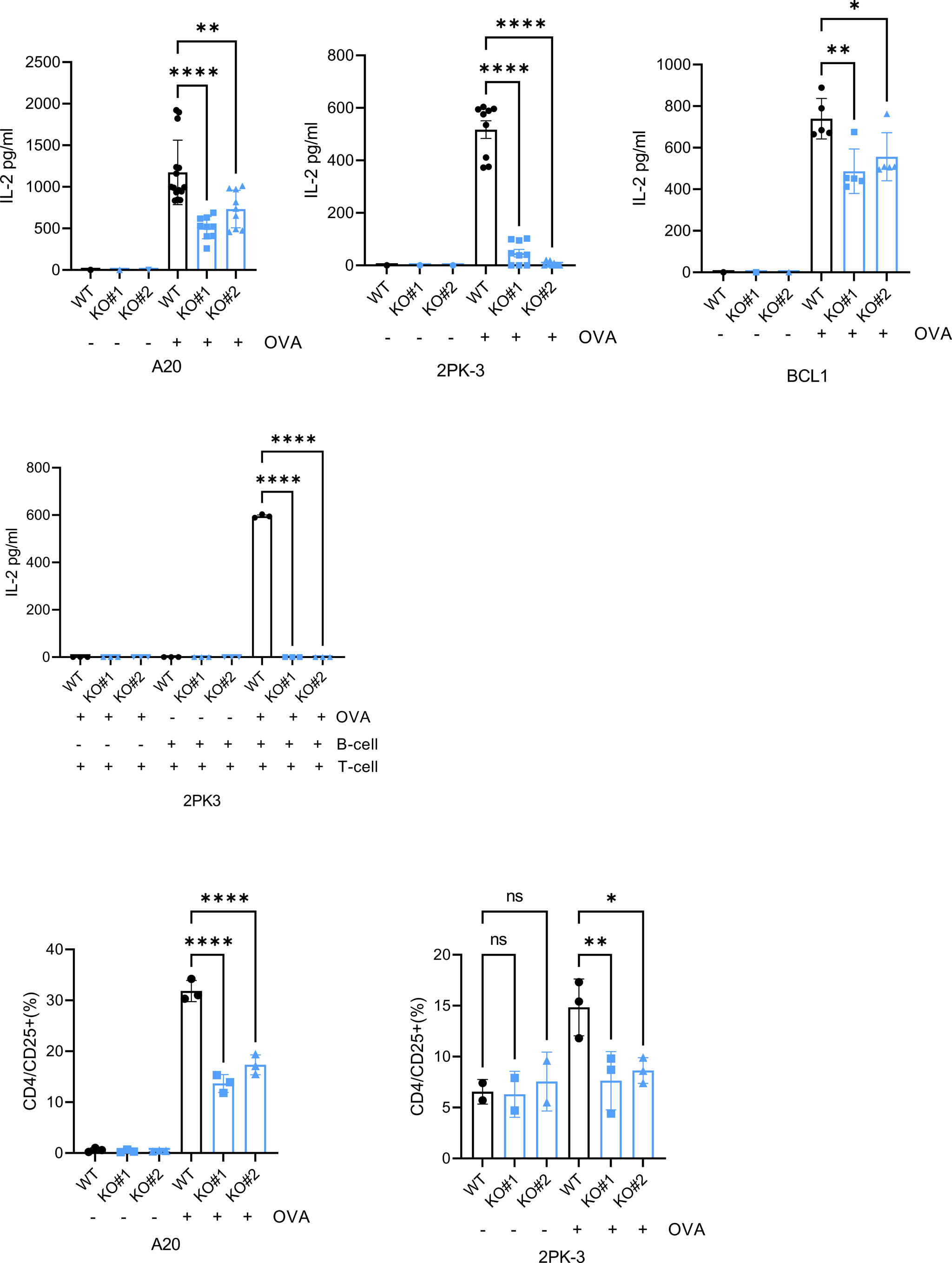
Activation of CD4+ DO-11.10 cells. **Top panels.** ELISA-based IL-2 quantification in conditioned media of CD4+ DO-11.10 cells co-cultured with mouse B cell lymphoma cell lines A20, BCL1 and 2PK-3, each IRF8 WT or KO, “loaded” or not with OVA. **Middle panel**. ELISA-based IL-2 quantification in conditioned media of CD4+ DO-11.10 cells exposed to IL-2 but not co-cultured with 2PK-3 lymphoma cells, or in conditioned media of CD4+ DO-11.10 cells co-cultured with the mouse B cell lymphoma cell line 2PK-3, IRF8 WT or KO, “loaded” or not with OVA. **Bottom panels.** CD25 quantification by FACS in CD4+ DO-11.10 murine cells co-cultured with mouse B cell lymphoma cell lines A20 and 2PK-3, each IRF8 WT or KO, “loaded” or not with OVA. Data are mean ±SD of two to three biological replicates, performed with one, two or three technical replicates. P values are from one ANOVA with Bonferroni post-test;, *(p<0.05), **(p<0.01), ***(p<0.001), **** (p<0.0001).

**Supplemental Figure 4C.**
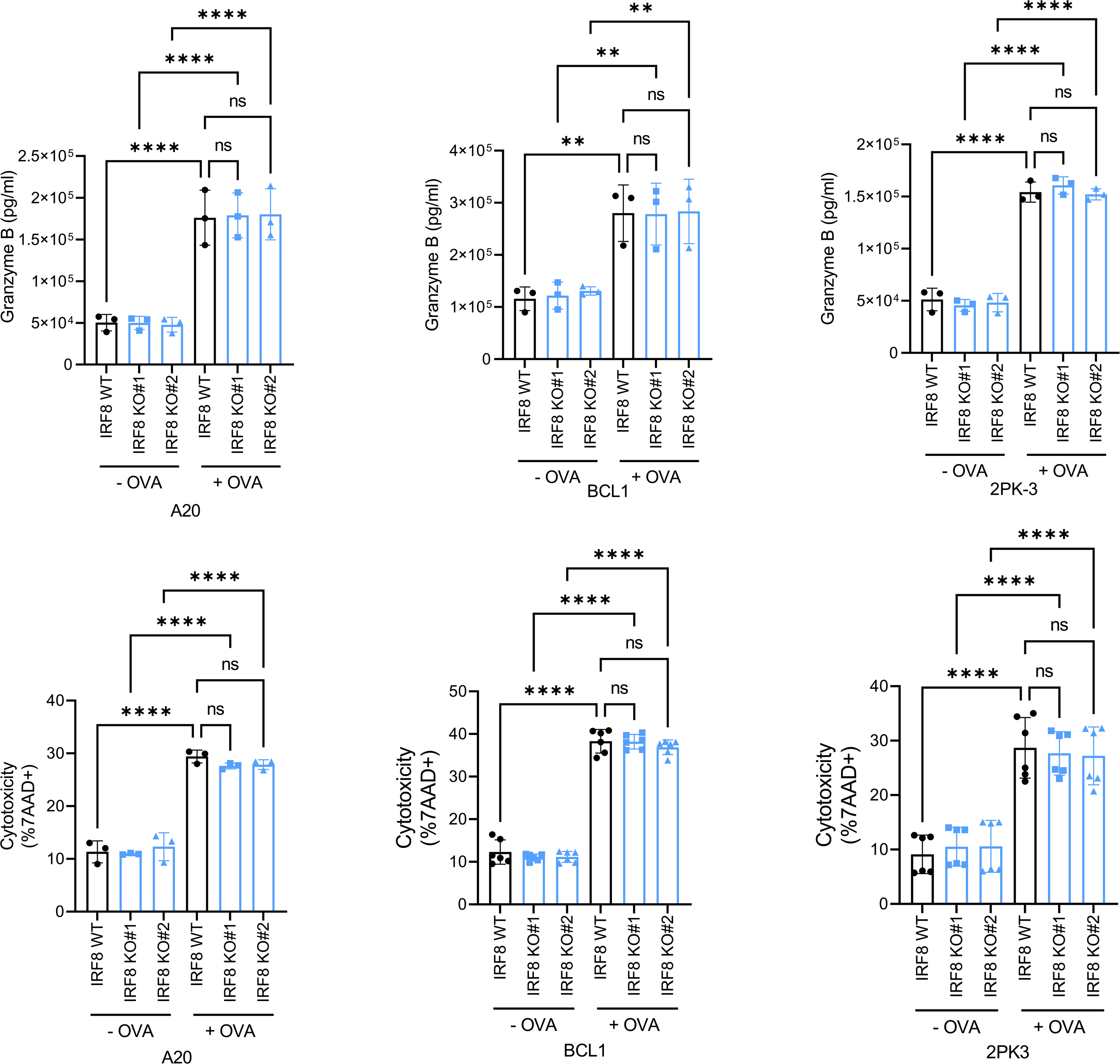
Activation of CD8+ OT-I cells Top panels. ELISA-based granzyme quantification in conditioned media of CD8 OT-I cells co-cultured with mouse B cell lymphoma cell lines A20, BCL1 and 2PK-3, each IRF8 WT or KO, “loaded” or not with OVA. **Bottom panels**. FACS-based quantification of 7AAD+ lymphoma cells (A20, BCL1, or 2PK-3), IRF8 WT or KO, “loaded” or not with OVA. Data are mean ±SD of three biological replicates, performed with one, two or three technical replicates. P values are from one-way ANOVA with Bonferroni post-test;, *(p<0.05), **(p<0.01), ***(p<0.001), **** (p<0.0001).

**Supplemental Figure 4D.**
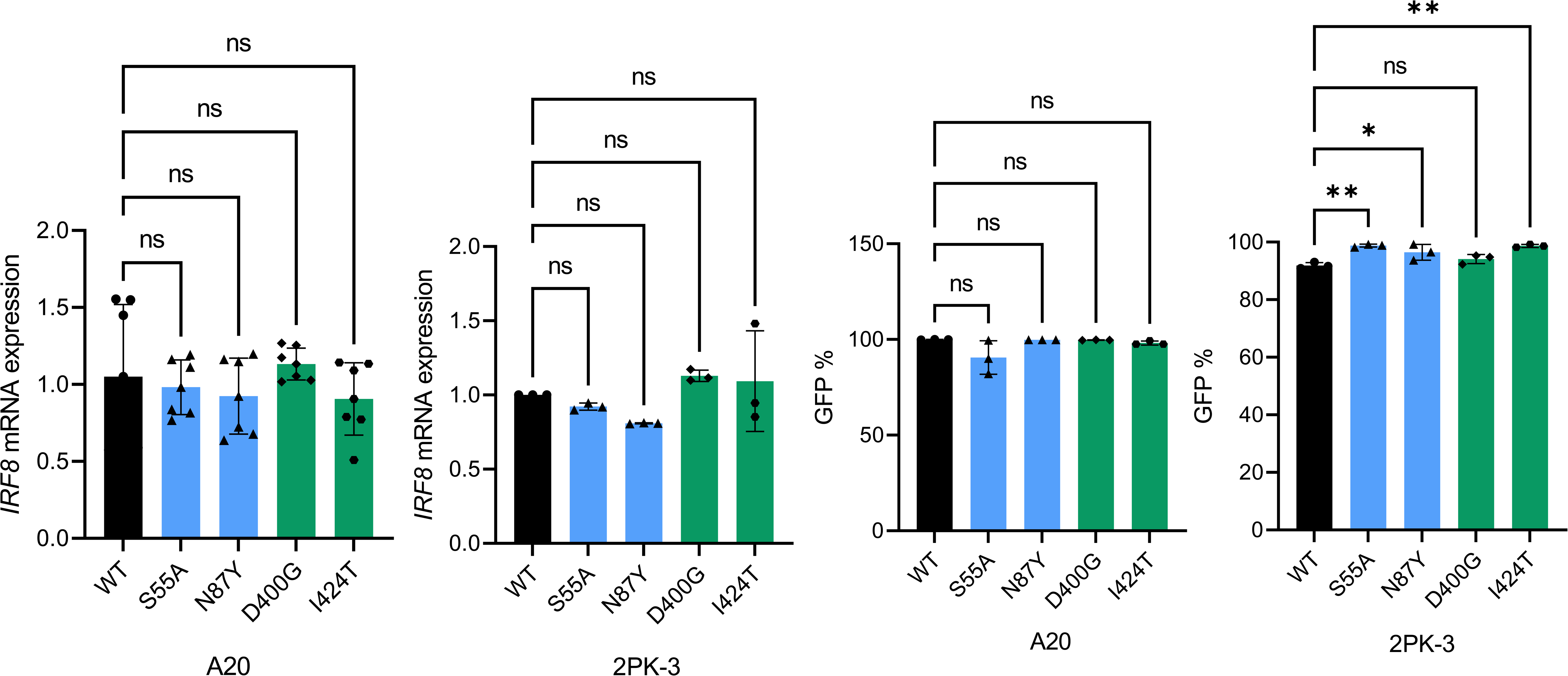
Characterization of IRF8 WT or mutant mouse B cell lymphoma cell lines. **Left to right**. Q-RT-PCR-based quantification of IRF8 mRNA expression in mouse B cell lymphoma cell lines A20 and 2PK-3 stably expressing human IRF8 WT or mutant in a retrovirus vector bicistronic for GFP (MSCV-eGFP) – GFP%, defined by FACS, is shown to the right. Data are mean ±SD of three biological replicates, performed with one or two technical replicates. P values are from one-way ANOVA with Bonferroni post-test;, *(p<0.05), **(p<0.01).

**Supplemental Figure 4E.**
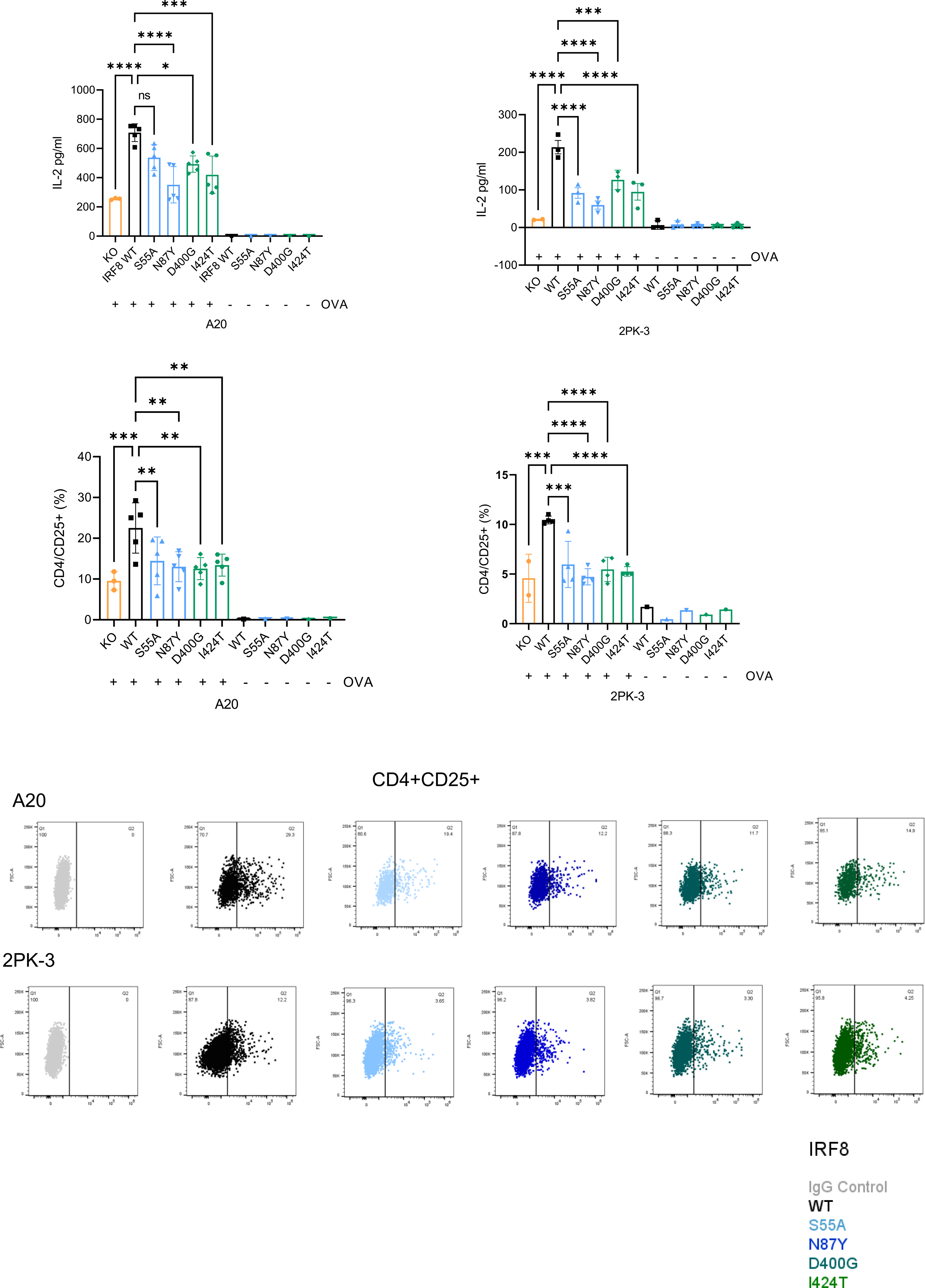
Activation of CD4+ DO-11.10 cells. **Top panels.** ELISA-based IL-2 quantification in conditioned media of CD4+ DO-11.10 cells co-cultured with mouse B cell lymphoma cell lines A20 (left) or 2PK-3 (right), each expressing IRF8 WT or mutant, “loaded” or not with OVA. **Middle panels.** CD25 quantification by FACS in CD4+ DO-11.10 murine cells co-cultured with mouse B cell lymphoma cell lines A20 (left) or 2PK-3 (right), each expressing IRF8 WT or mutant, “loaded” or not with OVA. Data are mean ±SD of three biological replicates, performed with one, two or three technical replicates. P values are from one ANOVA with Bonferroni post-test; *(p<0.05), **(p<0.01), ***(p<0.001), **** (p<0.0001). **Bottom panels.** Representative displays of CD25 quantification by FACS in CD4+ DO-11.10 murine cells following antigen (OVA) presentation by the B cell lymphoma cells A20 and 2PK-3, expressing IRF8 WT or the mutants S55A, N87Y, D400G and I424T.

**Supplemental Figure 5A.**
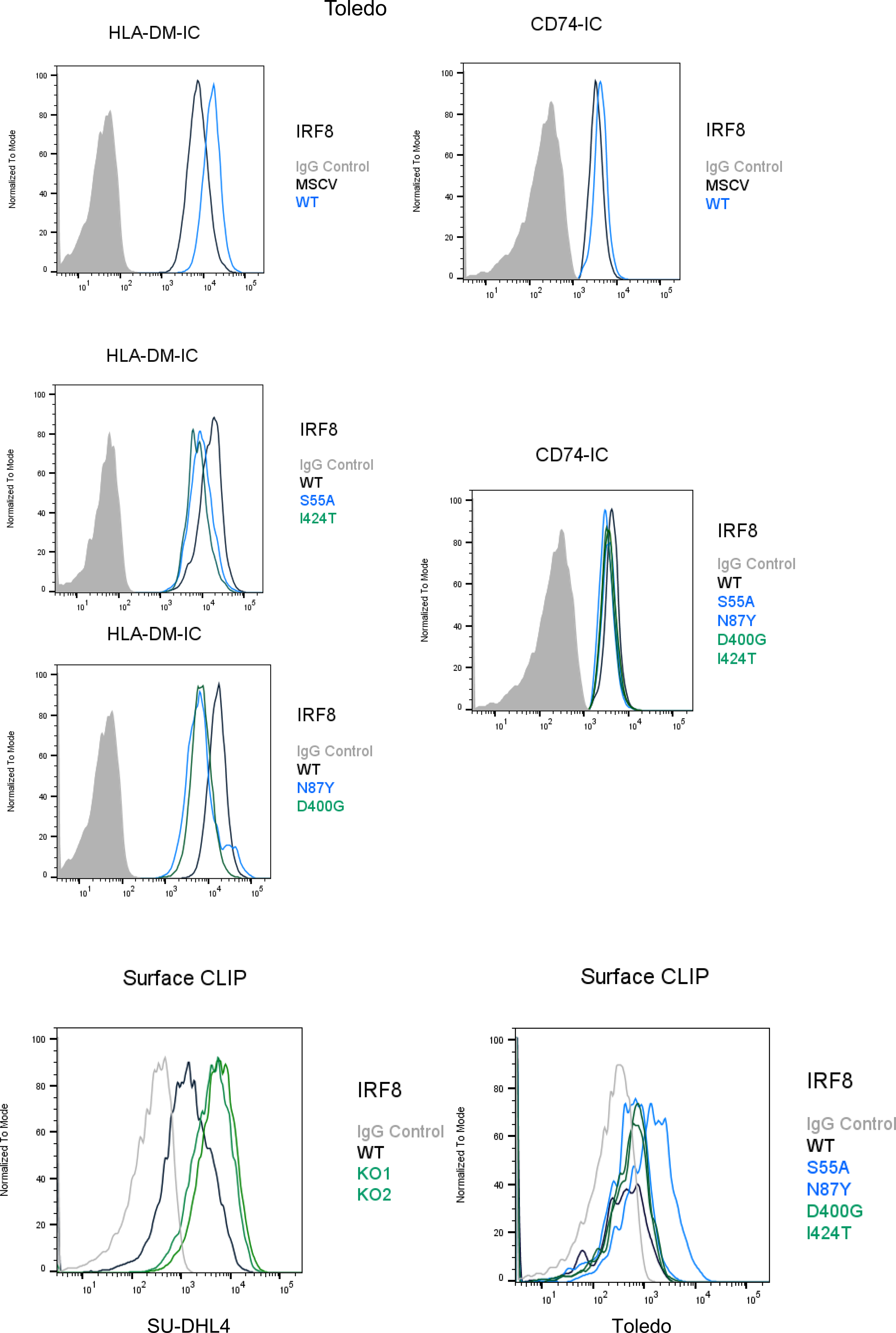
FACS analysis of DLBLC cell lines. **Top.** Representative histograms of FACS- based intra-cellular quantification of HLA-DM (left) or CD74 (right) in the DLBCL cell line Toledo expressing an empty vector (MSCV), IRF8 WT. **Middle.** Representative histograms of FACS-based intra-cellular quantification of HLA-DM (left) or CD74 (right) in the DLBCL cell line Toledo expressing IRF8 WT or IRF8 S55A, N87Y, D400G, I424T. **Bottom**. Representative histograms of FACS-based quantification of cell-surface CLIP in IRF8 WT or KO SU-DHL4 cell line (left). or in Toledo cells expressing IRF8 WT, S55A, N87Y, D400G, I424T (right).

**Supplemental Figure 5B.**
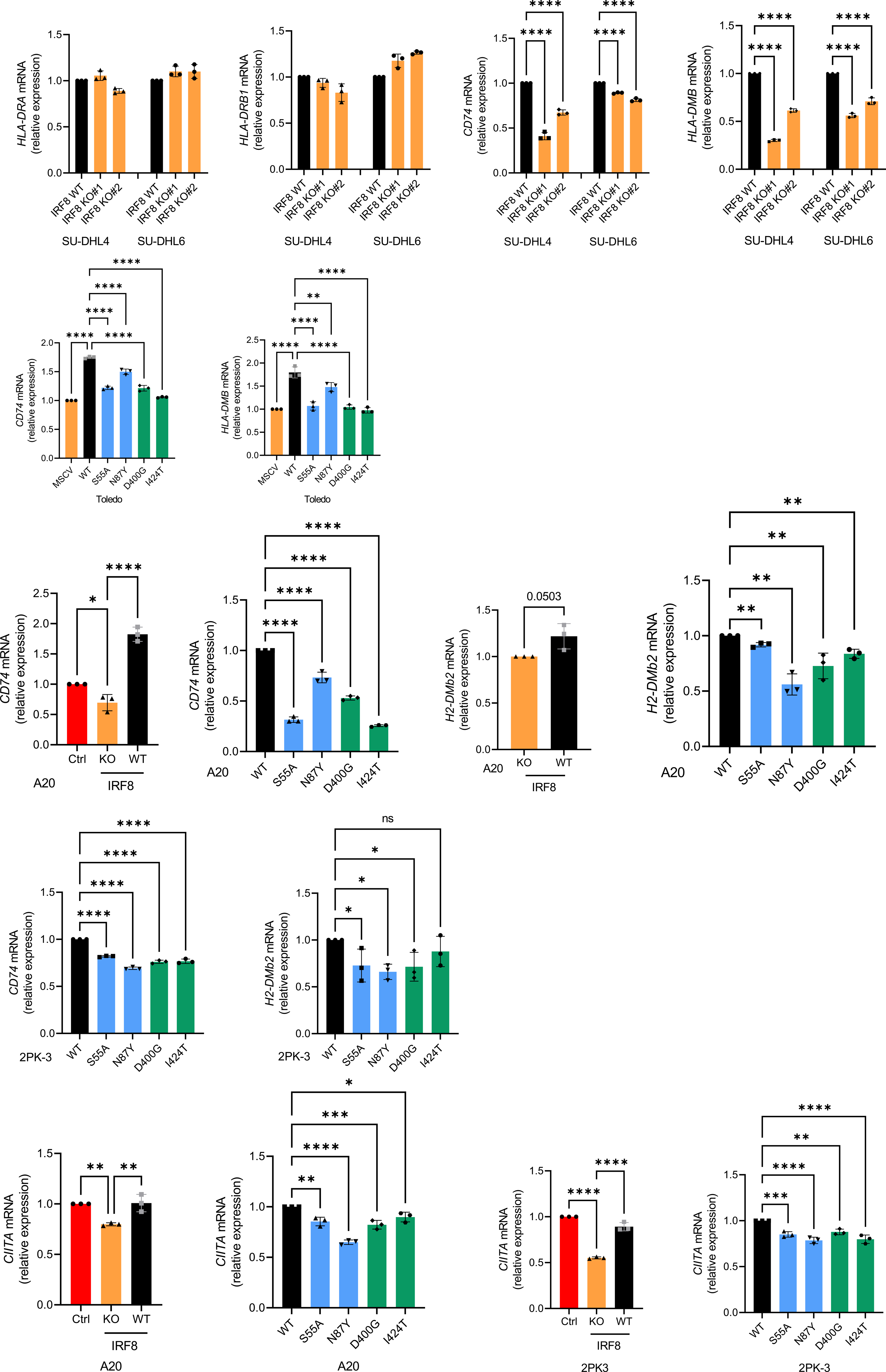
Q-RT-PCRs in IRF8 WT, KO or mutant DLBCL and mouse B cell lymphoma cell lines. **Top to bottom.** mRNA expression of *HLA-DRA*, *HLA-DRB1*, *CD74* and *HLA-DMB* in IRF8 WT or KO SU-DHL4 and SU-DHL6 cell lines. mRNA expression of *CD74* and *HLA-DMB* in Toledo cell line expressing an empty vector (MSCV), IRF8 WT or S55A, N87Y, D400G, I424T mutants. mRNA expression of *CD74* and *H2-DMb2* in A20 mouse B cell lymphoma cell line expressing an empty vector (ctrl), IRF8 KO, or IRF8 WT, S55A, N87Y, D400G, I424T mutants. mRNA expression of *CD74* and *H2-DMb2* in 2PK-3 mouse B cell lymphoma cell line expressing IRF8 WT, or mutants S55A, N87Y, D400G, I424T. mRNA expression of *CIITA* in A20 and 2PK-3 in mouse B cell lymphoma cell lines expressing an empty vector (ctrl), IRF8 KO, or IRF8 WT, S55A, N87Y, D400G, I424T mutants. Data are mean ±SD of three biological replicates. P values are from one-way ANOVA with Bonferroni or Fisher’s LSD post-test, or from two-tailed Student’s t-test. *(p<0.05), **(p<0.01), ***(p<0.001), **** (p<0.0001).

**Supplemental Figure 5C.**
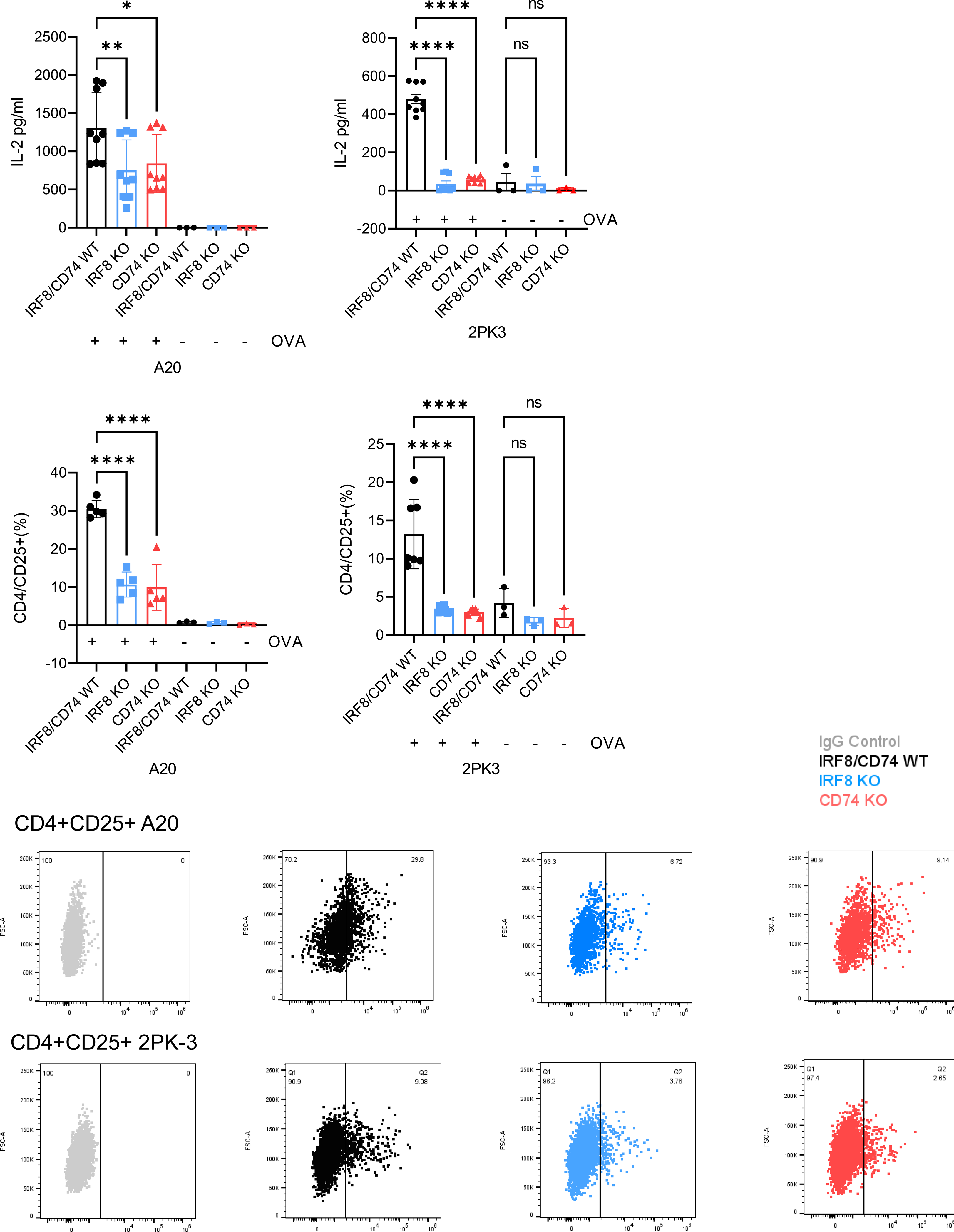
Activation of CD4+ DO-11.10 cells. **Top panels.** ELISA-based IL-2 quantification in conditioned media of CD4+ DO-11.10 cells co-cultured with mouse B cell lymphoma cell lines A20 (left) or 2PK-3 (right), each expressing IRF8/CD74 WT or each individual KO, “loaded” or not with OVA. **Middle panels.** CD25 quantification by FACS in CD4+ DO-11.10 murine cells co-cultured with mouse B cell lymphoma cell lines A20 (left) or 2PK-3 (right), each expressing IRF8/CD74 WT or each individual KO, “loaded” or not with OVA. Data are mean ±SD of three biological replicates, performed with one, two or three technical replicates. P values are from one ANOVA with Bonferroni or Fisher’s LSD post-test; *(p<0.05), **(p<0.01), ***(p<0.001), **** (p<0.0001). **Bottom panels.** Representative displays of CD25 quantification by FACS in CD4+ DO-11.10 murine cells following antigen (OVA) presentation by the B cell lymphoma cells A20 and 2PK-3, expressing IRF8/CD74 WT or each individual KO.

**Supplemental Figure 5D.**
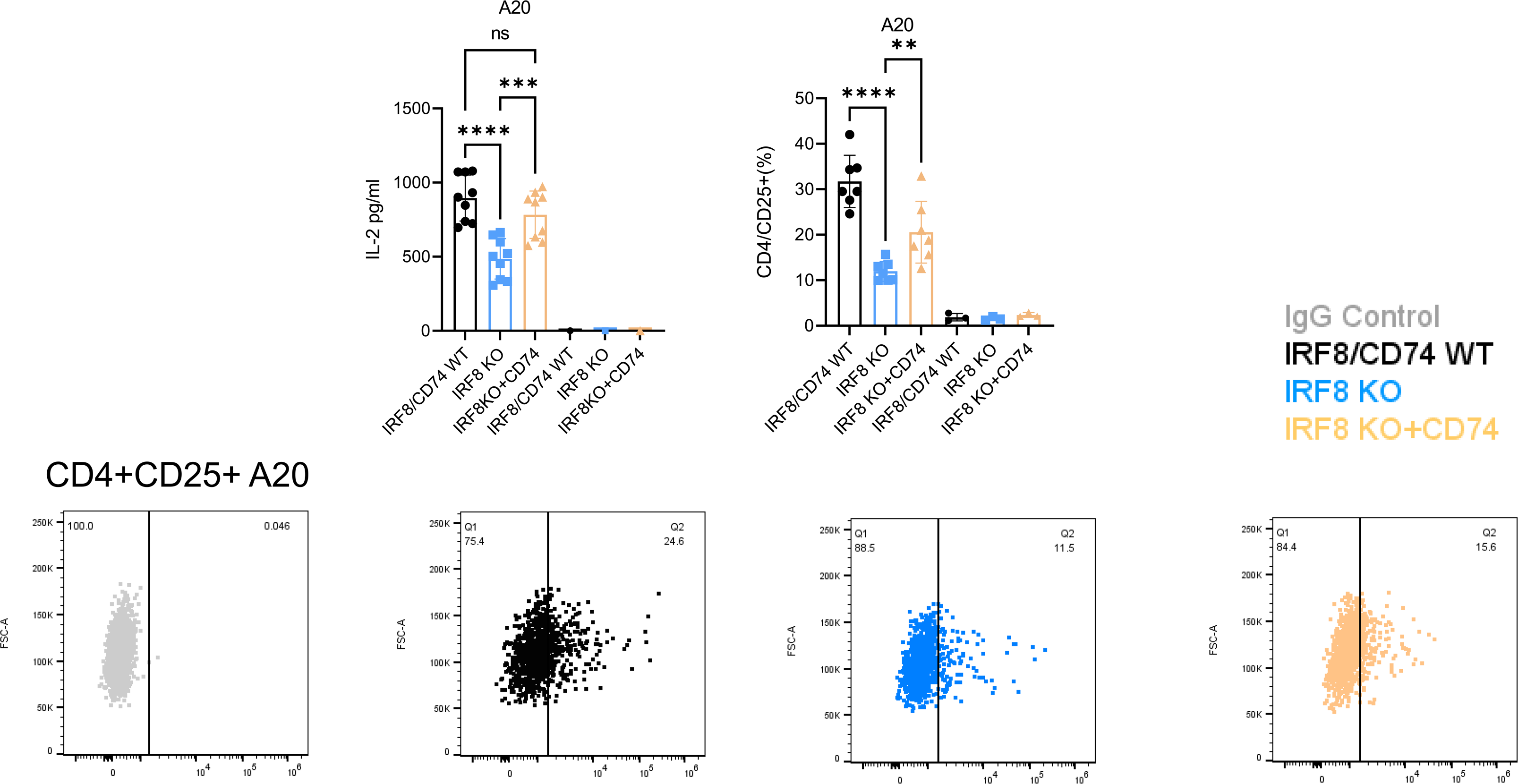
Activation of CD4+ DO-11.10 cells. **Top left bar-graph.** ELISA-based IL-2 quantification in conditioned media of CD4+ DO-11.10 cells co-cultured with the mouse B cell lymphoma cell line A20 expressing IRF8/CD74 WT, IRF8 KO, or IRF8 KO + ectopically expressed CD74, “loaded” or not with OVA. **Top right bar-graph.** CD25 quantification by FACS CD4+ DO-11.10 cells co-cultured with the mouse B cell lymphoma cell line A20 expressing IRF8/CD74 WT, IRF8 KO, or IRF8 KO + ectopically expressed CD74, “loaded” or not with OVA. Data are mean ±SD of three biological replicates, performed with one or three technical replicates. P values are from one-way ANOVA with Bonferroni post-test; **(p<0.01), ***(p<0.001), **** (p<0.0001). **Bottom panels.** Representative displays of CD25 quantification by FACS in CD4+ DO-11.10 murine cells following antigen (OVA) presentation by the B cell lymphoma cell line A20 expressing IRF8/CD74 WT, IRF8 KO or or IRF8 KO + ectopically expressed CD74.

**Supplemental Figure 5E.**
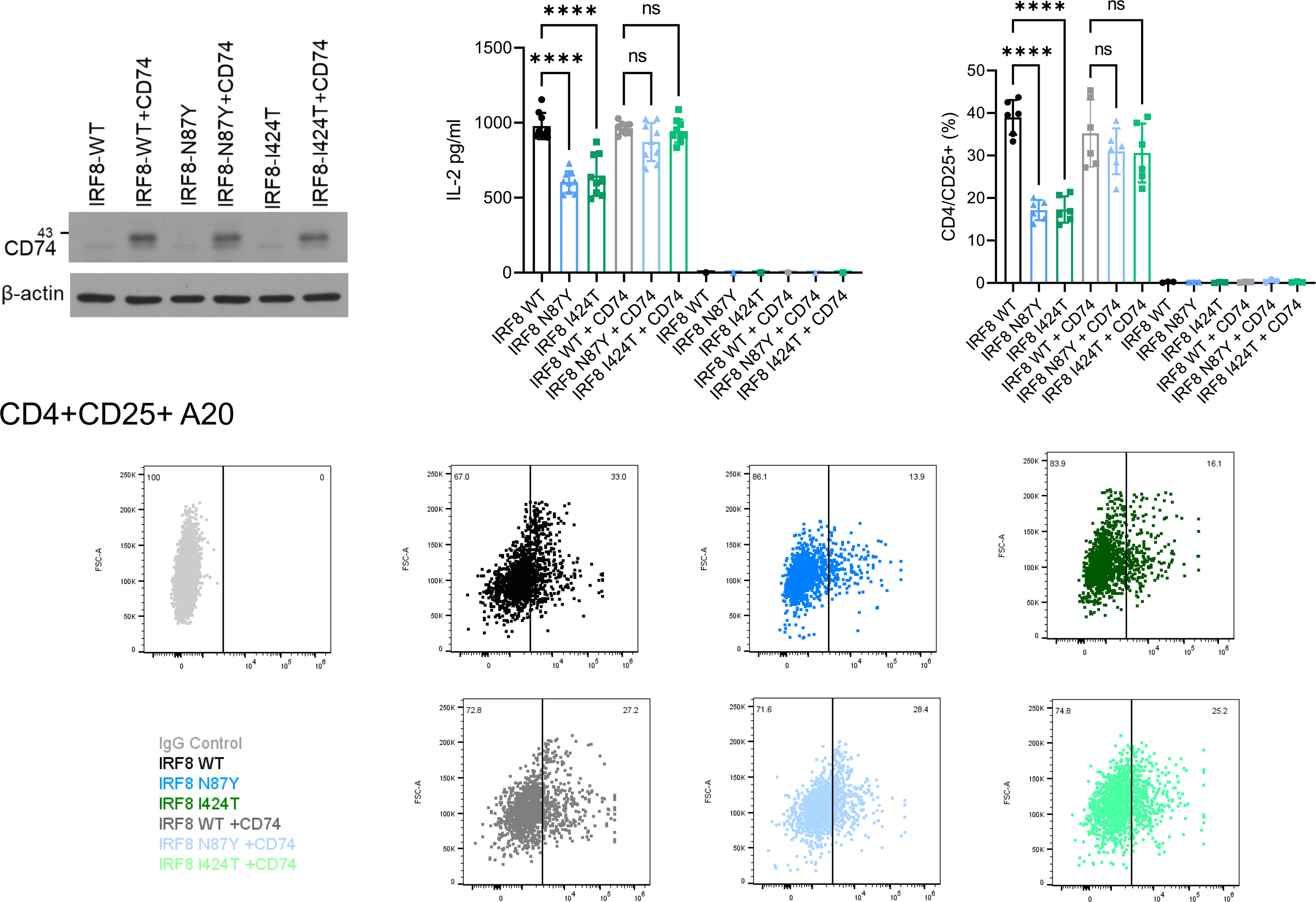
Activation of CD4+ DO-11.10 cells. **Top, left to right.** CD74 WB in A20 models expressing IRF8 WT, N87Y or I424T -/+ CD74. Bar-graphs; **left**, ELISA-based IL-2 quantification in conditioned media of CD4+ DO-11.10 cells co-cultured with the mouse B cell lymphoma cell line A20 expressing IRF8 WT, N87Y or I424T -/+ ectopically expressed CD74, “loaded” or not with OVA, **right,** CD25 quantification by FACS in CD4+ DO-11.10 cells co-cultured with the mouse B cell lymphoma cell line A20 expressing IRF8 WT, N87Y or I424T -/+ ectopically expressed CD74, “loaded” or not with OVA. Data are mean ±SD of three biological replicates, performed with one, two or three technical replicates. P values are from one-way ANOVA with Bonferroni post-test; **** (p<0.0001). **Bottom panels.** Representative displays of CD25 quantification by FACS in CD4+ DO-11.10 murine cells following antigen (OVA) presentation by the B cell lymphoma cell line A20 expressing IRF8 WT, N87Y or I424T -/+ ectopically expressed CD74.

**Supplemental Figure 6A.**
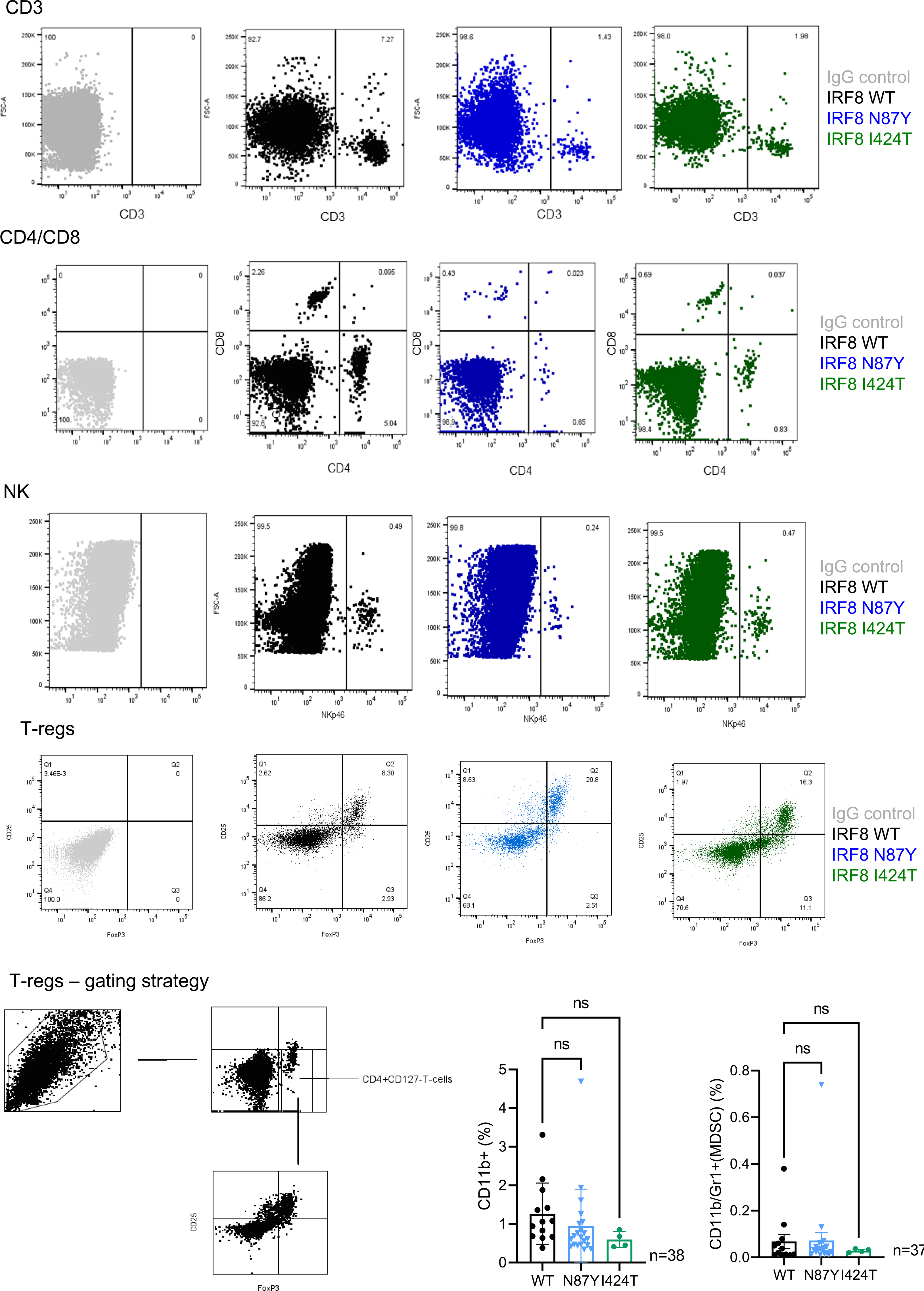
FACS analysis of lymphoma microenvironment. **Top to bottom.** Representative displays of CD3, CD4/CD8, NK, T-regs measurements in the TME of lymphomas expressing IRF8 WT, N87Y or I424T. Gating strategy for T-regs is also. Bar-graphs at the bottom right show the FACS-based quantification of monocytes and MDSC in the TME of lymphomas expressing IRF8 WT, N87Y or I424T. Data shown are mean ± SD of 38 and 37 mice, respectively. Statistical significance was tested with one-way ANOVA and Bonferroni post-test.

**Supplemental Figure 6B.**
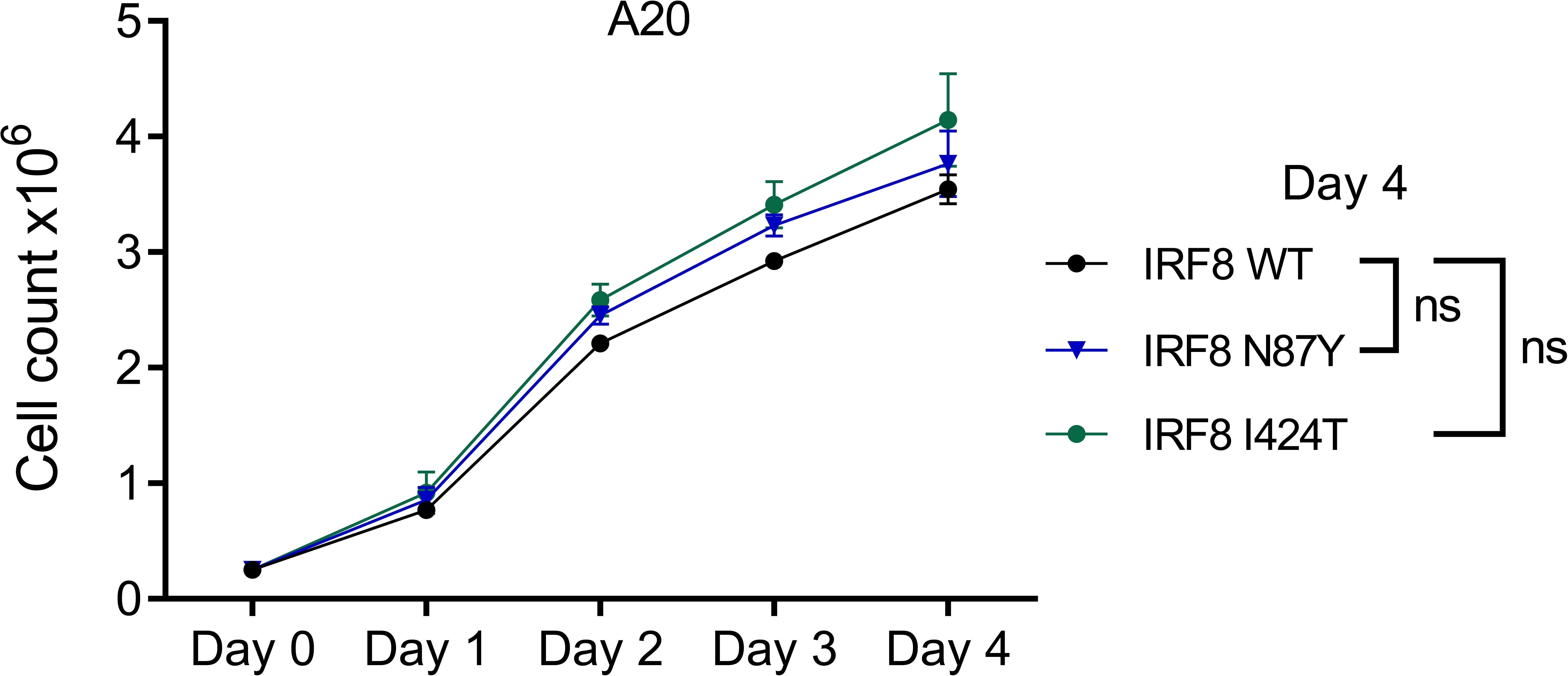
In vitro growth curve of B cell lymphoma cell line A20 expressing IRF8 WT, N87Y or I424T mutants, determined with automated fluorescent cell counter. Data are mean ± SD of three biological replicates. P value (non-significant, ns) is from two-sided Student’s t-test, calculated in WT vs. each mutant, at the day 4 time point.

**Supplemental Figure 6C.**
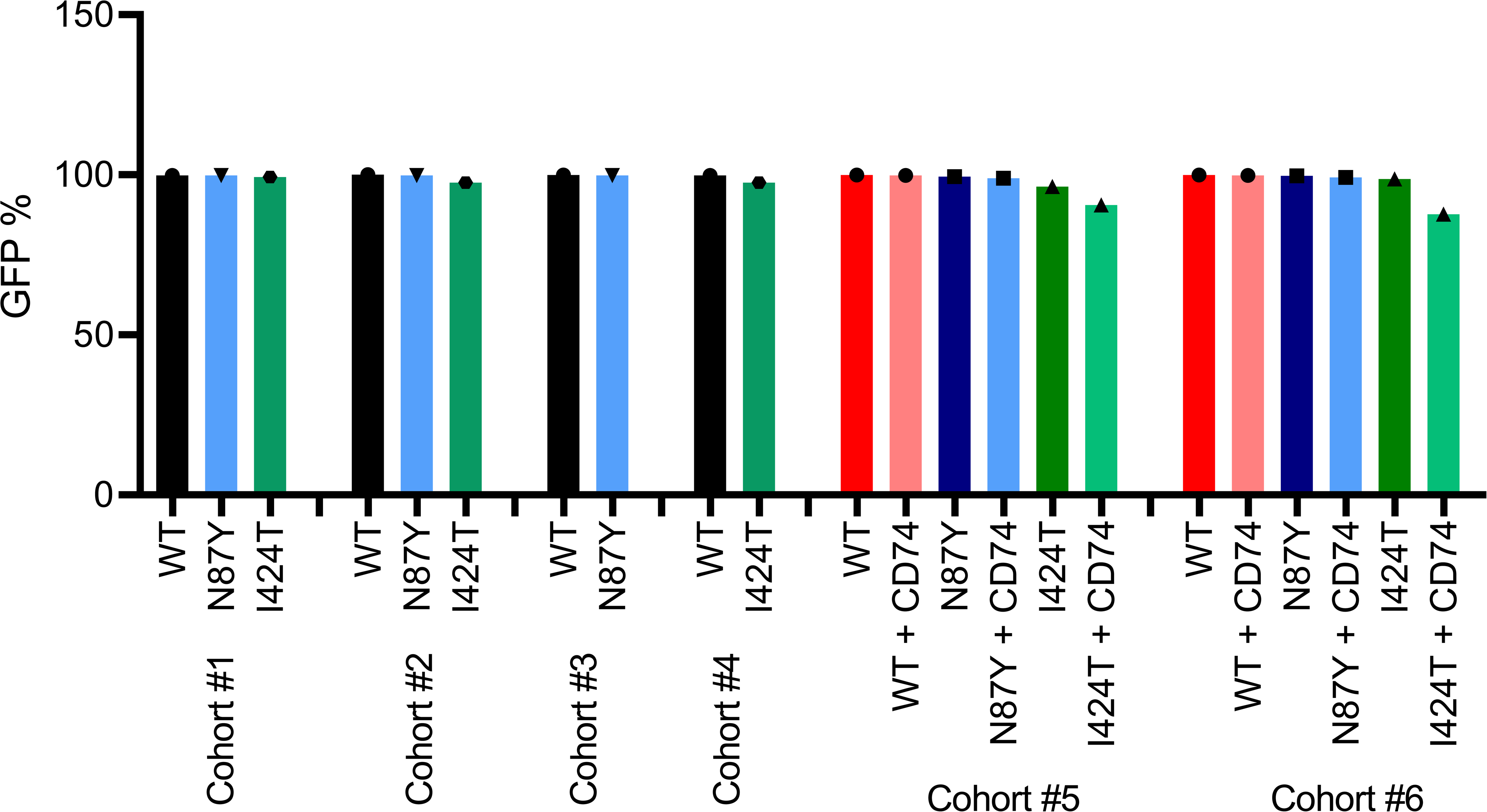
GFP quantification. (% of +cells) determined by FACS in A20 cells inject in BALB/c mice in each of the six cohorts analyzed.

**Supplemental Figure 6D.**
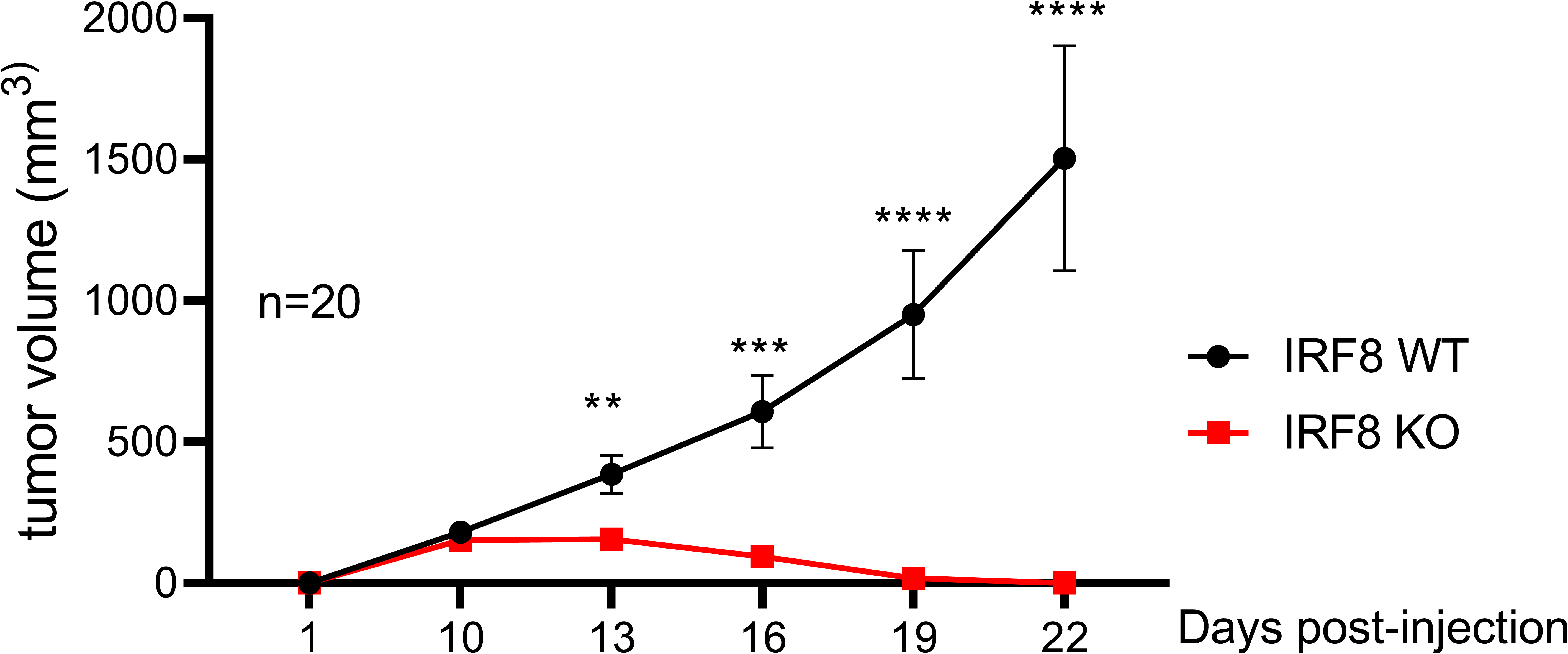
In vivo growth curve (tumor volume) of IRF8 WT or KO A20 lymphoma cells injected in BALB/c mice. Data are mean ± SEM of two independent cohorts; p values are from two-sided Student’s t-test. **(p<0.01), ***(p<0.001), **** (p<0.0001).

**Supplemental Figure 6E.**
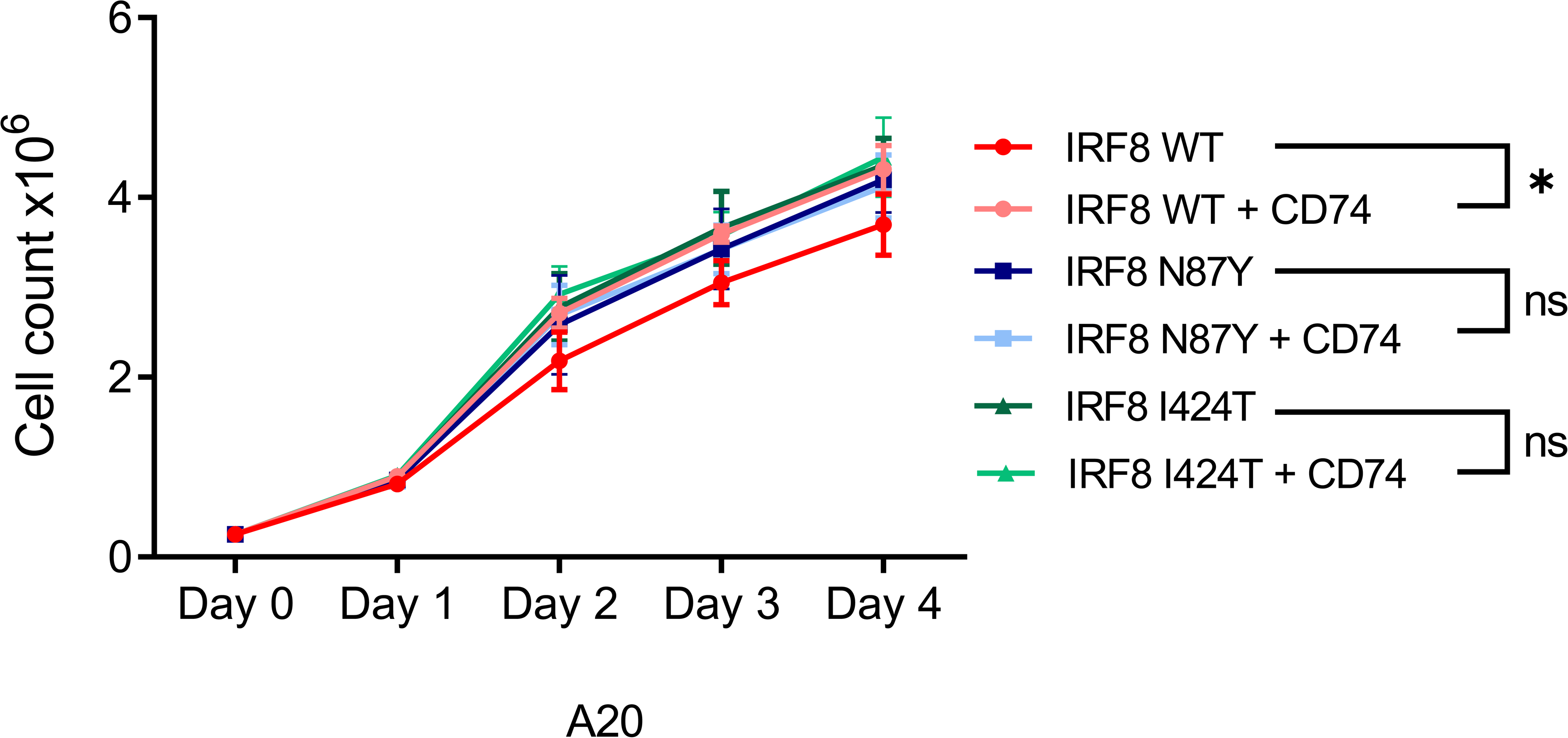
In vitro growth curve of B cell lymphoma cell line A20 expressing IRF8 WT, N87Y or I424T mutants, -/+ CD74 stable ectopic expression, determined with automated fluorescent cell counter. Data are mean ± SD of three biological replicates. P values are from two-sided Student’s t-test, calculated at the day 4 time point.

**Supplemental Figure 6F.**
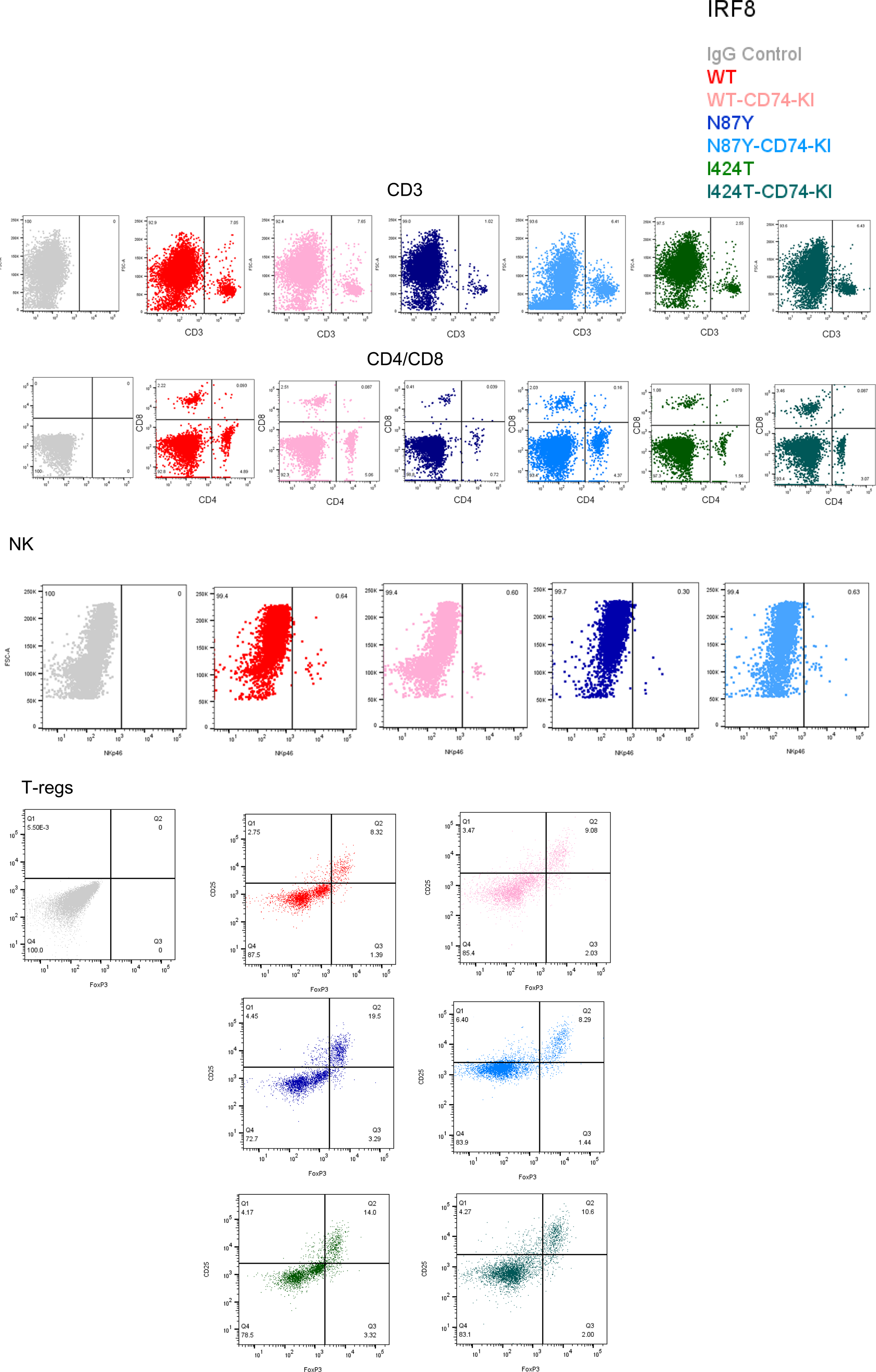
FACS analysis of lymphoma microenvironment. **Top to bottom.** Representative displays of CD3, CD4/CD8, NK, T-regs measurements in the TME of lymphomas expressing IRF8 WT, N87Y or I424T, -/+ CD74 ectopic expression. NK was quantified only in IRF8 WT and N87Y mutant. Gating strategy for T-regs was shown in Supplemental Figure 6A. Color labeling scheme is shown at the top right.

**Supplemental Figure 6G.**
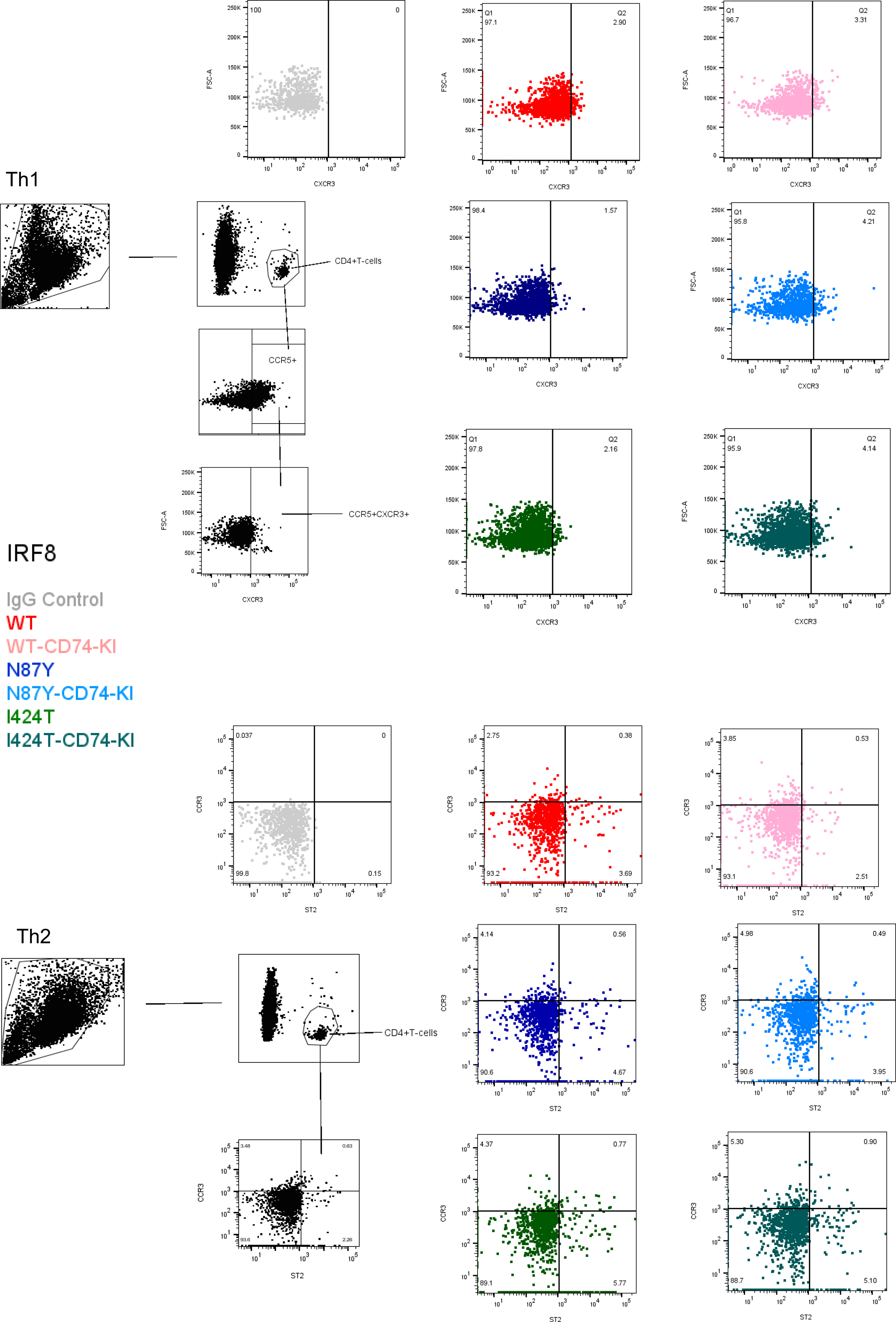
FACS analysis of lymphoma microenvironment. **Top to bottom.** Representative displays of Th1 and Th2 measurements in the TME of lymphomas expressing IRF8 WT, N87Y or I424T, -/+ CD74 ectopic expression. The gating strategies and color labeling scheme are shown to the left of the figure.

**Supplemental Figure 6H.**
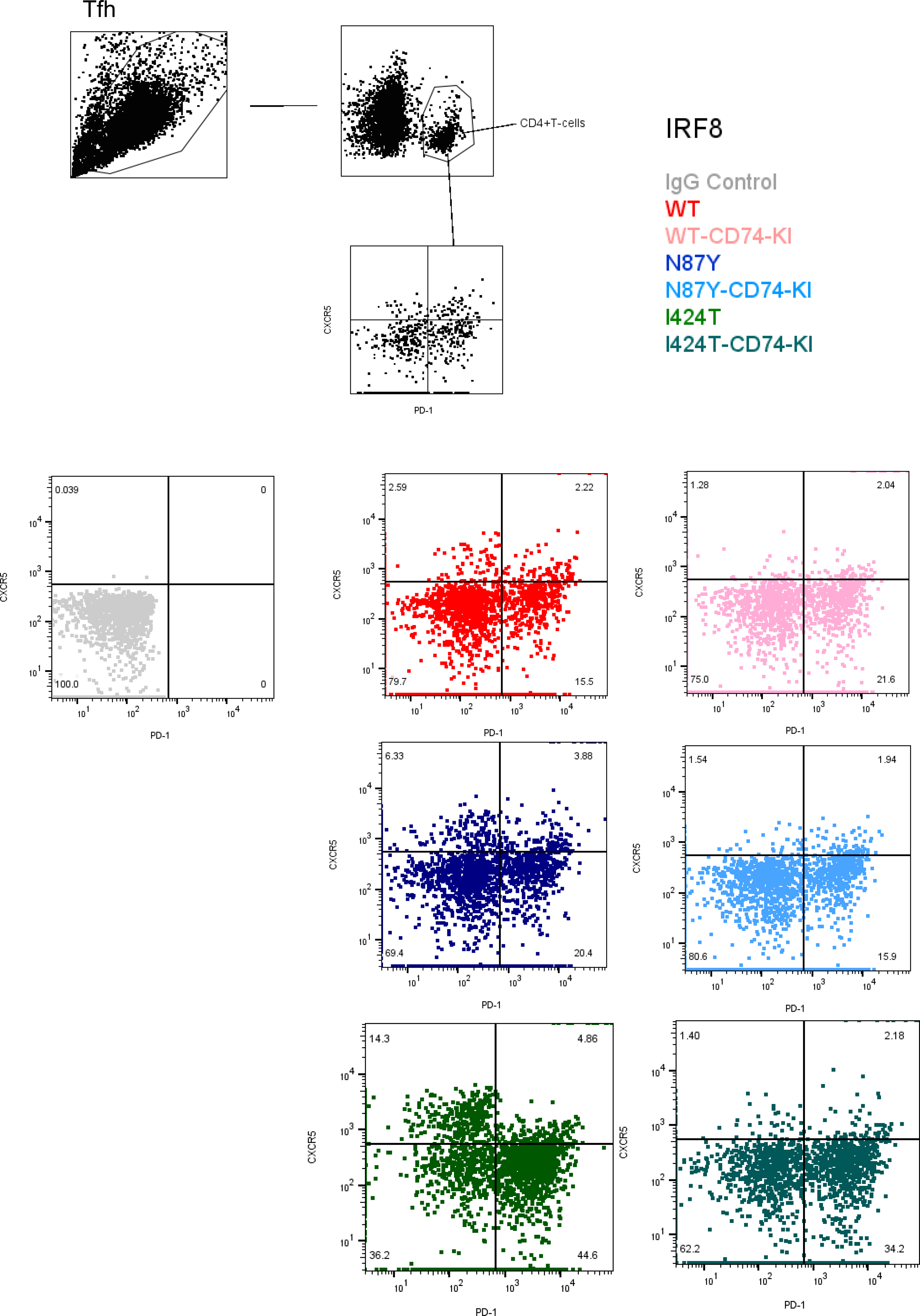
FACS analysis of lymphoma microenvironment. Representative displays of Tfh cell measurements in the TME of lymphomas expressing IRF8 WT, N87Y or I424T, -/+ CD74 ectopic expression. The gating strategies and color labeling scheme are shown at the top of the figure.

**Supplemental Figure 7A.**
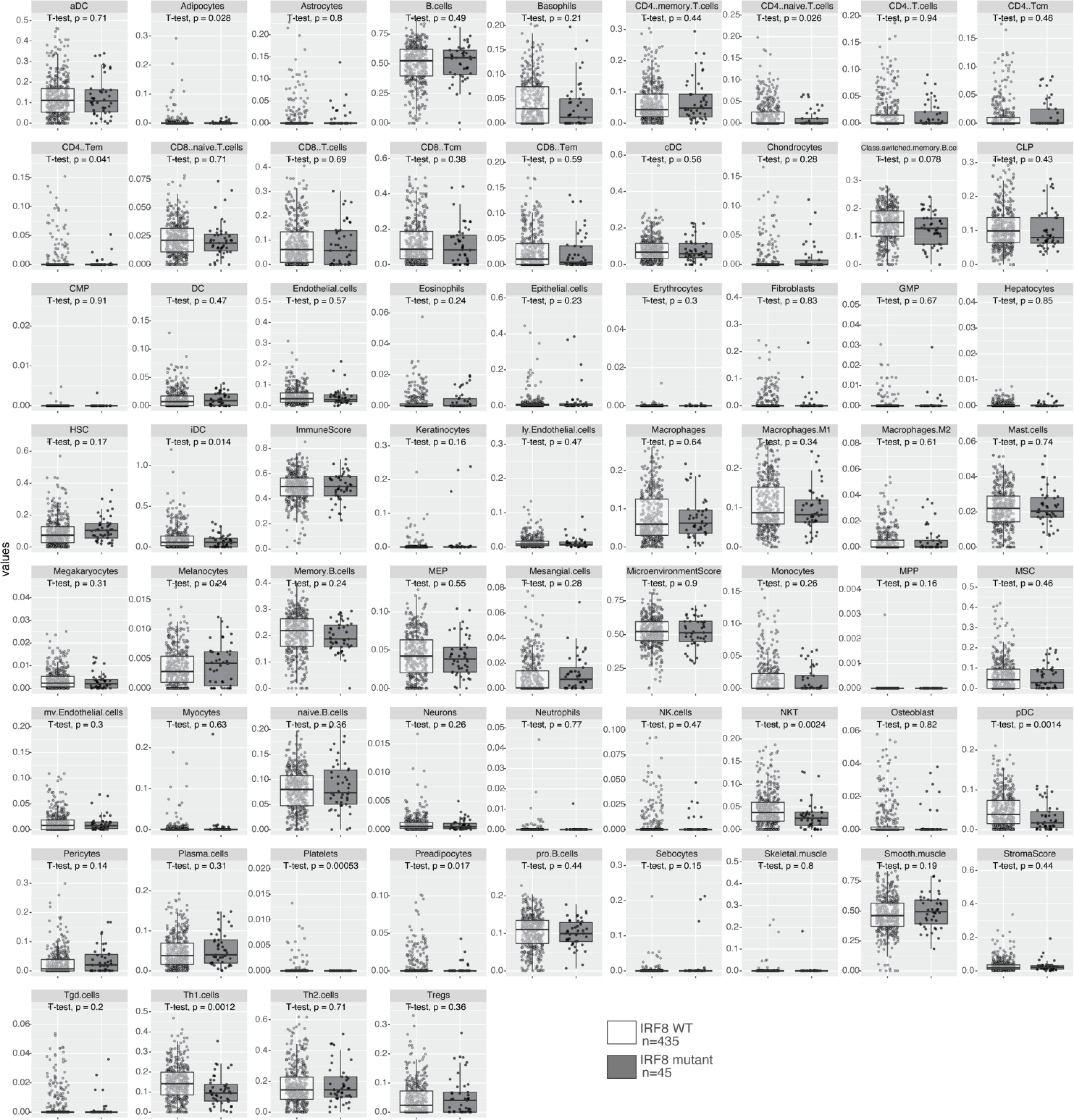
Immune composition of the lymphoma microenvironment of IRF8-WT and mutant human primary DLBCLs. Estimates of 64 cell type distributions scores from 480 DLBCL RNAseq (from dbGAPphs001444.v2.p1) using xCELL in IRF8 mutant (gray, n=45) or wildtype (WT, white, n=435) tumors, boxplots depict median and interquartile range, p-values are from two-sided Student’s t-tests. Related to Figure 7A.

**Supplemental Figure 7B.**
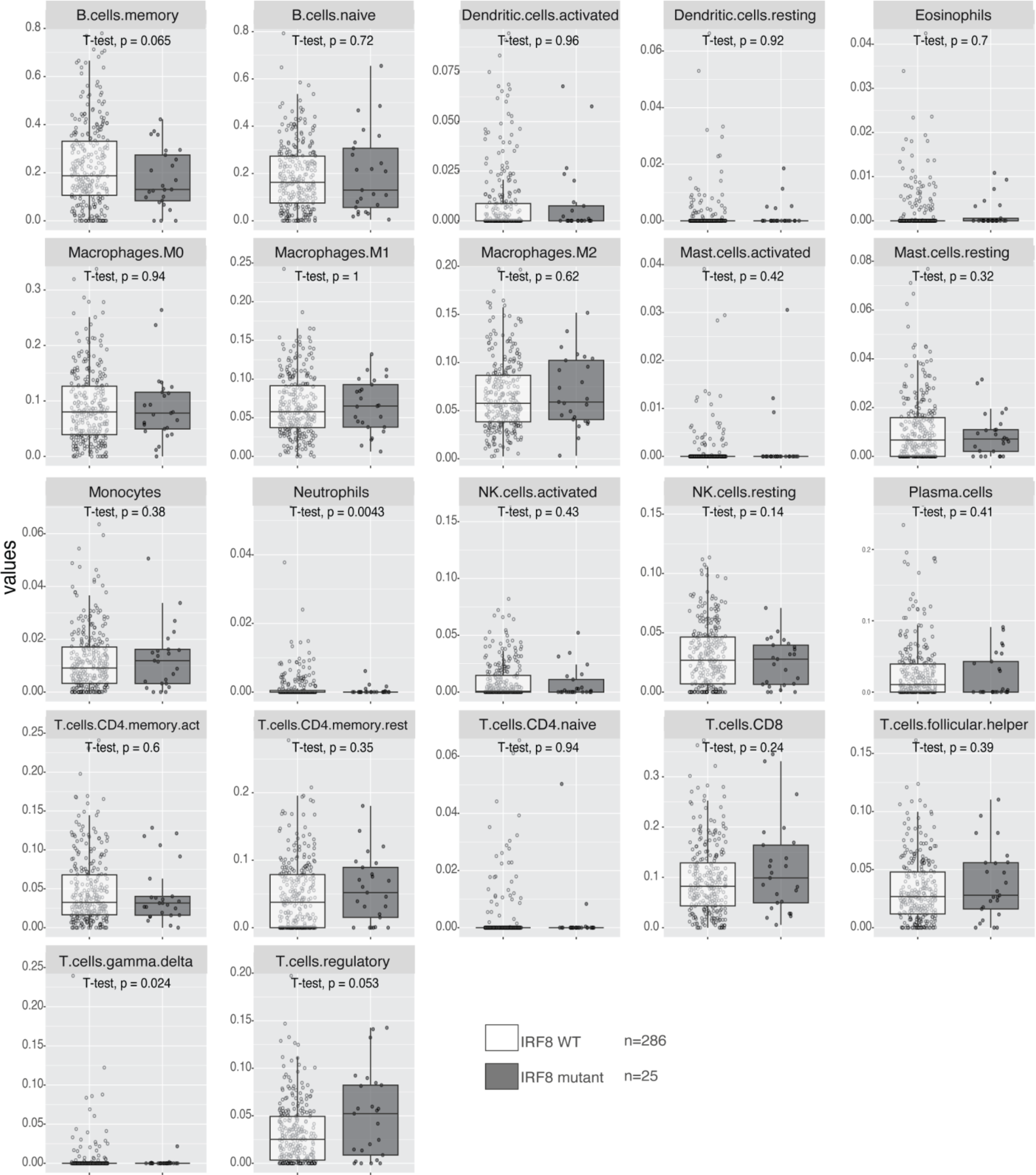
Immune composition of the lymphoma microenvironment of IRF8-WT and mutant human primary DLBCLs. CIBERSTORx-estimation of proportions of 22 immune cell types in a DLBLC sub-cohort (IRF8 WT, n = 286, IRF8 mutant, n = 25), that excludes tumors with mutation in other genes known to deregulate antigen processing/presentation (CIITA, B2M, CREBBP, EP300 and EZH2). Boxplots depict median and IQR. P values are from two-sided Student’s t-tests. Related to Figure 7B.

**Supplemental Figure 7C.**
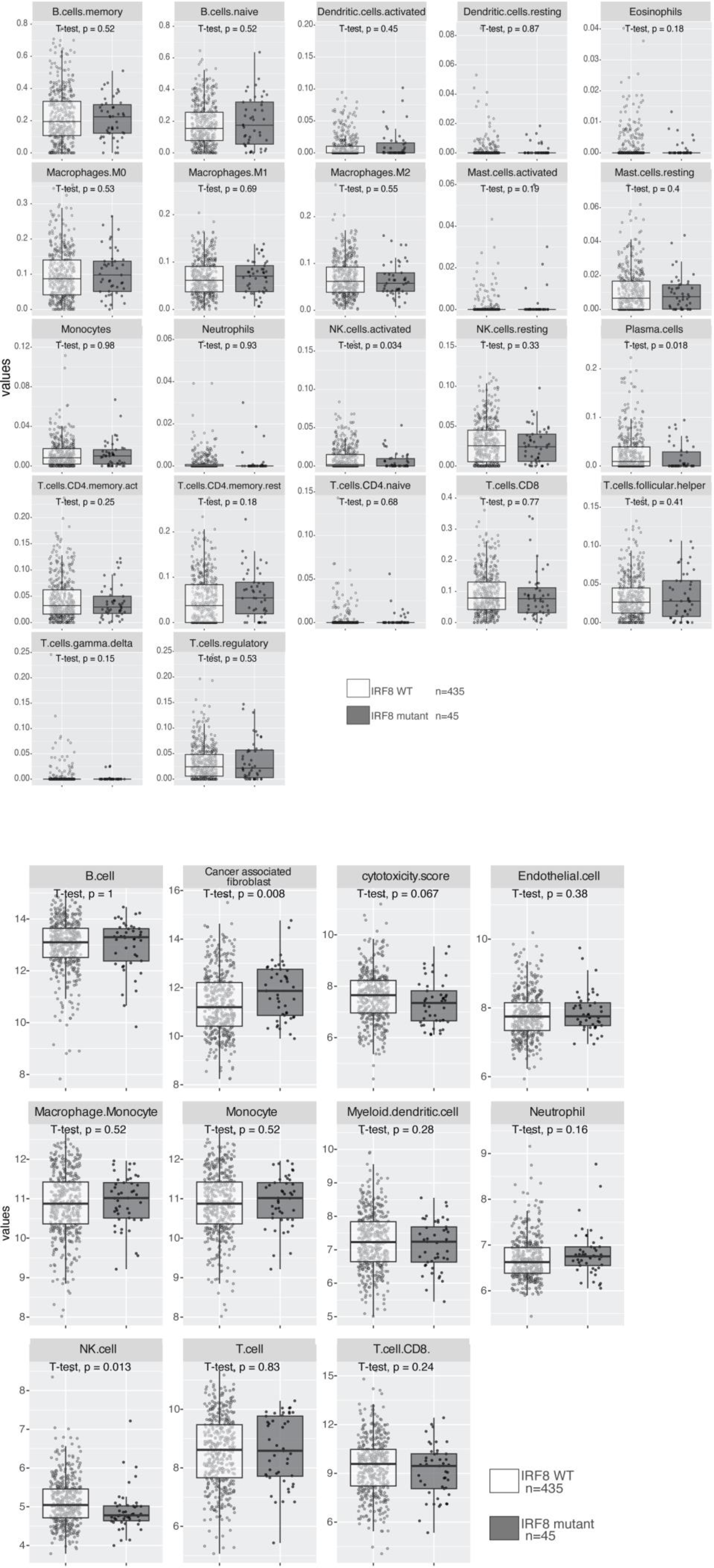
Immune composition of the lymphoma microenvironment of IRF8-WT and mutant human primary DLBCLs. **Top.** CIBERSTORx-estimation of proportions of 22 immune cell types in IRF8 mutant (gray, n=45) or wildtype (WT, white, n=435) DLBCL RNAseq (from dbGAPphs001444.v2.p1). Boxplots depict median and IQR. P values are from two-sided Students’ t-test. **Bottom.** Estimates of 11 immune cell types or scores distributions from 480 DLBCL RNAseq (from dbGAPphs001444.v2.p1) using MCP-counter in IRF8 mutant (gray, n=45) or wildtype (WT, white, n=435) tumors. Boxplots depict median and interquartile range, p-values are two-sided Student’s t-tests. Related to Figure 7C.

**Supplemental Figure 8A.**
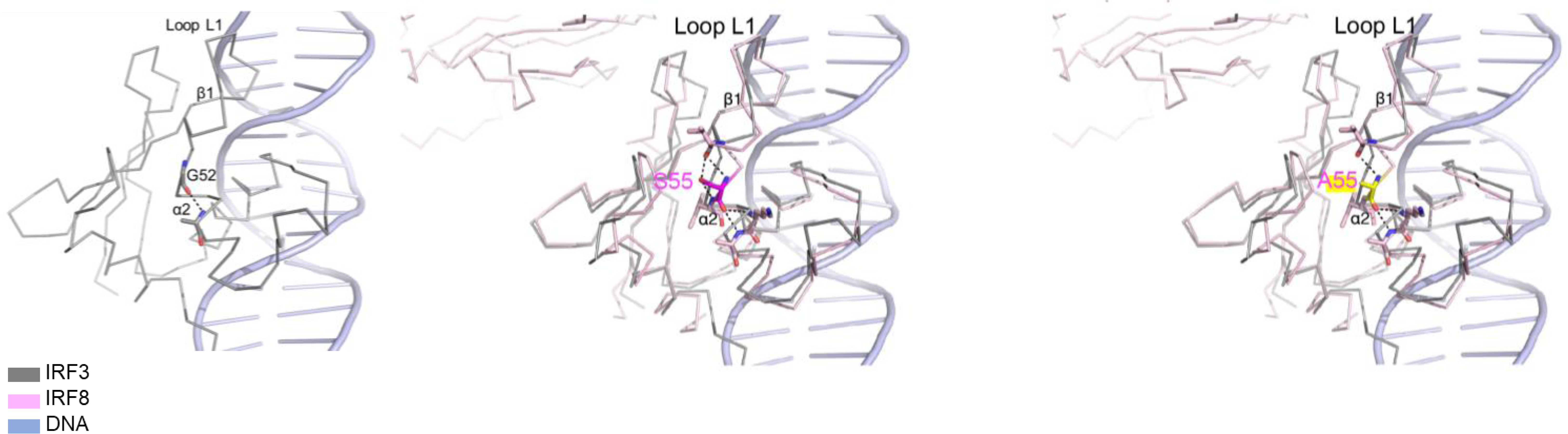
modeling of IRF8 mutation S55A. **Left**, solved structure of IRF3/DNA, displaying aa G52, equivalent to S55 in IRF8**. Middle,** overlay of the DNA binding domains of IRF3 (grey) and IRF8 (pink) with close-up in the S55 residue and its relation to the α2 helix of IRF8’s DBD and loop1 between α2 and β1 in contact with DNA. **Right,** A55 substitution, which may result in loss of the stabilizing interaction between the amino acid side chain and loop1.

**Supplemental Figure 8B.**
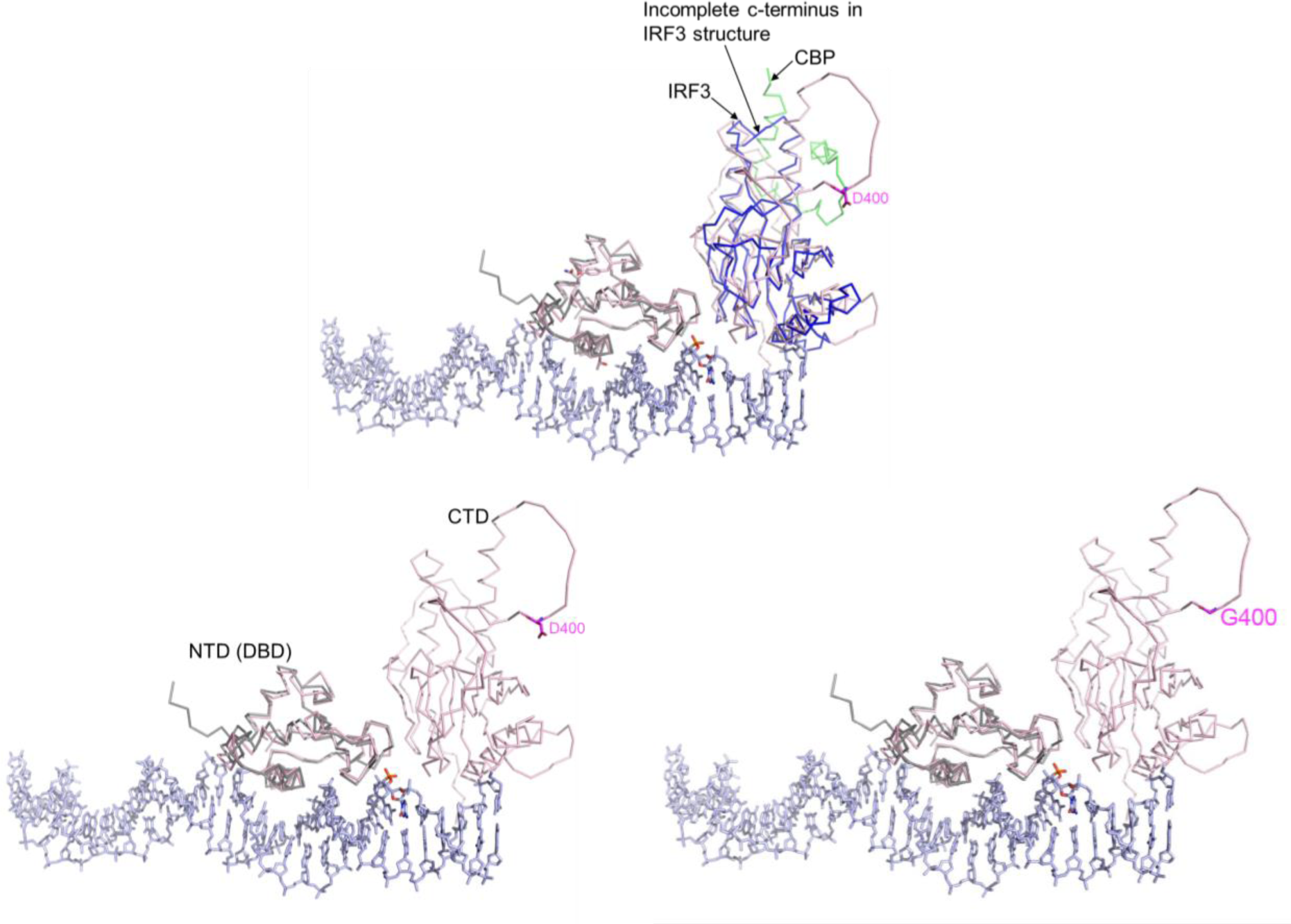
modeling of IRF8 mutation D400G. **Top** - overlay of the CTDs of IRF3 (blue ribbons) in IRF3 CTD/CBP complex (PDB: 1ZOQ) and IRF8 (pink). IRF3 CTD lacks the outer loop and c- terminal tail regions in the crystal structure. **Bottom** - an overlay of DBDs of IRF3 (PDB: 2PI0, grey, protein; blue, DNA) and IRF8 (AlphaFold model, pink) showing D400 (magenta stick) (**left**) or G400 (**right**) in an “outer loop”.

**Supplementary Table 1.** Summary of publicly available cohorts of Diffuse Large B-Cell Lymphomas (DLBCLs) reporting IRF8 mutations.

| Study Name | DLBCLs (n) | IRF8 mutant tumors (n) | % samples with IRF8 mutation | Source |
| --- | --- | --- | --- | --- |
| Diffuse Large B cell Lymphoma (DFCI, Nat Med 2018) | 135 | 9 | 6.7 | cBioportal and Chapuy et al. (2018) Nat. Med. 24, 679–690. |
| Diffuse Large B-cell Lymphoma (BCGSC, Blood 2013) | 30 | 3 | 10.0 | cBioportal and Morin et al. (2013) Blood 122 (7), 1256-65. |
| Diffuse Large B-Cell Lymphoma (TCGA, PanCancer Atlas) | 41 | 4 | 9.8 | cBioportal and Hoadley et al. (2018) Cell 173(2), 291-304. |
| Diffuse Large B-Cell Lymphoma (Duke, Cell 2017) | 1001 | 64 | 6.4 | cBioportal and Reddy et al. (2017) Cell 171(2), 481-494. |
| Mature B-cell malignancies (MD Anderson Cancer Center) | 148 | 8 | 5.4 | cBioPortal and Ma et al. (2022) Haematologica 107(3), 690-701. |
| Diffuse Large B-Cell Lymphoma (NCI, Staudt (NEJM, 2018, dbGAP) | 574 | 49 | 8.5 | dbGAP and Schmitz et al. (2018) N Engl J Med. 378(15),1396-1407. |
| Diffuse Large B-Cell Lymphoma (Lacy, UK, Blood, 2020) | 929 | 73 | 7.9 | Lacy et al. (2020) Blood, 135(20), 1759-71. |
| Diffuse Large B-Cell Lymphoma (LymphGen, Cancer Cell, 2020 and Ennishi et al, 2019a/b) | 329 | 34 | 10.3 | Wright et al. (2020) Cancer Cell 37, 551–568; Ennishi et al. (2019a) J. Clin. Oncol 37, 190–201; Ennishi et al (2019b) Cancer Discov. 9, 546–563. |
| Total | 3187 | 244 | 8.1 |  |

**Supplementary Table 2.**
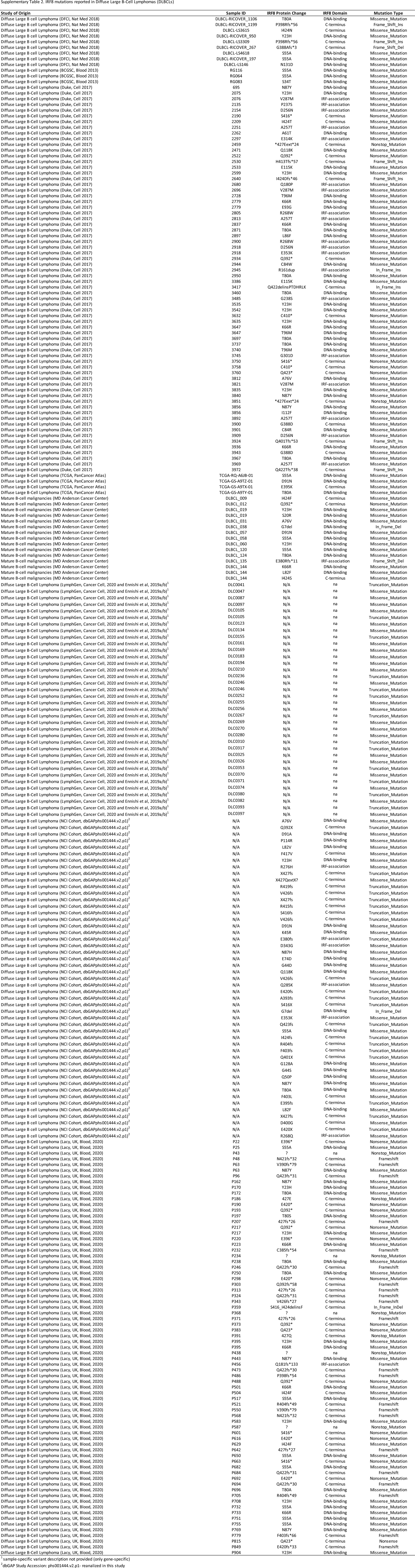
IRF8 mutations reported in Diffuse Large B-Cell Lymphomas (DLBCLs)

**Supplementary Table 3.** IRF8 variant allele frequency (VAF) % in DLBCLs.

| Study of Origin | Sample ID | IRF8 Protein Change | VAF ( %) | More than one |
| --- | --- | --- | --- | --- |
| Mature B-cell malignancies (MD Anderson Cancer Center) | DLBCL_009 | I424F | 37% |  |
| Mature B-cell malignancies (MD Anderson Cancer Center) | DLBCL_012 | Q392* | 15% |  |
| Mature B-cell malignancies (MD Anderson Cancer Center) | DLBCL_019 | Y23H | 38% | Yes |
| Mature B-cell malignancies (MD Anderson Cancer Center) | DLBCL_019 | S20R | 38% | Yes |
| Mature B-cell malignancies (MD Anderson Cancer Center) | DLBCL_031 | A76V | 35% |  |
| Mature B-cell malignancies (MD Anderson Cancer Center) | DLBCL_038 | G7del | 43% |  |
| Mature B-cell malignancies (MD Anderson Cancer Center) | DLBCL_057 | D91N | 42% |  |
| Mature B-cell malignancies (MD Anderson Cancer Center) | DLBCL_058 | S55A | 84% |  |
| Mature B-cell malignancies (MD Anderson Cancer Center) | DLBCL_060 | Y23H | 12% |  |
| Mature B-cell malignancies (MD Anderson Cancer Center) | DLBCL_120 | S55A | 23% |  |
| Mature B-cell malignancies (MD Anderson Cancer Center) | DLBCL_124 | T80A | 35% |  |
| Mature B-cell malignancies (MD Anderson Cancer Center) | DLBCL_135 | E380Rfs*11 | 28% |  |
| Mature B-cell malignancies (MD Anderson Cancer Center) | DLBCL_144 | K66R | 72% | Yes |
| Mature B-cell malignancies (MD Anderson Cancer Center) | DLBCL_144 | L82F | 73% | Yes |
| Mature B-cell malignancies (MD Anderson Cancer Center) | DLBCL_144 | I424S | 63% | Yes |
| Diffuse Large B cell Lymphoma (NCI Cohort, dbGAPphs001444.v2.p1*) | DLBCL10463 | Glu353Lys | 33% |  |
| Diffuse Large B cell Lymphoma (NCI Cohort, dbGAPphs001444.v2.p1*) | DLBCL10478 | Arg419fs | 68% |  |
| Diffuse Large B cell Lymphoma (NCI Cohort, dbGAPphs001444.v2.p1*) | DLBCL10488 | Glu395fs | 36% | Yes |
| Diffuse Large B cell Lymphoma (NCI Cohort, dbGAPphs001444.v2.p1*) | DLBCL10488 | Gln118Lys | 30% | Yes |
| Diffuse Large B cell Lymphoma (NCI Cohort, dbGAPphs001444.v2.p1*) | DLBCL10502 | Gln392* | 21% |  |
| Diffuse Large B cell Lymphoma (NCI Cohort, dbGAPphs001444.v2.p1*) | DLBCL10512 | Ter427Glnext*? | 1% |  |
| Diffuse Large B cell Lymphoma (NCI Cohort, dbGAPphs001444.v2.p1*) | DLBCL10516 | Asn421fs | 18% |  |
| Diffuse Large B cell Lymphoma (NCI Cohort, dbGAPphs001444.v2.p1*) | DLBCL10518 | Ter437fs | 49% | Yes |
| Diffuse Large B cell Lymphoma (NCI Cohort, dbGAPphs001444.v2.p1*) | DLBCL10518 | Val426fs | 49% | Yes |
| Diffuse Large B cell Lymphoma (NCI Cohort, dbGAPphs001444.v2.p1*) | DLBCL10519 | Tyr23His | 26% |  |
| Diffuse Large B cell Lymphoma (NCI Cohort, dbGAPphs001444.v2.p1*) | DLBCL10523 | Thr80Ala | 22% | Yes |
| Diffuse Large B cell Lymphoma (NCI Cohort, dbGAPphs001444.v2.p1*) | DLBCL10523 | Glu420* | 26% | Yes |
| Diffuse Large B cell Lymphoma (NCI Cohort, dbGAPphs001444.v2.p1*) | DLBCL10523 | Asn421fs | 26% | Yes |
| Diffuse Large B cell Lymphoma (NCI Cohort, dbGAPphs001444.v2.p1*) | DLBCL10527 | Thr80Ala | 13% | Yes |
| Diffuse Large B cell Lymphoma (NCI Cohort, dbGAPphs001444.v2.p1*) | DLBCL10527 | Leu82Phe | 13% | Yes |
| Diffuse Large B cell Lymphoma (NCI Cohort, dbGAPphs001444.v2.p1*) | DLBCL10527 | Glu74Asp | 16% | Yes |
| Diffuse Large B cell Lymphoma (NCI Cohort, dbGAPphs001444.v2.p1*) | DLBCL10532 | Thr80Ala | 39% |  |
| Diffuse Large B cell Lymphoma (NCI Cohort, dbGAPphs001444.v2.p1*) | DLBCL10553 | Asp91Asn | 43% |  |
| Diffuse Large B cell Lymphoma (NCI Cohort, dbGAPphs001444.v2.p1*) | DLBCL10554 | Glu380fs | 23% |  |
| Diffuse Large B cell Lymphoma (NCI Cohort, dbGAPphs001444.v2.p1*) | DLBCL10782 | Gly7del | 52% |  |
| Diffuse Large B cell Lymphoma (NCI Cohort, dbGAPphs001444.v2.p1*) | DLBCL10843 | Asn87Tyr | 41% | Yes |
| Diffuse Large B cell Lymphoma (NCI Cohort, dbGAPphs001444.v2.p1*) | DLBCL10843 | Asp91Ala | 42% | Yes |
| Diffuse Large B cell Lymphoma (NCI Cohort, dbGAPphs001444.v2.p1*) | DLBCL10894 | Arg415fs | 33% | Yes |
| Diffuse Large B cell Lymphoma (NCI Cohort, dbGAPphs001444.v2.p1*) | DLBCL10894 | Ser416del | 33% | Yes |
| Diffuse Large B cell Lymphoma (NCI Cohort, dbGAPphs001444.v2.p1*) | DLBCL10913 | Ser416del | 30% |  |
| Diffuse Large B cell Lymphoma (NCI Cohort, dbGAPphs001444.v2.p1*) | DLBCL10940 | Gln392* | 40% |  |
| Diffuse Large B cell Lymphoma (NCI Cohort, dbGAPphs001444.v2.p1*) | DLBCL10941 | Gln392* | 14% |  |
| Diffuse Large B cell Lymphoma (NCI Cohort, dbGAPphs001444.v2.p1*) | DLBCL10950 | Tyr23His | 30% |  |
| Diffuse Large B cell Lymphoma (NCI Cohort, dbGAPphs001444.v2.p1*) | DLBCL10965 | Gln392* | 22% |  |
| Diffuse Large B cell Lymphoma (NCI Cohort, dbGAPphs001444.v2.p1*) | DLBCL10993 | Asp400Gly | 25% |  |
| Diffuse Large B cell Lymphoma (NCI Cohort, dbGAPphs001444.v2.p1*) | DLBCL11181 | Asn87Tyr | 41% |  |
| Diffuse Large B cell Lymphoma (NCI Cohort, dbGAPphs001444.v2.p1*) | DLBCL11191 | Asn421fs | 11% |  |
| Diffuse Large B cell Lymphoma (NCI Cohort, dbGAPphs001444.v2.p1*) | DLBCL11430 | Ser55Ala | 23% |  |
| Diffuse Large B cell Lymphoma (NCI Cohort, dbGAPphs001444.v2.p1*) | DLBCL11477 | Phe403fs | 57% |  |
| Diffuse Large B cell Lymphoma (NCI Cohort, dbGAPphs001444.v2.p1*) | DLBCL11478 | Gln401* | 13% |  |
| Diffuse Large B cell Lymphoma (NCI Cohort, dbGAPphs001444.v2.p1*) | DLBCL11479 | Gln285Lys | 21% |  |
| Diffuse Large B cell Lymphoma (NCI Cohort, dbGAPphs001444.v2.p1*) | DLBCL11494 | Ter427fs | 11% |  |
| Diffuse Large B cell Lymphoma (NCI Cohort, dbGAPphs001444.v2.p1*) | DLBCL11497 | Ser55Ala | 43% |  |
| Diffuse Large B cell Lymphoma (NCI Cohort, dbGAPphs001444.v2.p1*) | DLBCL11507 | Thr80Ala | 47% |  |
| Diffuse Large B cell Lymphoma (NCI Cohort, dbGAPphs001444.v2.p1*) | DLBCL11519 | Gln118Lys | 35% |  |
| Diffuse Large B cell Lymphoma (NCI Cohort, dbGAPphs001444.v2.p1*) | DLBCL11526 | Glu395fs | 26% |  |
| Diffuse Large B cell Lymphoma (NCI Cohort, dbGAPphs001444.v2.p1*) | DLBCL11532 | Ter427Glnext*? | 45% |  |
| Diffuse Large B cell Lymphoma (NCI Cohort, dbGAPphs001444.v2.p1*) | DLBCL11541 | Val426fs | 36% |  |
| Diffuse Large B cell Lymphoma (NCI Cohort, dbGAPphs001444.v2.p1*) | DLBCL11542 | Gly44Ser | 32% | Yes |
| Diffuse Large B cell Lymphoma (NCI Cohort, dbGAPphs001444.v2.p1*) | DLBCL11542 | Gly44Asp | 32% | Yes |
| Diffuse Large B cell Lymphoma (NCI Cohort, dbGAPphs001444.v2.p1*) | DLBCL11542 | Ter427Glnext*? | 28% | Yes |
| Diffuse Large B cell Lymphoma (NCI Cohort, dbGAPphs001444.v2.p1*) | DLBCL11563 | Tyr23His | 12% |  |
| Diffuse Large B cell Lymphoma (NCI Cohort, dbGAPphs001444.v2.p1*) | DLBCL11568 | Leu82Val | 9% |  |
| Diffuse Large B cell Lymphoma (NCI Cohort, dbGAPphs001444.v2.p1*) | DLBCL11578 | Ser55Ala | 89% |  |
| Diffuse Large B cell Lymphoma (NCI Cohort, dbGAPphs001444.v2.p1*) | DLBCL11581 | Arg276His | 47% |  |
| Diffuse Large B cell Lymphoma (NCI Cohort, dbGAPphs001444.v2.p1*) | DLBCL11589 | Asn87Tyr | 23% |  |
| Diffuse Large B cell Lymphoma (NCI Cohort, dbGAPphs001444.v2.p1*) | DLBCL11590 | Tyr23His | 52% |  |
| Diffuse Large B cell Lymphoma (NCI Cohort, dbGAPphs001444.v2.p1*) | DLBCL11672 | Arg404fs | 79% |  |
| Diffuse Large B cell Lymphoma (NCI Cohort, dbGAPphs001444.v2.p1*) | DLBCL11675 | Pro114Arg | 59% |  |
| Diffuse Large B cell Lymphoma (NCI Cohort, dbGAPphs001444.v2.p1*) | DLBCL11680 | Ser55Ala | 38% |  |
| Diffuse Large B cell Lymphoma (DFCI, Nat Med 2018) | DLBCL-LS146 | N131D | 64% |  |
| Diffuse Large B cell Lymphoma (DFCI, Nat Med 2018) | DLBCL-LS3309 | P398Rfs*56 | 77% |  |
| Diffuse Large B cell Lymphoma (DFCI, Nat Med 2018) | DLBCL-LS3615 | I424N | 19% |  |
| Diffuse Large B cell Lymphoma (DFCI, Nat Med 2018) | DLBCL-LS4618 | S55A | 26% |  |
| Diffuse Large B cell Lymphoma (DFCI, Nat Med 2018) | DLBCL-RICOVER_1106 | T80A | 40% |  |
| Diffuse Large B cell Lymphoma (DFCI, Nat Med 2018) | DLBCL-RICOVER_1199 | P398Rfs*56 | 15% |  |
| Diffuse Large B cell Lymphoma (DFCI, Nat Med 2018) | DLBCL-RICOVER_197 | S55A | 56% |  |
| Diffuse Large B cell Lymphoma (DFCI, Nat Med 2018) | DLBCL-RICOVER_267 | G388Afs*3 | 52% |  |
| Diffuse Large B cell Lymphoma (DFCI, Nat Med 2018) | DLBCL-RICOVER_950 | Y23H | 21% |  |
| Diffuse Large B-Cell Lymphoma (TCGA, PanCancer Atlas) | TCGA-GS-A9TX-01 | E395K | 7% |  |
| Diffuse Large B-Cell Lymphoma (TCGA, PanCancer Atlas) | TCGA-GS-A9TY-01 | T80A | 62% |  |
| Diffuse Large B-Cell Lymphoma (TCGA, PanCancer Atlas) | TCGA-GS-A9TZ-01 | D91N | 36% |  |
| Diffuse Large B-Cell Lymphoma (TCGA, PanCancer Atlas) | TCGA-RQ-A6JB-01 | S55A | 78% |  |

**Supplementary Table 4.** Cell of origin (COO) classification of Diffuse Large B-Cell Lymphomas (DLBCLs) carrying IRF8 mutation from publicly available cohorts.

| Study of Origin | Sample ID | COO Class | Subgroup |
| --- | --- | --- | --- |
| Diffuse Large B cell Lymphoma (NCI Cohort, dbGAPphs001444.v2.p1) | DLBCL10463 | ABC | MCD |
| Diffuse Large B cell Lymphoma (NCI Cohort, dbGAPphs001444.v2.p1) | DLBCL10478 | GCB | ST2 |
| Diffuse Large B cell Lymphoma (NCI Cohort, dbGAPphs001444.v2.p1) | DLBCL10488 | Unclass | BN2 |
| Diffuse Large B cell Lymphoma (NCI Cohort, dbGAPphs001444.v2.p1) | DLBCL10502 | GCB | Other |
| Diffuse Large B cell Lymphoma (NCI Cohort, dbGAPphs001444.v2.p1) | DLBCL10512 | GCB | EZB |
| Diffuse Large B cell Lymphoma (NCI Cohort, dbGAPphs001444.v2.p1) | DLBCL10516 | ABC | Other |
| Diffuse Large B cell Lymphoma (NCI Cohort, dbGAPphs001444.v2.p1) | DLBCL10518 | GCB | EZB |
| Diffuse Large B cell Lymphoma (NCI Cohort, dbGAPphs001444.v2.p1) | DLBCL10519 | GCB | EZB |
| Diffuse Large B cell Lymphoma (NCI Cohort, dbGAPphs001444.v2.p1) | DLBCL10523 | GCB | EZB |
| Diffuse Large B cell Lymphoma (NCI Cohort, dbGAPphs001444.v2.p1) | DLBCL10527 | Unclass | Other |
| Diffuse Large B cell Lymphoma (NCI Cohort, dbGAPphs001444.v2.p1) | DLBCL10532 | ABC | MCD |
| Diffuse Large B cell Lymphoma (NCI Cohort, dbGAPphs001444.v2.p1) | DLBCL10553 | GCB | ST2 |
| Diffuse Large B cell Lymphoma (NCI Cohort, dbGAPphs001444.v2.p1) | DLBCL10554 | GCB | EZB |
| Diffuse Large B cell Lymphoma (NCI Cohort, dbGAPphs001444.v2.p1) | DLBCL10782 | GCB | Mixed |
| Diffuse Large B cell Lymphoma (NCI Cohort, dbGAPphs001444.v2.p1) | DLBCL10843 | ABC | BN2 |
| Diffuse Large B cell Lymphoma (NCI Cohort, dbGAPphs001444.v2.p1) | DLBCL10894 | ABC | Other |
| Diffuse Large B cell Lymphoma (NCI Cohort, dbGAPphs001444.v2.p1) | DLBCL10913 | ABC | BN2 |
| Diffuse Large B cell Lymphoma (NCI Cohort, dbGAPphs001444.v2.p1) | DLBCL10940 | ABC | MCD |
| Diffuse Large B cell Lymphoma (NCI Cohort, dbGAPphs001444.v2.p1) | DLBCL10941 | ABC | Mixed |
| Diffuse Large B cell Lymphoma (NCI Cohort, dbGAPphs001444.v2.p1) | DLBCL10950 | ABC | Other |
| Diffuse Large B cell Lymphoma (NCI Cohort, dbGAPphs001444.v2.p1) | DLBCL10965 | ABC | MCD |
| Diffuse Large B cell Lymphoma (NCI Cohort, dbGAPphs001444.v2.p1) | DLBCL10993 | ABC | BN2 |
| Diffuse Large B cell Lymphoma (NCI Cohort, dbGAPphs001444.v2.p1) | DLBCL11181 | Unclass | BN2 |
| Diffuse Large B cell Lymphoma (NCI Cohort, dbGAPphs001444.v2.p1) | DLBCL11191 | Unclass | BN2 |
| Diffuse Large B cell Lymphoma (NCI Cohort, dbGAPphs001444.v2.p1) | DLBCL11430 | ABC | Other |
| Diffuse Large B cell Lymphoma (NCI Cohort, dbGAPphs001444.v2.p1) | DLBCL11477 | Unclass | BN2 |
| Diffuse Large B cell Lymphoma (NCI Cohort, dbGAPphs001444.v2.p1) | DLBCL11478 | Unclass | BN2 |
| Diffuse Large B cell Lymphoma (NCI Cohort, dbGAPphs001444.v2.p1) | DLBCL11479 | Unclass | Other |
| Diffuse Large B cell Lymphoma (NCI Cohort, dbGAPphs001444.v2.p1) | DLBCL11494 | Unclass | BN2 |
| Diffuse Large B cell Lymphoma (NCI Cohort, dbGAPphs001444.v2.p1) | DLBCL11497 | ABC | EZB |
| Diffuse Large B cell Lymphoma (NCI Cohort, dbGAPphs001444.v2.p1) | DLBCL11507 | Unclass | A53 |
| Diffuse Large B cell Lymphoma (NCI Cohort, dbGAPphs001444.v2.p1) | DLBCL11519 | GCB | Other |
| Diffuse Large B cell Lymphoma (NCI Cohort, dbGAPphs001444.v2.p1) | DLBCL11526 | GCB | EZB |
| Diffuse Large B cell Lymphoma (NCI Cohort, dbGAPphs001444.v2.p1) | DLBCL11532 | GCB | EZB |
| Diffuse Large B cell Lymphoma (NCI Cohort, dbGAPphs001444.v2.p1) | DLBCL11541 | GCB | Other |
| Diffuse Large B cell Lymphoma (NCI Cohort, dbGAPphs001444.v2.p1) | DLBCL11542 | GCB | EZB |
| Diffuse Large B cell Lymphoma (NCI Cohort, dbGAPphs001444.v2.p1) | DLBCL11563 | GCB | EZB |
| Diffuse Large B cell Lymphoma (NCI Cohort, dbGAPphs001444.v2.p1) | DLBCL11568 | GCB | EZB |
| Diffuse Large B cell Lymphoma (NCI Cohort, dbGAPphs001444.v2.p1) | DLBCL11578 | GCB | Other |
| Diffuse Large B cell Lymphoma (NCI Cohort, dbGAPphs001444.v2.p1) | DLBCL11581 | GCB | EZB |
| Diffuse Large B cell Lymphoma (NCI Cohort, dbGAPphs001444.v2.p1) | DLBCL11589 | GCB | EZB |
| Diffuse Large B cell Lymphoma (NCI Cohort, dbGAPphs001444.v2.p1) | DLBCL11590 | GCB | EZB |
| Diffuse Large B cell Lymphoma (NCI Cohort, dbGAPphs001444.v2.p1) | DLBCL11672 | GCB | EZB |
| Diffuse Large B cell Lymphoma (NCI Cohort, dbGAPphs001444.v2.p1) | DLBCL11675 | GCB | EZB |
| Diffuse Large B cell Lymphoma (NCI Cohort, dbGAPphs001444.v2.p1) | DLBCL11680 | GCB | EZB |
| Diffuse Large B cell Lymphoma (DFCI, Nat Med 2018) | DLBCL-LS146 | GCB | A53 |
| Diffuse Large B cell Lymphoma (DFCI, Nat Med 2018) | DLBCL-LS3309 | GCB | EZB/A53 |
| Diffuse Large B cell Lymphoma (DFCI, Nat Med 2018) | DLBCL-LS3615 | Unclass | Other |
| Diffuse Large B cell Lymphoma (DFCI, Nat Med 2018) | DLBCL-LS4618 | GCB | EZB |
| Diffuse Large B cell Lymphoma (DFCI, Nat Med 2018) | DLBCL-RICOVER_1106 | ABC | MCD |
| Diffuse Large B cell Lymphoma (DFCI, Nat Med 2018) | DLBCL-RICOVER_197 | GCB | EZB |
| Diffuse Large B cell Lymphoma (DFCI, Nat Med 2018) | DLBCL-RICOVER_267 | GCB | EZB |
| BCC (Cancer Cell/LymphGen- (original Ennishi, J. Clin. Oncol ) | DLC0041 | GCB | EZB |
| BCC (Cancer Cell/LymphGen- (original Ennishi, J. Clin. Oncol ) | DLC0047 | GCB | EZB |
| BCC (Cancer Cell/LymphGen- (original Ennishi, J. Clin. Oncol ) | DLC0087 | GCB | EZB |
| BCC (Cancer Cell/LymphGen- (original Ennishi, J. Clin. Oncol ) | DLC0097 | GCB | EZB/ST2/A53 |
| BCC (Cancer Cell/LymphGen- (original Ennishi, J. Clin. Oncol ) | DLC0105 | Unclass | Other |
| BCC (Cancer Cell/LymphGen- (original Ennishi, J. Clin. Oncol ) | DLC0123 | Unclass | Other |
| BCC (Cancer Cell/LymphGen- (original Ennishi, J. Clin. Oncol ) | DLC0134 | GCB | EZB |
| BCC (Cancer Cell/LymphGen- (original Ennishi, J. Clin. Oncol ) | DLC0155 | GCB | EZB |
| BCC (Cancer Cell/LymphGen- (original Ennishi, J. Clin. Oncol ) | DLC0161 | GCB | Other |
| BCC (Cancer Cell/LymphGen- (original Ennishi, J. Clin. Oncol ) | DLC0169 | GCB | ST2 |
| BCC (Cancer Cell/LymphGen- (original Ennishi, J. Clin. Oncol ) | DLC0183 | Unclass | ST2 |
| BCC (Cancer Cell/LymphGen- (original Ennishi, J. Clin. Oncol ) | DLC0194 | GCB | EZB |
| BCC (Cancer Cell/LymphGen- (original Ennishi, J. Clin. Oncol ) | DLC0210 | ABC | N1 |
| BCC (Cancer Cell/LymphGen- (original Ennishi, J. Clin. Oncol ) | DLC0236 | GCB | EZB |
| BCC (Cancer Cell/LymphGen- (original Ennishi, J. Clin. Oncol ) | DLC0246 | GCB | BN2/MCD |
| BCC (Cancer Cell/LymphGen- (original Ennishi, J. Clin. Oncol ) | DLC0252 | GCB | EZB/A53 |
| BCC (Cancer Cell/LymphGen- (original Ennishi, J. Clin. Oncol ) | DLC0255 | GCB | Other |
| BCC (Cancer Cell/LymphGen- (original Ennishi, J. Clin. Oncol ) | DLC0256 | GCB | EZB |
| BCC (Cancer Cell/LymphGen- (original Ennishi, J. Clin. Oncol ) | DLC0267 | GCB | EZB/A53 |
| BCC (Cancer Cell/LymphGen- (original Ennishi, J. Clin. Oncol ) | DLC0269 | GCB | BN2 |
| BCC (Cancer Cell/LymphGen- (original Ennishi, J. Clin. Oncol ) | DLC0270 | GCB | EZB |
| BCC (Cancer Cell/LymphGen- (original Ennishi, J. Clin. Oncol ) | DLC0280 | GCB | EZB/ST2/A53 |
| BCC (Cancer Cell/LymphGen- (original Ennishi, J. Clin. Oncol ) | DLC0310 | GCB | BN2/EZB |
| BCC (Cancer Cell/LymphGen- (original Ennishi, J. Clin. Oncol ) | DLC0317 | ABC | Other |
| BCC (Cancer Cell/LymphGen- (original Ennishi, J. Clin. Oncol ) | DLC0325 | GCB | ST2 |
| BCC (Cancer Cell/LymphGen- (original Ennishi, J. Clin. Oncol ) | DLC0326 | GCB | EZB |
| BCC (Cancer Cell/LymphGen- (original Ennishi, J. Clin. Oncol ) | DLC0353 | GCB | EZB |
| BCC (Cancer Cell/LymphGen- (original Ennishi, J. Clin. Oncol ) | DLC0370 | ABC | EZB |
| BCC (Cancer Cell/LymphGen- (original Ennishi, J. Clin. Oncol ) | DLC0371 | GCB | EZB |
| BCC (Cancer Cell/LymphGen- (original Ennishi, J. Clin. Oncol ) | DLC0374 | GCB | EZB |
| BCC (Cancer Cell/LymphGen- (original Ennishi, J. Clin. Oncol ) | DLC0380 | GCB | EZB |
| BCC (Cancer Cell/LymphGen- (original Ennishi, J. Clin. Oncol ) | DLC0382 | GCB | EZB |
| BCC (Cancer Cell/LymphGen- (original Ennishi, J. Clin. Oncol ) | DLC0393 | GCB | EZB |
| BCC (Cancer Cell/LymphGen- (original Ennishi, J. Clin. Oncol ) | DLC0397 | Unclass | ST2/A53 |
| Diffuse Large B-Cell Lymphoma (TCGA, PanCancer Atlas) | TCGA-GS-A9TX | GCB | Other |
| Diffuse Large B-Cell Lymphoma (TCGA, PanCancer Atlas) | TCGA-GS-A9TZ | GCB | ST2 |
| Diffuse Large B-Cell Lymphoma (TCGA, PanCancer Atlas) | TCGA-RQ-A6JB | GCB | EZB |
| Diffuse Large B cell Lymphoma (DFCI, Nat Med 2018) | DLBCL-RICOVER_1199 | N/A | EZB |
| Diffuse Large B cell Lymphoma (DFCI, Nat Med 2018) | DLBCL-RICOVER_950 | N/A | Other |

**Supplementary Table 5.**
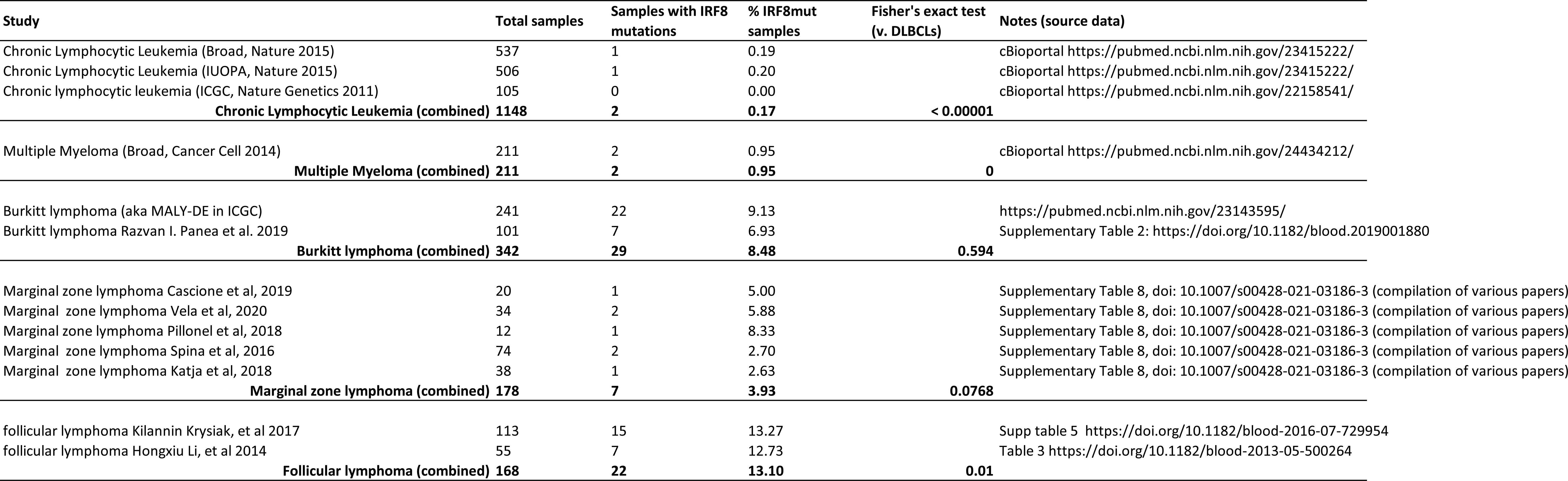
Summary of IRF8 mutation rates in published cohorts of other mature B-cell malignancies.

| Study | Total samples | Samples with IRF8 mutations | % IRF8mut samples | Fisher's exact test (v. DLBCLs) | Notes (source data) |
| --- | --- | --- | --- | --- | --- |
| Chronic Lymphocytic Leukemia (Broad, Nature 2015) | 537 | 1 | 0.19 |  | cBioportal https://pubmed.ncbi.nlm.nih.gov/23415222/ |
| Chronic Lymphocytic Leukemia (IUOPA, Nature 2015) | 506 | 1 | 0.20 |  | cBioportal https://pubmed.ncbi.nlm.nih.gov/23415222/ |
| Chronic lymphocytic leukemia (ICGC, Nature Genetics 2011) | 105 | 0 | 0.00 |  | cBioportal https://pubmed.ncbi.nlm.nih.gov/22158541/ |
| Chronic Lymphocytic Leukemia (combined) | 1148 | 2 | 0.17 | < 0.00001 |  |
| Multiple Myeloma (Broad, Cancer Cell 2014) | 211 | 2 | 0.95 |  | cBioportal https://pubmed.ncbi.nlm.nih.gov/24434212/ |
| Multiple Myeloma (combined) | 211 | 2 | 0.95 | 0 |  |
| Burkitt lymphoma (aka MALY-DE in ICGC) | 241 | 22 | 9.13 |  | https://pubmed.ncbi.nlm.nih.gov/23143595/ |
| Burkitt lymphoma Razvan I. Panea et al. 2019 | 101 | 7 | 6.93 |  | Supplementary Table 2: https://doi.org/10.1182/blood.2019001880 |
| Burkitt lymphoma (combined) | 342 | 29 | 8.48 | 0.594 |  |
| Marginal zone lymphoma Cascione et al, 2019 | 20 | 1 | 5.00 |  | Supplementary Table 8, doi: 10.1007/s00428-021-03186-3 (compilation of various papers) |
| Marginal zone lymphoma Vela et al, 2020 | 34 | 2 | 5.88 |  | Supplementary Table 8, doi: 10.1007/s00428-021-03186-3 (compilation of various papers) |
| Marginal zone lymphoma Pillonel et al, 2018 | 12 | 1 | 8.33 |  | Supplementary Table 8, doi: 10.1007/s00428-021-03186-3 (compilation of various papers) |
| Marginal zone lymphoma Spina et al, 2016 | 74 | 2 | 2.70 |  | Supplementary Table 8, doi: 10.1007/s00428-021-03186-3 (compilation of various papers) |
| Marginal zone lymphoma Katja et al, 2018 | 38 | 1 | 2.63 |  | Supplementary Table 8, doi: 10.1007/s00428-021-03186-3 (compilation of various papers) |
| Marginal zone lymphoma (combined) | 178 | 7 | 3.93 | 0.0768 |  |
| follicular lymphoma Kilannin Krysiak, et al 2017 | 113 | 15 | 13.27 |  | Supp table 5 https://doi.org/10.1182/blood-2016-07-729954 |
| follicular lymphoma Hongxiu Li, et al 2014 | 55 | 7 | 12.73 |  | Table 3 https://doi.org/10.1182/blood-2013-05-500264 |
| Follicular lymphoma (combined) | 168 | 22 | 13.10 | 0.01 |  |

**Supplementary Table 6.**
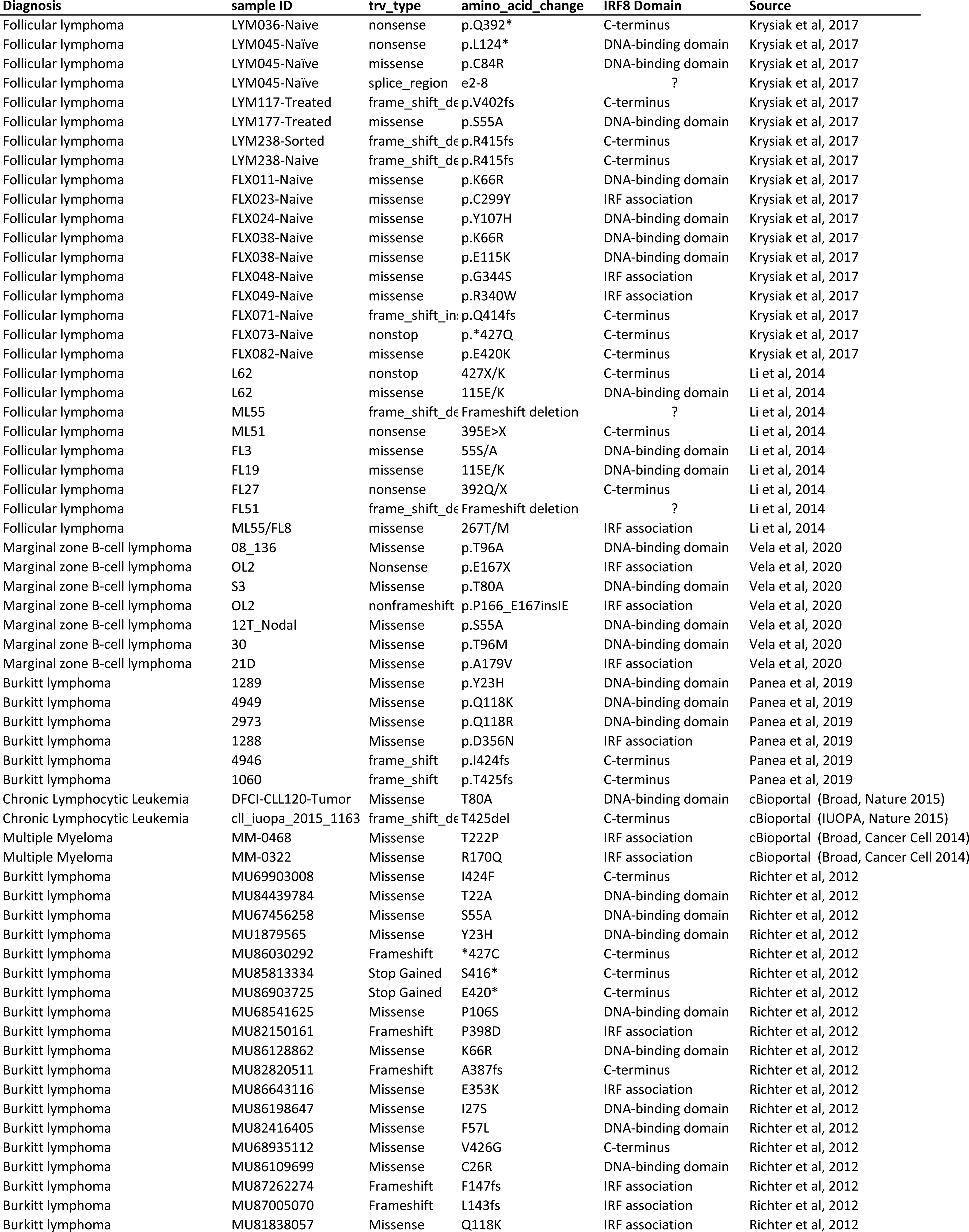
List of IRF8 mutations in published cohorts of other mature B-cell malignancies.

**Supplementary Table 7.**
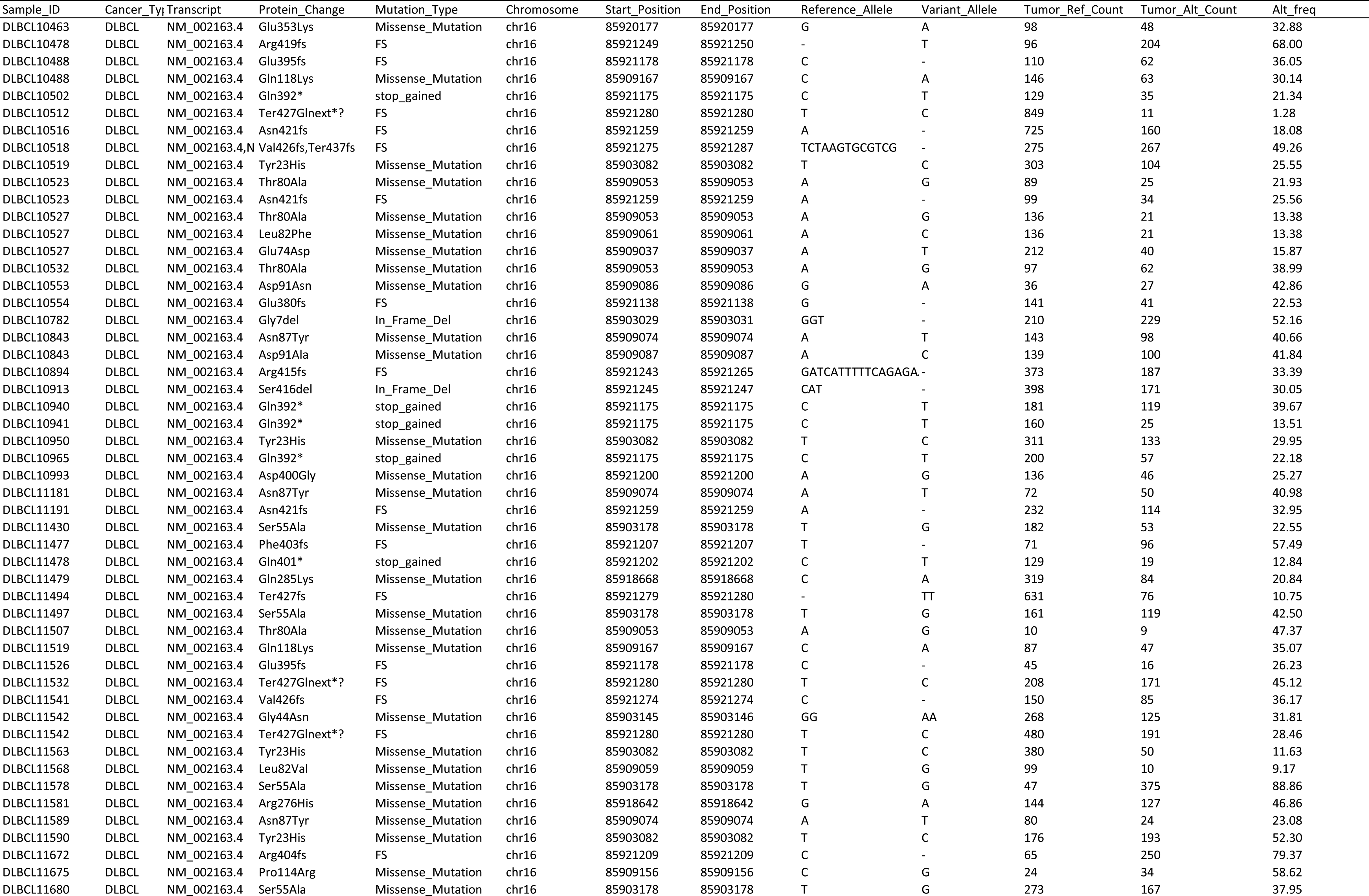
IRF8 mutations reported in Diffuse Large B-Cell Lymphomas (DLBCLs) from NCI cohort, reanalyzed from dbGAPphs001444.v2.p1.

**Supplemental Table 8.**
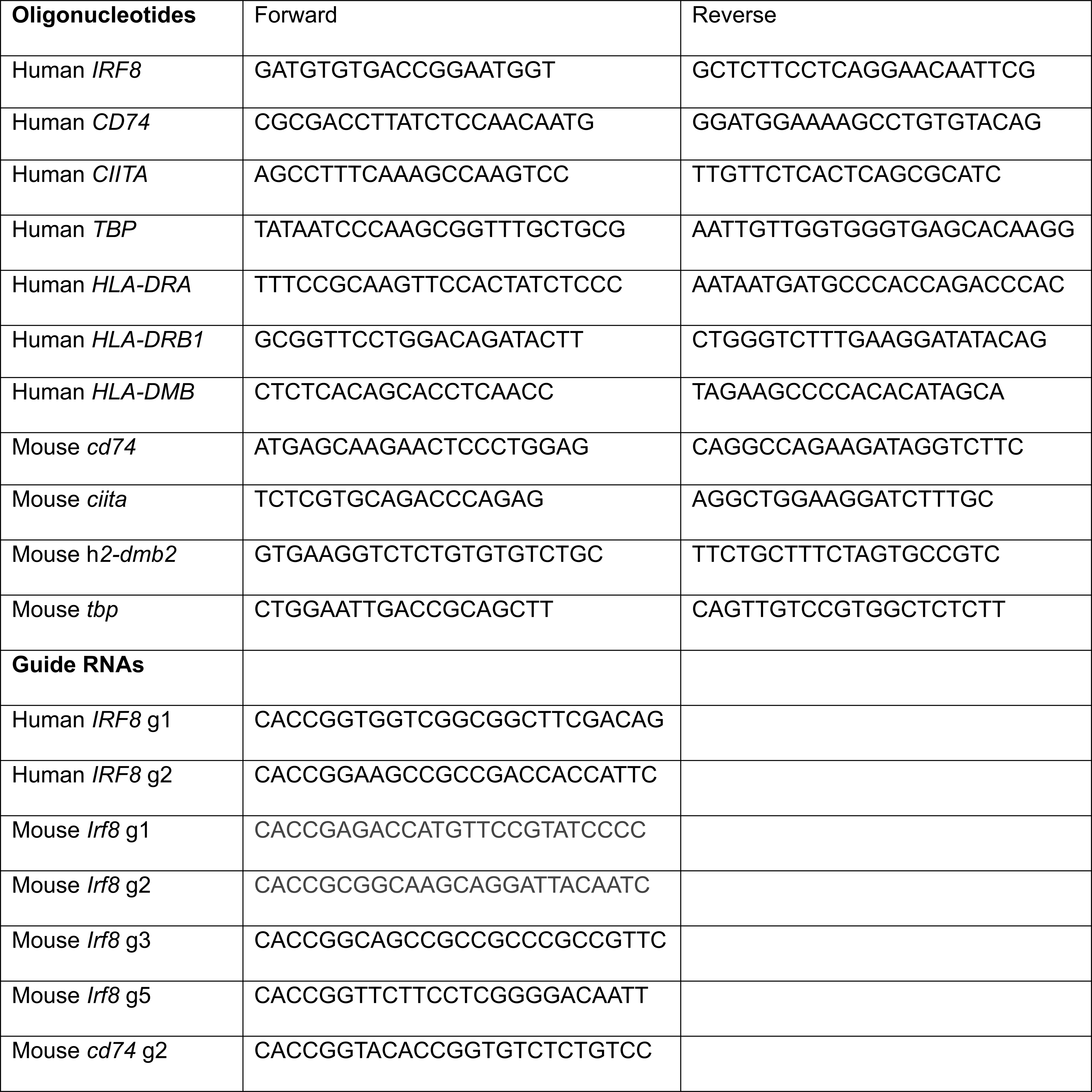

## References

1. Thorsson, V., et al. The Immune Landscape of Cancer. Immunity 48, 812–830 e814 (2018).

2. Wellenstein, M.D. & de Visser, K.E. Cancer-Cell-Intrinsic Mechanisms Shaping the Tumor Immune Landscape. Immunity 48, 399–416 (2018).

3. Sharma, P., et al. Immune checkpoint therapy-current perspectives and future directions. Cell 186, 1652–1669 (2023).

4. Challa-Malladi, M., et al. Combined genetic inactivation of beta2-Microglobulin and CD58 reveals frequent escape from immune recognition in diffuse large B cell lymphoma. Cancer Cell 20, 728–740 (2011).

5. Steidl, C., et al. MHC class II transactivator CIITA is a recurrent gene fusion partner in lymphoid cancers. Nature 471, 377–381 (2011).

6. Mottok, A., et al. Genomic Alterations in CIITA Are Frequent in Primary Mediastinal Large B Cell Lymphoma and Are Associated with Diminished MHC Class II Expression. Cell reports 13, 1418–1431 (2015).

7. Ennishi, D., et al. Molecular and Genetic Characterization of MHC Deficiency Identifies EZH2 as Therapeutic Target for Enhancing Immune Recognition. Cancer discovery 9, 546–563 (2019).

8. Hashwah, H., et al. Inactivation of CREBBP expands the germinal center B cell compartment, down-regulates MHCII expression and promotes DLBCL growth. Proc Natl Acad Sci U S A 114, 9701–9706 (2017).

9. Green, M.R., et al. Mutations in early follicular lymphoma progenitors are associated with suppressed antigen presentation. Proc Natl Acad Sci U S A 112, E1116–1125 (2015).

10. Morin, R.D., Arthur, S.E. & Hodson, D.J. Molecular profiling in diffuse large B-cell lymphoma: why so many types of subtypes? Br J Haematol 196, 814–829 (2022).

11. Ennishi, D., Hsi, E.D., Steidl, C. & Scott, D.W. Toward a New Molecular Taxonomy of Diffuse Large B-cell Lymphoma. Cancer discovery 10, 1267–1281 (2020).

12. Chapuy, B., et al. Molecular subtypes of diffuse large B cell lymphoma are associated with distinct pathogenic mechanisms and outcomes. Nat Med 24, 679–690 (2018).

13. Schmitz, R., et al. Genetics and Pathogenesis of Diffuse Large B-Cell Lymphoma. N Engl J Med 378, 1396–1407 (2018).

14. Pasqualucci, L. & Dalla-Favera, R. Genetics of diffuse large B-cell lymphoma. Blood 131, 2307–2319 (2018).

15. Bouamar, H., et al. A capture-sequencing strategy identifies IRF8, EBF1, and APRIL as novel IGH fusion partners in B-cell lymphoma. Blood 122, 726–733 (2013).

16. Chapuy, B., et al. Discovery and characterization of super-enhancer-associated dependencies in diffuse large B cell lymphoma. Cancer Cell 24, 777–790 (2013).

17. Bal, E., et al. Super-enhancer hypermutation alters oncogene expression in B cell lymphoma. Nature 607, 808–815 (2022).

18. Wang, H., et al. IRF8 regulates B-cell lineage specification, commitment, and differentiation. Blood 112, 4028–4038 (2008).

19. Yanez, A. & Goodridge, H.S. Interferon regulatory factor 8 and the regulation of neutrophil, monocyte, and dendritic cell production. Curr Opin Hematol 23, 11–17 (2016).

20. Holtschke, T., et al. Immunodeficiency and chronic myelogenous leukemia-like syndrome in mice with a targeted mutation of the ICSBP gene. Cell 87, 307–317 (1996).

21. Salem, S., et al. Functional characterization of the human dendritic cell immunodeficiency associated with the IRF8(K108E) mutation. Blood 124, 1894–1904 (2014).

22. Salem, S., Salem, D. & Gros, P. Role of IRF8 in immune cells functions, protection against infections, and susceptibility to inflammatory diseases. Hum Genet 139, 707–721 (2020).

23. Mace, E.M., et al. Biallelic mutations in IRF8 impair human NK cell maturation and function. The Journal of clinical investigation 127, 306–320 (2017).

24. Hambleton, S., et al. IRF8 mutations and human dendritic-cell immunodeficiency. N Engl J Med 365, 127–138 (2011).

25. Martinez, A., et al. Expression of the interferon regulatory factor 8/ICSBP-1 in human reactive lymphoid tissues and B-cell lymphomas: a novel germinal center marker. The American journal of surgical pathology 32, 1190–1200 (2008).

26. Lee, C.H., et al. Regulation of the germinal center gene program by interferon (IFN) regulatory factor 8/IFN consensus sequence-binding protein. J Exp Med 203, 63–72 (2006).

27. Feng, J., et al. IFN regulatory factor 8 restricts the size of the marginal zone and follicular B cell pools. J Immunol 186, 1458–1466 (2011).

28. Carotta, S., et al. The transcription factors IRF8 and PU.1 negatively regulate plasma cell differentiation. J Exp Med 211, 2169–2181 (2014).

29. Wang, H., et al. Transcription factors IRF8 and PU.1 are required for follicular B cell development and BCL6-driven germinal center responses. Proc Natl Acad Sci U S A 116, 9511–9520 (2019).

30. Tamura, T., Thotakura, P., Tanaka, T.S., Ko, M.S. & Ozato, K. Identification of target genes and a unique cis element regulated by IRF-8 in developing macrophages. Blood 106, 1938–1947 (2005).

31. Eagle, K., et al. Transcriptional Plasticity Drives Leukemia Immune Escape. Blood Cancer Discov 3, 394–409 (2022).

32. Marquis, J.F., et al. Interferon regulatory factor 8 regulates pathways for antigen presentation in myeloid cells and during tuberculosis. PLoS genetics 7, e1002097 (2011).

33. Shin, D.M., Lee, C.H. & Morse, H.C., 3rd. IRF8 governs expression of genes involved in innate and adaptive immunity in human and mouse germinal center B cells. PloS one 6, e27384 (2011).

34. Pishesha, N., Harmand, T.J. & Ploegh, H.L. A guide to antigen processing and presentation. Nat Rev Immunol 22, 751–764 (2022).

35. Devaiah, B.N. & Singer, D.S. CIITA and Its Dual Roles in MHC Gene Transcription. Front Immunol 4, 476 (2013).

36. Lacy, S.E., et al. Targeted sequencing in DLBCL, molecular subtypes, and outcomes: a Haematological Malignancy Research Network report. Blood 135, 1759–1771 (2020).

37. Reddy, A., et al. Genetic and Functional Drivers of Diffuse Large B Cell Lymphoma. Cell 171, 481–494 e415 (2017).

38. Ennishi, D., et al. Double-Hit Gene Expression Signature Defines a Distinct Subgroup of Germinal Center B-Cell-Like Diffuse Large B-Cell Lymphoma. J Clin Oncol 37, 190–201 (2019).

39. Morin, R.D., et al. Mutational and structural analysis of diffuse large B-cell lymphoma using whole-genome sequencing. Blood 122, 1256–1265 (2013).

40. Ma, M.C.J., et al. Subtype-specific and co-occurring genetic alterations in B-cell non-Hodgkin lymphoma. Haematologica 107, 690–701 (2022).

41. Wright, G.W., et al. A Probabilistic Classification Tool for Genetic Subtypes of Diffuse Large B Cell Lymphoma with Therapeutic Implications. Cancer Cell 37, 551–568 e514 (2020).

42. Krysiak, K., et al. Recurrent somatic mutations affecting B-cell receptor signaling pathway genes in follicular lymphoma. Blood 129, 473–483 (2017).

43. Li, H., et al. Mutations in linker histone genes HIST1H1 B, C, D, and E; OCT2 (POU2F2); IRF8; and ARID1A underlying the pathogenesis of follicular lymphoma. Blood 123, 1487–1498 (2014).

44. Panea, R.I., et al. The whole-genome landscape of Burkitt lymphoma subtypes. Blood 134, 1598–1607 (2019).

45. Richter, J., et al. Recurrent mutation of the ID3 gene in Burkitt lymphoma identified by integrated genome, exome and transcriptome sequencing. Nat Genet 44, 1316–1320 (2012).

46. Vela, V., Juskevicius, D., Dirnhofer, S., Menter, T. & Tzankov, A. Mutational landscape of marginal zone B- cell lymphomas of various origin: organotypic alterations and diagnostic potential for assignment of organ origin. Virchows Arch 480, 403–413 (2022).

47. Xu, Y., et al. Loss of IRF8 Inhibits the Growth of Diffuse Large B-cell Lymphoma. J Cancer 6, 953–961 (2015).

48. Smith, M.A., et al. Positive regulatory domain I (PRDM1) and IRF8/PU.1 counter-regulate MHC class II transactivator (CIITA) expression during dendritic cell maturation. J Biol Chem 286, 7893–7904 (2011).

49. Vila-del Sol, V., Punzon, C. & Fresno, M. IFN-gamma-induced TNF-alpha expression is regulated by interferon regulatory factors 1 and 8 in mouse macrophages. J Immunol 181, 4461–4470 (2008).

50. Liao, J., et al. Converting Lymphoma Cells into Potent Antigen-Presenting Cells for Interferon-Induced Tumor Regression. Cancer immunology research 5, 560–570 (2017).

51. Roy, B.M., Zhukov, D.V. & Maynard, J.A. Flanking residues are central to DO11.10 T cell hybridoma stimulation by ovalbumin 323-339. PloS one 7, e47585 (2012).

52. Miyagawa, F., Gutermuth, J., Zhang, H. & Katz, S.I. The use of mouse models to better understand mechanisms of autoimmunity and tolerance. J Autoimmun 35, 192–198 (2010).

53. de Charette, M. & Houot, R. Hide or defend, the two strategies of lymphoma immune evasion: potential implications for immunotherapy. Haematologica 103, 1256–1268 (2018).

54. Scharer, C.D., et al. Genome-wide CIITA-binding profile identifies sequence preferences that dictate function versus recruitment. Nucleic Acids Res 43, 3128–3142 (2015).

55. Nixon, B.G., et al. Tumor-associated macrophages expressing the transcription factor IRF8 promote T cell exhaustion in cancer. Immunity 55, 2044–2058 e2045 (2022).

56. Butler, M.J. & Aguiar, R.C.T. Biology Informs Treatment Choices in Diffuse Large B Cell Lymphoma. Trends in cancer 3, 871–882 (2017).

57. Aran, D., Hu, Z. & Butte, A.J. xCell: digitally portraying the tissue cellular heterogeneity landscape. Genome Biol 18, 220 (2017).

58. Krijgsman, D., Hokland, M. & Kuppen, P.J.K. The Role of Natural Killer T Cells in Cancer-A Phenotypical and Functional Approach. Front Immunol 9, 367 (2018).

59. Swiecki, M. & Colonna, M. The multifaceted biology of plasmacytoid dendritic cells. Nat Rev Immunol 15, 471–485 (2015).

60. Oliveira, G. & Wu, C.J. Dynamics and specificities of T cells in cancer immunotherapy. Nat Rev Cancer 23, 295–316 (2023).

61. Sworder, B.J., et al. Determinants of resistance to engineered T cell therapies targeting CD19 in large B cell lymphomas. Cancer Cell 41, 210–225 e215 (2023).

62. Newman, A.M., et al. Robust enumeration of cell subsets from tissue expression profiles. Nat Methods 12, 453–457 (2015).

63. Becht, E., et al. Estimating the population abundance of tissue-infiltrating immune and stromal cell populations using gene expression. Genome Biol 17, 218 (2016).

64. Escalante, C.R., Nistal-Villan, E., Shen, L., Garcia-Sastre, A. & Aggarwal, A.K. Structure of IRF-3 bound to the PRDIII-I regulatory element of the human interferon-beta enhancer. Mol Cell 26, 703–716 (2007).

65. Jumper, J., et al. Highly accurate protein structure prediction with AlphaFold. Nature 596, 583–589 (2021).

66. Levi, B.Z., Hashmueli, S., Gleit-Kielmanowicz, M., Azriel, A. & Meraro, D. ICSBP/IRF-8 transactivation: a tale of protein-protein interaction. J Interferon Cytokine Res 22, 153–160 (2002).

67. Adams, N.M., et al. Transcription Factor IRF8 Orchestrates the Adaptive Natural Killer Cell Response. Immunity 48, 1172–1182 e1176 (2018).

68. Sun, J.C., et al. Proinflammatory cytokine signaling required for the generation of natural killer cell memory. J Exp Med 209, 947–954 (2012).

69. Schiavoni, G., et al. ICSBP is essential for the development of mouse type I interferon-producing cells and for the generation and activation of CD8alpha(+) dendritic cells. J Exp Med 196, 1415–1425 (2002).

70. Shulman, Z., et al. T follicular helper cell dynamics in germinal centers. Science 341, 673–677 (2013).

71. Dheilly, E., et al. Cathepsin S Regulates Antigen Processing and T Cell Activity in Non-Hodgkin Lymphoma. Cancer Cell 37, 674–689 e612 (2020).

72. Boice, M., et al. Loss of the HVEM Tumor Suppressor in Lymphoma and Restoration by Modified CAR-T Cells. Cell 167, 405–418 e413 (2016).

73. Gutierrez-Melo, N. & Baumjohann, D. T follicular helper cells in cancer. Trends in cancer 9, 309–325 (2023).

74. Ortega, M., et al. A microRNA-mediated regulatory loop modulates NOTCH and MYC oncogenic signals in B- and T-cell malignancies. Leukemia 29, 968–976 (2015).

75. Lin, A.P., et al. MYC, mitochondrial metabolism and O-GlcNAcylation converge to modulate the activity and subcellular localization of DNA and RNA demethylases. Leukemia 36, 1150–1159 (2022).

76. Suhasini, A.N., et al. A phosphodiesterase 4B-dependent interplay between tumor cells and the microenvironment regulates angiogenesis in B-cell lymphoma. Leukemia 30, 617–626 (2016).

77. Lin, A.P., et al. D2HGDH regulates alpha-ketoglutarate levels and dioxygenase function by modulating IDH2. Nature communications 6, 7768 (2015).

78. Sasi, B., et al. Regulation of PD-L1 expression is a novel facet of cyclic-AMP-mediated immunosuppression. Leukemia 35, 1990–2001 (2021).

79. Cooney, J.D., et al. Synergistic Targeting of the Regulatory and Catalytic Subunits of PI3Kdelta in Mature B-cell Malignancies. Clinical cancer research: an official journal of the American Association for Cancer Research 24, 1103–1113 (2018).

80. Koboldt, D.C., et al. VarScan 2: somatic mutation and copy number alteration discovery in cancer by exome sequencing. Genome Res 22, 568–576 (2012).

81. Cerami, E., et al. The cBio cancer genomics portal: an open platform for exploring multidimensional cancer genomics data. Cancer discovery 2, 401–404 (2012).

82. Qiu, Z., et al. MYC Regulation of D2HGDH and L2HGDH Influences the Epigenome and Epitranscriptome. Cell Chem Biol 27, 538–550 e537 (2020).

83. Bouamar, H., et al. MicroRNA 155 Control of p53 Activity Is Context Dependent and Mediated by Aicda and Socs1. Mol Cell Biol 35, 1329–1340 (2015).

84. Bustos-Moran, E., et al. Aurora A controls CD8(+) T cell cytotoxic activity and antiviral response. Scientific reports 9, 2211 (2019).

85. Qiu, Z., et al. Generation and characterization of the Emicro-Irf8 mouse model. Cancer Genet 245, 6–16 (2020).

86. Ethiraj, P., et al. Cyclic-AMP signalling, MYC and hypoxia-inducible factor 1alpha intersect to regulate angiogenesis in B-cell lymphoma. Br J Haematol 198, 349–359 (2022).

87. Jiang, D. & Aguiar, R.C. MicroRNA-155 controls RB phosphorylation in normal and malignant B lymphocytes via the noncanonical TGF-beta1/SMAD5 signaling module. Blood 123, 86–93 (2014).

88. Emsley, P. & Cowtan, K. Coot: model-building tools for molecular graphics. Acta Crystallogr D Biol Crystallogr 60, 2126–2132 (2004).

